# Acclimation to enhanced photorespiratory conditions increases the effectiveness of C_2_ photosynthesis in C_3_-C_4_ intermediates

**DOI:** 10.64898/2026.09.22.753555

**Authors:** Arthur Leung, Martha Ludwig, Tammy L. Sage, Rowan F. Sage

## Abstract

**Summary:**

- C_2_ photosynthesis reduces photorespiratory carbon loss by shuttling glycine from mesophyll (M) to bundle sheath (BS) tissues, where glycine decarboxylase (GDC) releases CO_2_ that accumulates around rubisco. C_2_ photosynthesis has been considered a constitutive character, but as a metabolic trait, it may acclimate to the rate of photorespiratory glycine production.
- We grew eleven species from two eudicot and two grass lineages spanning C_3_ to C_2_ phenotypes at 32 °C days and 200 or 800 µmol mol^-1^ CO_2_. Acclimation was assessed using leaf gas exchange, GDC P-protein immunolocalization, M and BS ultrastructure, photosynthetic enzyme activity, and photosynthesis modelling.
- Low relative to high growth CO_2_ enhanced C_2_-related phenotypes in sub-C_2_ and C_2_ species but not C_3_ or proto-Kranz species. Sub-C_2_ species positioned chloroplasts closer to mitochondria and had lower modelled BS conductance. C_2_ species increased rubisco capacity and GDC allocation to the BS. Aspartate aminotransferase activity increased in both sub-C_2_ and C_2_ species.
- Acclimation to greater photorespiratory conditions strengthens C_2_ photosynthesis, but only in species whose BS has crossed a threshold of organelle and GDC investment. We hypothesize that acclimation could expose C_2_-related traits to selection and facilitate the origin of C_4_ photosynthesis.

## Introduction

Photorespiration is one of the largest metabolic fluxes in C_3_ leaves, leading to substantial declines in photosynthetic efficiency and capacity at elevated temperature and reduced atmospheric CO_2_ concentrations of recent geological time (Ehleringer *et al*., 1991; Walker *et al*., 2016). Under high temperature and low intercellular CO_2_, photorespiration can release over half of the previously fixed carbon through the activity of mitochondrial glycine decarboxylase (GDC) (Sharkey, 1988; Ehleringer *et al*., 1991). Close positioning of chloroplasts, mitochondria, and peroxisomes facilitates pathway flux and allows C_3_ leaves to recapture 10–20% of photorespired CO_2_ in their mesophyll cells (Brown *et al*., 1983b; Busch *et al*., 2013; Midorikawa *et al*., 2022).

Numerous plant lineages localize GDC expression in an inner leaf compartment such as the vascular sheath (often the parenchymatous bundle sheath, BS), which elevates the CO_2_ concentration in the sheath cells (Rawsthorne *et al*., 1988b; Hylton *et al*., 1988; Keerberg *et al*., 2014). In such species, photorespiratory glycine is shuttled from mesophyll (M) to BS tissues, which after glycine decarboxylation enhances the carboxylation efficiency of rubisco in the BS chloroplasts (Keerberg *et al*., 2014). The glycine shuttle and the associated alterations to photorespiratory metabolism that concentrate CO_2_ in the BS are together termed C_2_ photosynthesis (Vogan *et al*., 2007). Phylogenetic and physiological studies indicate that C_2_ photosynthesis precedes and facilitates the evolution of C_4_ photosynthesis, in part by establishing key biochemical and Kranz anatomical characteristics for the C_4_ cycle and putting in place rapid metabolite trafficking systems (Monson & Rawsthorne, 2000; Bauwe, 2010; Sage *et al*., 2012). Investigating the evolutionary assembly of C_2_ photosynthesis is therefore a central approach to understanding how and why C_4_ photosynthesis evolved (Sage, 2017).

Ancestral C_3_ lineages that gave rise to C_2_ photosynthesis are hypothesized to express a series of leaf phenotypes that progressively strengthen the capacity of BS tissues to recapture photorespiratory CO_2_ (Monson & Rawsthorne, 2000; Sage *et al*., 2012), in a pattern that may reflect anatomical and physiological potentiation (*sensu* Blount *et al*., 2008; see also Christin *et al*., 2013 and Griffiths *et al*., 2013). Potentiation, also termed activation, of the BS involves producing more chloroplasts and mitochondria that enable the BS to better metabolize glycine overflow from the M tissues (Hylton *et al*., 1988; Gowik & Westhoff, 2011; Sage *et al*., 2012).

Potentiation is argued to be present in ‘proto-Kranz’ and in certain C_3_ species which express a derived ‘C_3_+’ character state proposed to enhance refixation of photorespired CO_2_ in BS cells and by doing so may represent an early step in C_2_ and C_4_ evolution (Gowik & Westhoff, 2011; Muhaidat *et al*., 2011). In proto-Kranz species examined to date, BS cells have more mitochondria which are positioned centripetally against the inner cell wall, where the mitochondria presumably metabolize photorespiratory glycine and lengthen the diffusion path of released CO_2_ back to M cells (Brown *et al*., 1983a; Muhaidat *et al*., 2011; Christin *et al*., 2013; Sage *et al*., 2013). Whether conditions that promote photorespiration can induce the activated C_3_+ and proto-Kranz character states remains untested, and would reveal whether the character states are largely properties of genotype.

A character state representing further potentiation is the ‘sub-C_2_’ phenotype, in which mitochondrial GDC expression is enhanced in BS cells, yielding a modest reduction in the photosynthetic CO_2_ compensation point (Γ) at high temperature (Adachi *et al*., 2023; Leung *et al*., 2024). Sub-C_2_ taxa have greater numbers of organelles and greater GDC expression in the BS relative to surrounding M cells (Sage *et al*., 2013; Khoshravesh *et al*., 2020; Leung *et al*., 2024; Alvarenga *et al*., 2025). Relative to full C_2_ taxa, M cells in sub-C_2_ species show considerable capacity to decarboxylate glycine, as indicated by retention of GDC subunits in the M and Γ values intermediate between C_3_ and C_2_ values at warm and hot temperatures (Khoshravesh *et al*., 2020; Leung *et al*., 2024). By contrast, full C_2_ species restrict nearly all GDC subunits to the BS tissues and operate an efficient M-to-BS glycine shuttle, indicated by substantially lower Γ values than in C_3_ relatives (Morgan *et al*., 1993; Leung *et al*., 2024). The C_3_, proto-Kranz, sub-C_2_, and C_2_ phenotypes are suggested to be constitutive characters that are exhibited in distinct species, but whether the strength of the glycine shuttle within an individual differs with the photorespiratory conditions under which the leaf develops is uncertain. If acclimation can strengthen the glycine shuttle and move a sub-C_2_ phenotype towards a full C_2_ phenotype, then the traits described across the C_3_ to C_2_ continuum reflect both genotype and growth environment, potentially accelerating trait diversification and the evolutionary rise of C_2_ photosynthesis.

The ancestors of C_4_ plants likely occurred in warm and salinized and/or seasonally dry habitats where they experienced large fluctuations in photorespiration (Monson & Jaeger, 1991; Lundgren & Christin, 2017; Sage *et al*., 2018). Leaf temperature in such habitats often exceeds 40 °C while soil salinity and lapses between precipitation reduce stomatal conductance and thus CO_2_ diffusion into the M chloroplasts (Schuster & Monson, 1990; Monson & Jaeger, 1991).

Periods of high ribulose bisphosphate (RuBP) oxygenation can challenge the leaf’s ability to recover carbon from the phosphoglycolate pool; rate limitations within the photorespiratory pathway can feed back negatively onto CO_2_ assimilation or RuBP regeneration, slowing overall rates of photosynthesis (Eisenhut *et al*., 2017; López Calcagno *et al*., 2019; Timm *et al*., 2025). Activation of the BS tissues boosts photorespiratory capacity in the leaf, while also confining a fraction of leaf glycine decarboxylation to an internal compartment. CO_2_ and ammonia released in the BS can then be trapped and refixed effectively because the BS cell wall and numerous organelle membranes create a long diffusive path to slow their efflux (von Caemmerer & Furbank, 2003; Sage *et al*., 2014). In the proto-Kranz and sub-C_2_ phenotypes identified thus far, increasing GDC allocation to the BS is correlated with decreasing CO_2_ compensation point in the absence of day respiration (*C_i_*\*), supporting the hypothesis that increased trapping of photorespiratory CO_2_ is a major feature, or even a potentiator, of C_2_ evolution (Bauwe, 2010; Leung *et al*., 2024).

We hypothesize that leaves of C_3_ to C_2_ plants may acclimate to elevated photorespiratory potential in two ways. First, the rate of RuBP oxygenation could be lowered by deactivating rubisco, by limiting RuBP production, or by making less rubisco (Sharkey, 2005). Second, the photorespiratory capacity could be enhanced, particularly in a manner that more effectively traps and refixes photorespired CO_2_ and ammonia. Glycine and serine are key regulatory metabolites that modulate photorespiratory acclimation by regulating transcription of photorespiratory proteins such as GDC (Timm *et al*., 2013), and possibly, components of M-to- BS glycine shuttling. In C_3_ plants, acclimation of photorespiration is evident when plants are transferred from current atmospheres to high O_2_ or low CO_2_, which promote rubisco oxygenation relative to carboxylation. Values of Γ decreased in *Flaveria pringlei* and tobacco, and levels of transcripts encoding photorespiratory proteins increased in *Arabidopsis thaliana* (Byrd & Brown, 1989; Campbell *et al*., 2005; Li *et al*., 2014). In species with enhanced BS metabolic capacity, we hypothesize that overflow of photorespiratory metabolites into the BS tissues during elevated photorespiration triggers greater GDC expression in BS tissues through regulatory networks that already exist in C_3_ plants (Timm *et al*., 2013; Li *et al*., 2014).

To date, evidence for photorespiratory acclimation in C_3_-C_2_ intermediate and C_2_ phenotypes is limited, such that it remains uncertain whether increased glycine overflow can strengthen C_2_ character states. Lower Γ, consistent with strengthened C_2_ photosynthesis, has been reported in C_2_ species grown under conditions that raise photorespiration; for example, in *Flaveria* species grown at 21% or 40% versus 2% O_2_, 100 versus 400 µmol mol^-1^ CO_2_, and 40 °C versus 25 °C (Byrd & Brown, 1989; Teese, 1995; Wang *et al*., 2026); in *Moricandia arvensis* grown at 35 °C versus 20 °C (Gómez *et al*., 2020); and in *Chenopodium album* grown at 30 °C versus 20 °C under low soil nitrogen (Oono *et al*., 2022). Other studies showed inconsistent responses of Γ to growth temperature (in *Mollugo verticillata* populations; Hereford, 2017) or detected little or no acclimation under low versus current levels of growth CO_2_ (Vogan & Sage, 2012; Lundgren *et al*., 2019). In most cases the underlying mechanisms were not addressed (but see Oono *et al*., 2022). Interpretation is further complicated because Γ reflects respiration and rubisco carboxylation efficiency in addition to photorespiratory CO_2_ recapture (Brooks & Farquhar, 1985; von Caemmerer, 2000).

Here, we examined eleven species spanning the continuum from C_3_ to C_2_ photosynthesis, encompassing C_3_, proto-Kranz, sub-C_2_, and C_2_ phenotypes across two eudicot and two grass lineages. Plants were grown at 32 °C days and either 200 or 800 µmol mol^-1^ CO_2_ to manipulate photorespiration within geochemically realistic atmospheric conditions (CenCO_2_PIP Consortium *et al*., 2023). We assessed leaf gas exchange, GDC P-protein (GLDP) immunolocalization, M and vascular sheath ultrastructure, and C_4_ cycle enzyme activity. We also modelled steady-state photosynthetic parameters to evaluate BS physiology (von Caemmerer, 1989). Values of Γ and *C_i_*\* index leaf-level rubisco oxygenation relative to carboxylation. The number and positioning of BS chloroplasts and mitochondria, together with modelled conductance to CO_2_ efflux from the BS (*g*_bs_), index the potential to trap photorespired CO_2_ and ammonia. Allocation of GDC to the BS evaluates the strength of the glycine shuttle, while alanine and aspartate aminotransferase activities measure upregulation of key C_4_ cycle enzymes that support photorespiratory nitrogen fluxes (Mallmann *et al*., 2014). We addressed three specific questions: (i) Does low relative to high growth CO_2_ induce changes in CO_2_ compensation points, organelle positioning, GLDP localization, and aminotransferase activity? (ii) Does the magnitude of acclimation to low versus high CO_2_ depend on the degree of vascular sheath anatomical potentiation, and does its direction correlate with that of evolutionary change from C_3_ to C_2_ photosynthesis?

## Materials and Methods

### Plant material and experimental design

Eleven species of C_3_ to C_2_ phenotypes were studied: Flaveria cronquistii (C_3_), F. pringlei (C_3_), F. sonorensis (sub-C_2_), F. angustifolia (sub-C_2_), Tribulus forrestii (proto-Kranz), T. cristatus (C_2_), Homolepis glutinosa (C_3_), H. isocalycia (sub-C_2_), H. aturensis (C_2_), Steinchisma laxum (proto-Kranz), and S. hians (C_2_). The source information is given in prior work (Steinchisma, Khoshravesh et al., 2016; Flaveria, Adachi et al., 2023; Tribulus, Leung et al., 2024; Homolepis, Alvarenga et al., 2025), except for S. laxum which was obtained from the USDA Germplasm Resources Information Network (GRIN; PI 496379). Flaveria and Tribulus are eudicot genera and thus their vascular sheaths only possess single layer of parenchyma cells (Edwards & Voznesenskaya, 2010). In grasses, the inner sheath is commonly comprised of the mestome sheath, while the outer layer is the parenchymatous BS (Edwards & Voznesenskaya, 2010). In the grass genera Homolepis and Steinchisma, the outer parenchymatous layer is used for photosynthetic carbon reduction (Khoshravesh et al., 2016; Alvarenga et al., 2025). Because all four lineages here use the parenchymatous BS for photosynthetic carbon reduction, we focus primarily on BS physiology in this paper. Additionally, Flaveria pringlei has been described as proto-Kranz (Sage et al., 2013), but in the present accession its BS cells contained few organelles and little GLDP, placing it within the C_3_ category. Plants were propagated from cuttings (Flaveria, Homolepis) or seeds (Tribulus, Steinchisma), grown in 9-liter pots in a sandy-loam soil (Leung et al., 2024), watered every other day, and fertilized weekly (propagation, soil, fertilization, and CO_2_-control details are given in Methods S1).

Each species was represented by four plants, split evenly across two growth chamber experiments. Plants were initially grown at ambient CO_2_ in a greenhouse for at least four weeks, then transferred to growth CO_2_ concentrations of 200 or 800 µmol mol^-1^ and acclimated for at least 10 days before measurement. Plants then grew under a 13 h photoperiod at 550 to 750 µmol m^-2^ s^-1^ photosynthetic photon flux density (PPFD) and 32/22 °C day/night in GC-20 growth cabinets (BioChambers). Replicate treatments occurred in distinct GC-20 chambers. All reported measurements were made on leaves that matured during the 10-day acclimation period, except in the reversibility experiment, in which leaves had matured at 800 µmol mol^-1^ CO_2_ before transfer.

### Leaf gas exchange

Gas exchange was measured with two LI-6400XT systems fitted with 6400-02B LED light sources (Li-Cor Biosciences) at a leaf temperature of 30 °C following protocols from Leung *et al*. (2024). The CO_2_ compensation point (Γ) was determined at 1500 µmol m^-2^ s^-1^ PPFD from responses of net CO_2_ assimilation rate (*A*) to intercellular CO_2_ concentration (*C_i_*). The CO_2_ compensation point in the absence of day respiration (*C_i_*\*) and day respiration rate (*R_d_*) were determined by the Laisk method from *A*/*C_i_*responses measured at four lower PPFDs. The *C_i_* at the common intersection of *A*/*C_i_* responses approximates *C_i_*\*, and the negative of the corresponding *A* equals *R_d_* (Fig. 1; Fig. S1; Laisk, 1977; Brooks & Farquhar, 1985; Walker & Ort, 2015). The common intersection point was determined using the slope-intercept regression method of Walker & Ort (2015) (Fig. S2). Measurement CO_2_ sequences are given in Methods S2. In C_3_ species, *C_i_*\* approximates the chloroplastic CO_2_ compensation point in the absence of day respiration (Γ*). In C_2_ species, *C_i_*\* is used as a metric reflecting C_2_ photosynthetic capacity; the high-light *A*/*C_i_*response shows a shift to lower *C_i_* values, reducing Γ, whereas the low-light *A*/*C_i_*responses converge to enable determination of *C_i_*\* and *R_d_*(Fig. 1; Sage *et al*., 2013). We also report Γ values at an irradiance of 250 µmol m^-2^ s^-1^ (Γ_250_) as a metric of properties of the low-light *A*/*C_i_* responses used to calculate *C_i_*\*, such as CO_2_ refixation efficiency.

**Fig. 1.**
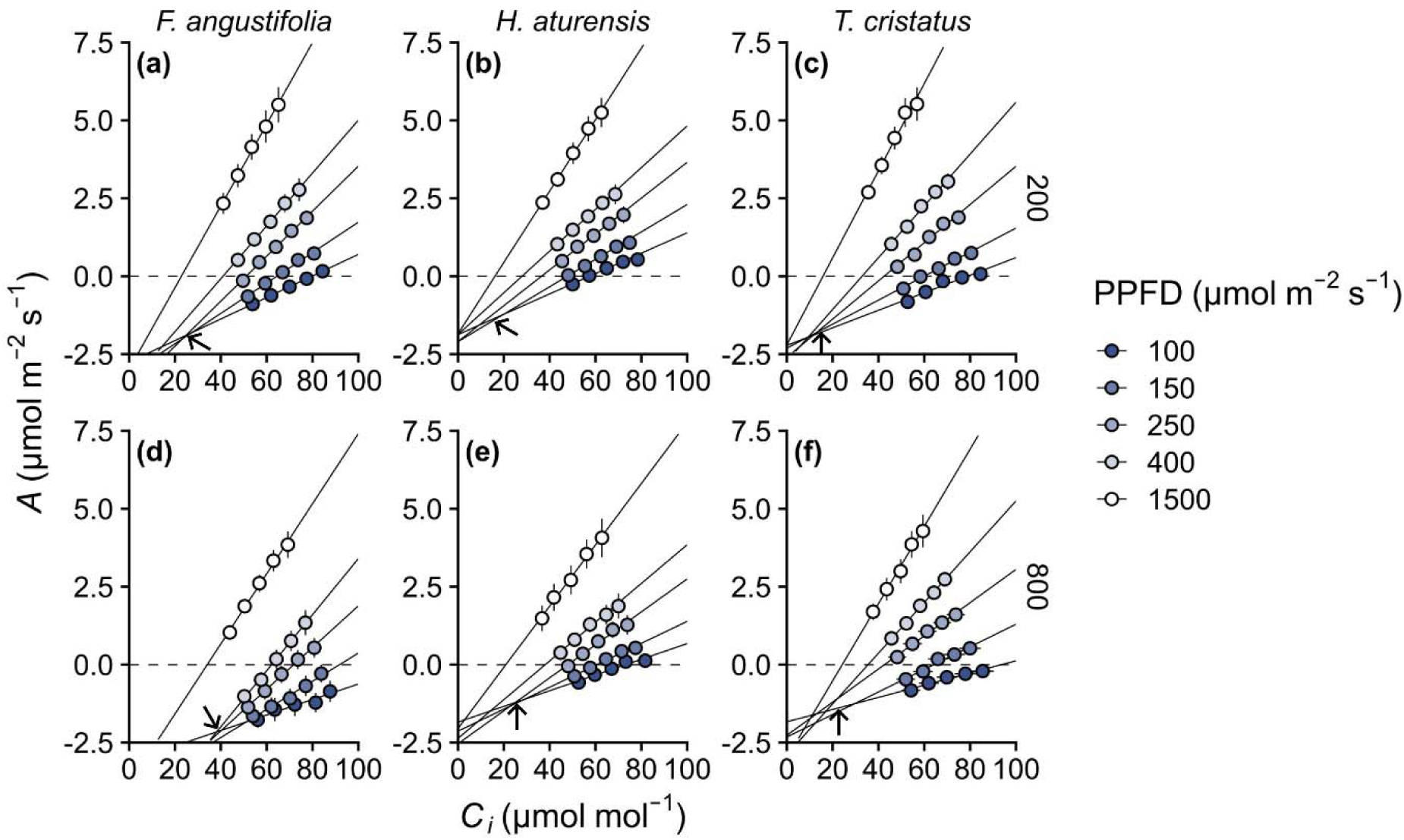
Gas exchange plasticity in response to growth CO_2_. (**a–f**), Response of net CO_2_ -assimilation (*A*) to intercellular CO_2_ concentration (*C_i_*) at 30 °C and five irradiances (100, 150,-250, 400, and 1500 µmol m^-2^ s^-1^) in three representative species from independent C_2_ lineages -(means ± SE; *n* = 4 plants). (**a–c**) Upper panels show plants grown at 200 µmol mol^-1^ CO_2_; (**d–f**) -lower panels show plants grown at 800 µmol mol^-1^ CO_2_. (**a,d**) *Flaveria angustifolia* (sub-C_2_), -(**b,e**) *Homolepis aturensis* (C_2_), and (**c,f**) *Tribulus cristatus* (C_2_). Lines are linear regressions at -each irradiance; arrow, their common intersection approximates the CO_2_ compensation point in the absence of day respiration (*C_i_*\*) (Brooks & Farquhar, 1985; Walker & Ort, 2015). For remaining species, see Fig. S1. For slope-intercept regressions used to determine the intersection point of *A*/*C_i_* responses, see Fig. S2.

### Enzyme activity

Leaves were collected 2 to 4 h into the photoperiod, frozen in liquid N_2_, and stored at -80 °C. Total leaf-level activities of phosphoenolpyruvate carboxylase (PEPC), NADP-malic enzyme (NADP-ME), NAD-malic enzyme (NAD-ME), aspartate aminotransferase (Asp-AT), and alanine aminotransferase (Ala-AT) were assayed at 30 °C by coupling NADPH or NADH oxidation or reduction to absorbance at 340 nm as described previously (Ashton *et al*., 1990; Adachi *et al*., 2023; Leung *et al*., 2024). Extraction and assay conditions are given in Methods S3.

### Light microscopy, transmission electron microscopy, and immunohistochemistry

Leaf tissue for microscopy was sampled from the middle of the youngest fully expanded leaf of each plant, between the midrib and the margin, 2 to 4 h into the photoperiod. Samples were prepared for light microscopy and transmission electron microscopy (TEM) to assess anatomy and ultrastructure as previously described (Khoshravesh *et al*., 2017, 2020; Leung *et al*., 2024). For light microscopy, 1.5 µm thick cross-sections were stained with toluidine blue and imaged on an Axioplan microscope (Zeiss; https://www.zeiss.com/) fitted with a DP71 camera (Olympus; https://www.olympus-global.com/). For immunolocalization, tissue from the same leaf region was fixed in 1% (v/v) glutaraldehyde and 1% (w/v) paraformaldehyde in 0.1 M sodium cacodylate buffer, pH 7.4, dehydrated in a graded ethanol series, and embedded in LR White.

Immunohistochemical detection of GLDP on 90 nm-thick sections used an anti-GLDP primary antibody and an 18 nm colloidal gold-conjugated secondary antibody. Sections were imaged at 80 kV on an HT7700 TEM (Hitachi; https://www.hitachi-hightech.com/). Antibody sources, dilutions, and blocking and wash steps are given in Methods S4.

### Quantification of structural features in light and transmission electron micrographs

For each plant, organelle and immunolabelling parameters were quantified using ImageJ v1.53 (Schneider *et al*., 2012) from five BS and five M cells. Cell values were averaged within a leaf to give one value per biological replicate. Chloroplast, mitochondria, and cell areas in planar cross-sections were traced to obtain organelle coverage (planar organelle area per cell area) and, from light micrographs, the BS and M fraction of chlorenchyma area. GLDP labelling density was measured as the number of colloidal gold particles per mitochondrial area and extrapolated to the BS allocation of leaf GLDP (*f*_GLDP_) from labelling density, mitochondrial coverage, and tissue fraction (Wang *et al*., 2017; Khoshravesh *et al*., 2020; Leung *et al*., 2024). Detailed procedures for quantifying leaf organelle and GLDP allocation are presented in Methods S5.

Chloroplast-to-mitochondrion and mitochondrion-to-chloroplast nearest neighbour edge distances and BS vacuole shape were also quantified. Both distances measure how tightly chloroplasts and mitochondria are packed against each other, but from opposite starting points. Measuring from each mitochondrion to its nearest chloroplast assesses whether photorespired CO_2_ is released next to a chloroplast. Measuring from each chloroplast to its nearest mitochondrion indexes the proportion of the chloroplasts positioned against a mitochondrion.

The nearest neighbour distance procedure is presented in Methods S6 and visualized in Figure S6.

Vacuole circularity describes the outline of the central vacuole, which is bounded by the surrounding cytoplasm, chloroplasts, and mitochondria. Organelles that protrude into the vacuolar space indent the tonoplast and lower circularity, whereas organelles arranged in an even layer against the cell wall leave a smoother vacuole outline. Circularity was measured to index how evenly chloroplasts and mitochondria are distributed around the cell periphery.

Vacuole parameter quantification procedures are presented in Methods S5.

### Modelling C_2_ photosynthesis

A steady-state model of C_2_ photosynthesis (von Caemmerer, 1989, 2000) was used to partition processes underlying measured *A*/*C_i_*responses into BS and M components and to estimate the maximal rubisco carboxylation rate (V*_c_*_max_) and BS conductance to CO_2_ (*g*_bs_). The model was empirically parameterized from ultrastructure: the BS allocation of GLDP was assumed to be the fraction of leaf photorespiratory glycine decarboxylated in the BS (α), the BS allocation of mitochondrial area was assumed to be the BS fraction of day respiration (*f*_Rd_) in the leaf, and the BS allocation of chloroplast area set the BS allocation of rubisco (*f*_rubisco_). Rubisco density does not differ significantly between BS and M chloroplasts in C_3_ and C_2_ plants (Bauwe, 1984; Ueno, 2011; Stata, 2023). Values of *V_c_*_max_ and *g*_bs_ were estimated by Bayesian fitting in ‘emce’ (Foreman-Mackey *et al*., 2013). Model fitting at light saturation was restricted to an ambient CO_2_ concentration (*C_a_*) less than or equal to 400 µmol mol^-1^, where *A* is the RuBP-saturated net CO_2_ assimilation rate and the von Caemmerer (1989) formulation applies. From the fitted parameters we derived the glycine shuttle rate, BS CO_2_ concentration (*C_s_*), BS net CO_2_ assimilation rate (*A_s_*), and BS-to-M CO_2_ leakage rate (*L*) at *C_a_* = 400 µmol mol^-1^. We used this formulation rather than energetically explicit models (Bellasio & Farquhar, 2019) because we did not measure electron transport and because our objective was relative CO_2_ treatment effects when *A* is assumed to be RuBP saturated. Full equations, assumptions, parameter values, and sensitivity analysis procedure are described in Methods S7, with the results of sensitivity analyses in Fig. S7, measured versus fitted *A*/*C_i_* responses in Fig. S8, and representative fitted *A*/*C_i_* responses for individual plants in Fig. S9.

### Statistical analyses and principal component analyses

All statistical analyses were performed using R v4.5.2 and values from each of the four biological replicates per species and CO_2_ treatment, calculated as the means of any technical replicates (R Core Team, 2025). Growth CO_2_ treatment, photosynthetic phenotype, and their interaction were tested as fixed effects in linear mixed-effects models (LME; Pinheiro *et al*., 1999). Inclusion of a growth chamber term in linear mixed models did not improve fit (evaluated using Akaike Information Criterion, AIC) and yielded qualitatively identical results. Planned contrasts for CO_2_ effects within each photosynthetic phenotype were evaluated using ‘emmeans’ (Lenth & Piaskowski, 2017), with one-sided tests where the direction of effect was predicted *a priori* from the hypothesis that low CO_2_ acclimation strengthens C_2_ photosynthesis (e.g., decreased Γ and *C_i_*\*); two-sided tests were used otherwise. We report LME as the primary statistical analysis in the text.

To account for phylogenetic non-independence, CO_2_ treatment comparisons within photosynthetic phenotypes were repeated as phylogenetic LME (PLME), and differences among photosynthetic phenotypes (Table 1) were tested by phylogenetic generalized least squares (PGLS) on species means, each averaged across the two CO_2_ treatments, with pairwise contrasts corrected for false discovery rate (Benjamini & Hochberg, 1995). Both used a phylogeny pruned from published trees (Zanne *et al*., 2014; Smith & Brown, 2018; Jin & Qian, 2022). For PGLS and PLME, the correlation structure (Brownian motion, Ornstein-Uhlenbeck, or Pagel’s λ) was the simplest within ΔAIC < 2 of the lowest finite AIC.

**Table 1.**
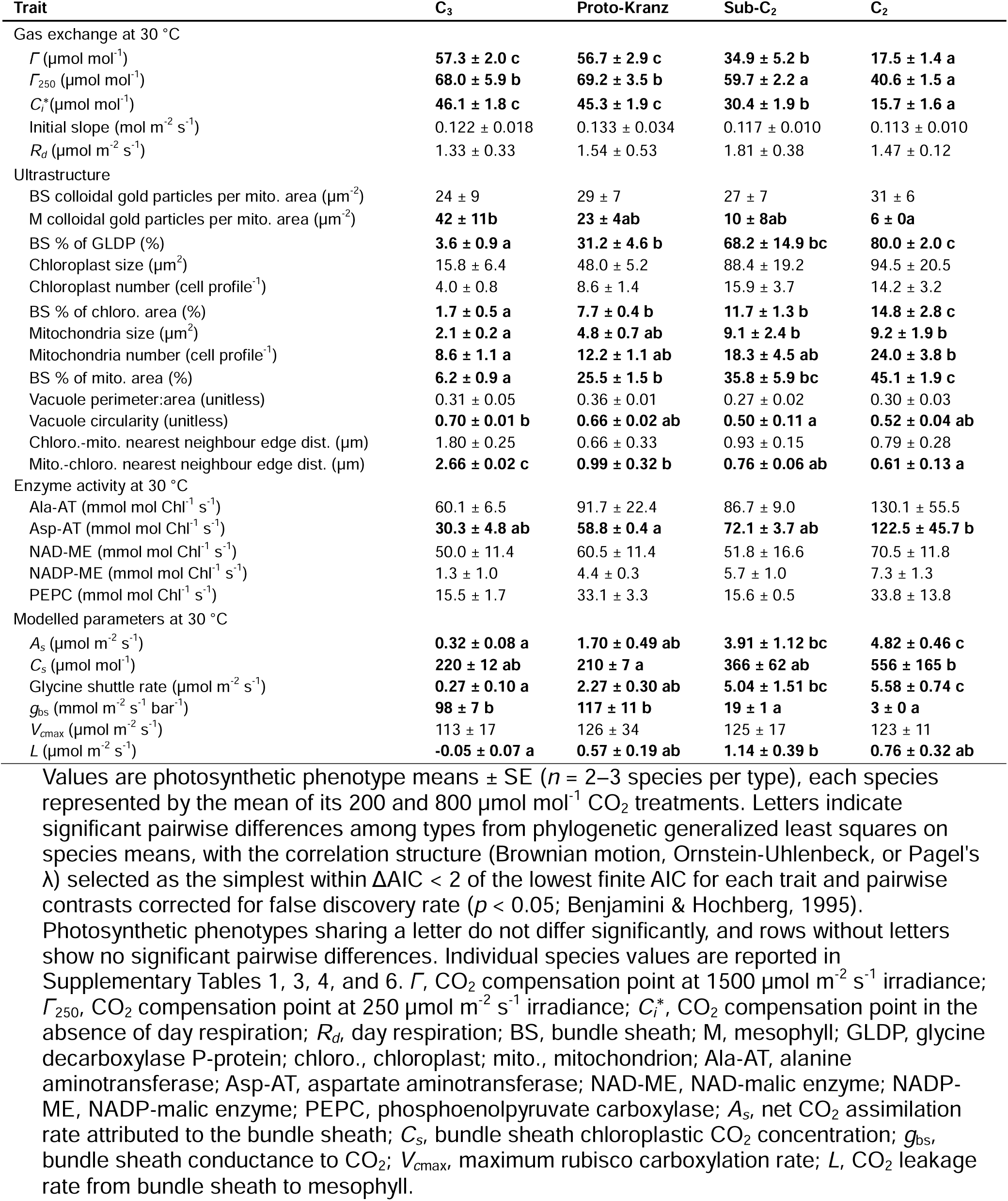
Trait differences among photosynthetic phenotypes.

The relationships of Asp-AT activity with glycine shuttle rate and of *C_i_*\* with *f*_GLDP_ were fitted by ordinary least squares on species means at each growth CO_2_. For each, we compared linear and quadratic forms, with and without a growth-CO_2_ interaction, and selected the simplest model within ΔAIC < 2 of the lowest AIC. The ordinary least squares and PGLS analyses gave consistent slopes. Species-level *t*-tests and a comparison of LME with PLME are reported in Tables S1-6.

Principal component analyses (PCA) used species means for each CO_2_ treatment as observations (*n* = 11 species x 2 CO_2_ treatments), performed with ‘prcomp’ in R v4.5.2 (R Core Team, 2025). The main PCA included the 15 traits showing significant CO_2_ treatment effects in the linear mixed-effects models. A PCA of all measured traits is shown in Fig. S10.

## Results

### Trait values of photosynthetic phenotypes

Mean trait values pooled across growth CO_2_ treatments establish the baseline against which the acclimation responses can be compared with the physiological shifts hypothesized to occur during C_2_ evolution (Table 1). Values of Γ and *C_i_*\* decreased from the C_3_ to the C_2_ phenotype (Fig. 1; Table 1). Relative to C_3_ species, sub-C_2_ and C_2_ species had greater chloroplast and mitochondrial area in BS cross sections, demonstrating greater organelle investment in the BS. Sub-C_2_ and C_2_ species also had greater GLDP immunolabelling, larger mitochondria, and shorter mitochondrion-to-chloroplast distances in BS cells than C_3_ species, and the C_2_ species additionally had more BS mitochondria than C_3_ relatives (Table 1). Proto-Kranz species differed from C_3_ species in having more chloroplast and mitochondrial coverage in the BS and closer spacing of BS chloroplasts and mitochondria. Mitochondria of the proto-Kranz, sub-C_2_, and C_2_ species were centripetally positioned in the BS, whereas mitochondria of the C_3_ species were dispersed around the BS periphery (Fig. 2; Fig. S3). Activities of Ala-AT were statistically similar between the four phenotypes, though the pooled values of sub-C_2_ and C_2_ species were 46% greater than those of the pooled C_3_ and proto-Kranz species (PLME *p* = 0.007; Table 1). Neither the activities of the two decarboxylases (NADP-ME and NAD-ME) nor that of PEPC differed between the four phenotypes, whereas Asp-AT activities were higher in sub-C_2_ than C_3_ species; the pooled Asp-AT activities of sub-C_2_ and C_2_ species were more than double those of the pooled C_3_ and proto-Kranz species (PLME *p* = 0.003; Table 1).

**Fig. 2.**
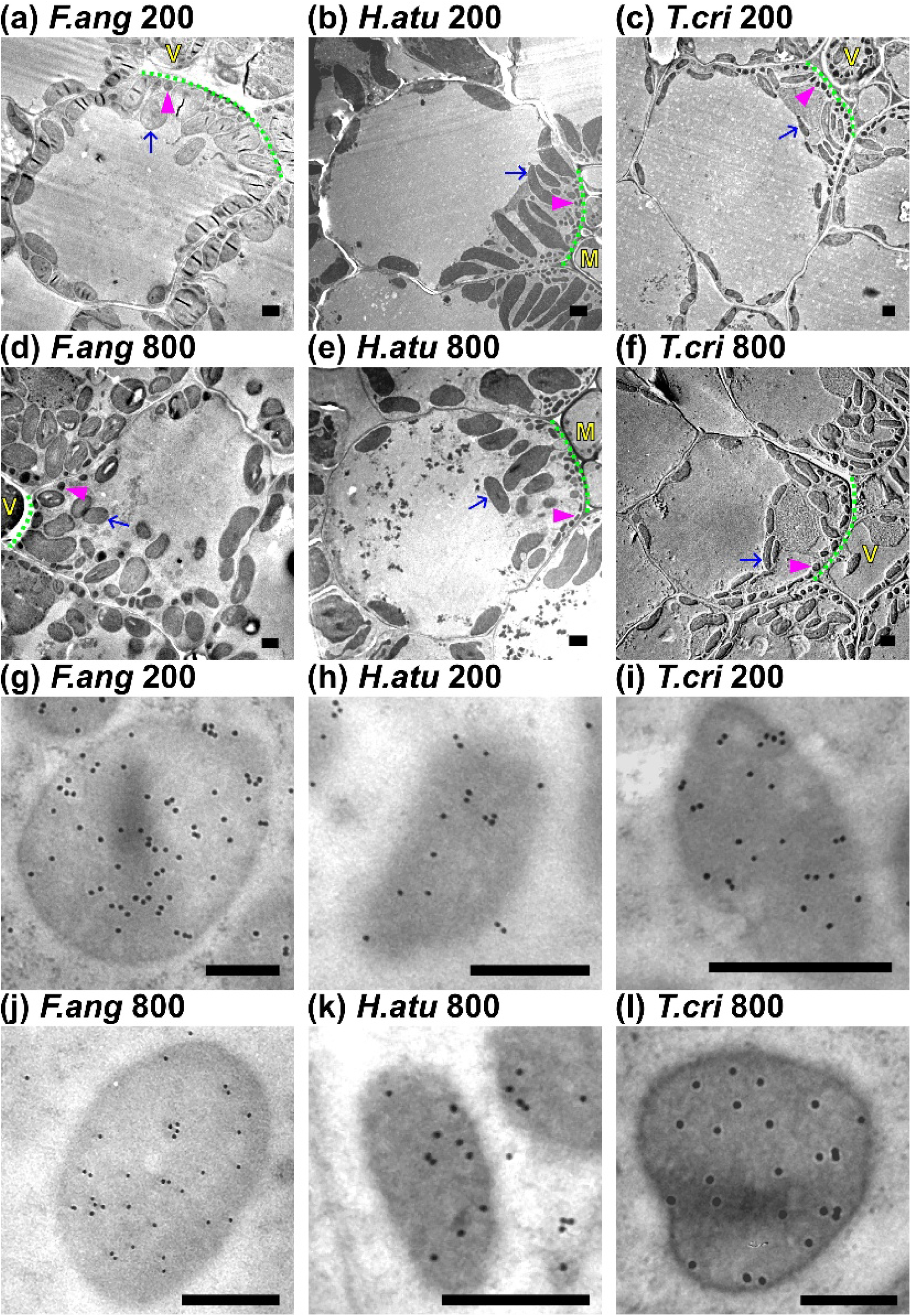
Ultrastructural plasticity in response to growth CO_2_. (**a–l**), Representative transmission electron micrographs of bundle sheath cells from three species representing independent C_2_ lineages. Whole-cell images are from plants grown at (**a–c**) 200 and (**d–f**) 800 µmol mol^-1^ CO_2_. High-magnification views of bundle sheath mitochondria are shown for plants grown at (**g–i**) 200 and (**j–l**) 800 µmol mol^-1^ CO_2_. Black dots in the high-magnification images are colloidal gold particles indicating the presence of the glycine decarboxylase P-protein (GLDP). Columns correspond to (**a,d,g,j**) Flaveria *angustifolia* (sub-C_2_), (**b,e,h,k**) *Homolepis aturensis* (C_2_), and (**c,f,i,l**) *Tribulus cristatus* (C_2_). Arrows, chloroplasts; arrowheads, mitochondria; M, mestome sheath; V, vein. The green dotted line indicates the cell wall in contact with the vein in eudicots or the mestome sheath in grasses. Scale bars: 2 µm (whole-cell images), 0.5 µm (mitochondrial images). For C_3_ species, see Fig. S3. For proto-Kranz species, see Fig. S4. For remaining sub- C_2_ and C_2_ species, see Fig. S5.

### Effects of growth CO_2_ treatment on gas exchange and enzyme activity

Mean Γ in sub-C_2_ plants grown at 200 µmol mol^-1^ CO_2_ was 18% lower than in sub-C_2_ plants grown at 800 µmol mol^-1^ CO_2_, and mean Γ in C_2_ species was 27% lower at 200 than at 800 µmol mol^-1^ CO_2_ (Table 2). Values of *C_i_*\* showed a larger CO_2_ treatment effect than Γ, being 29% lower in sub-C_2_ species and 42% lower in C_2_ species grown at 200 than at 800 µmol mol^-1^ CO_2_ (Table 2). By contrast, in the C_3_ and proto-Kranz species, neither Γ nor *C_i_*\* differed between plants grown at 200 and 800 µmol mol^-1^ CO_2_ (Table 2). The Γ measured at 250 µmol m^-2^ s^-1^ irradiance (Γ_250_) was also lower in C_3_ and sub-C_2_ species grown at 200 than at 800 µmol mol^-1^ CO_2_; in C_2_ species the treatment difference in Γ_250_ could not be resolved, although the means trended in the same direction as Γ and *C_i_*\* (Table 2). Estimated *R_d_* was lower at low than high growth CO_2_ in C_3_ species but not in the other photosynthetic phenotypes (Table 2). Treatment effects on Γ and *C_i_*\* within individual species were not always statistically significant, but the direction of the effect was consistent across sub-C_2_ and C_2_ species, enabling resolution of treatment effects within phenotypes (Table S1). The initial slope of the *A*/*C_i_* response at high light did not differ between treatments, indicating no apparent effect on rubisco carboxylation efficiency (Table 2).

**Table 2.**
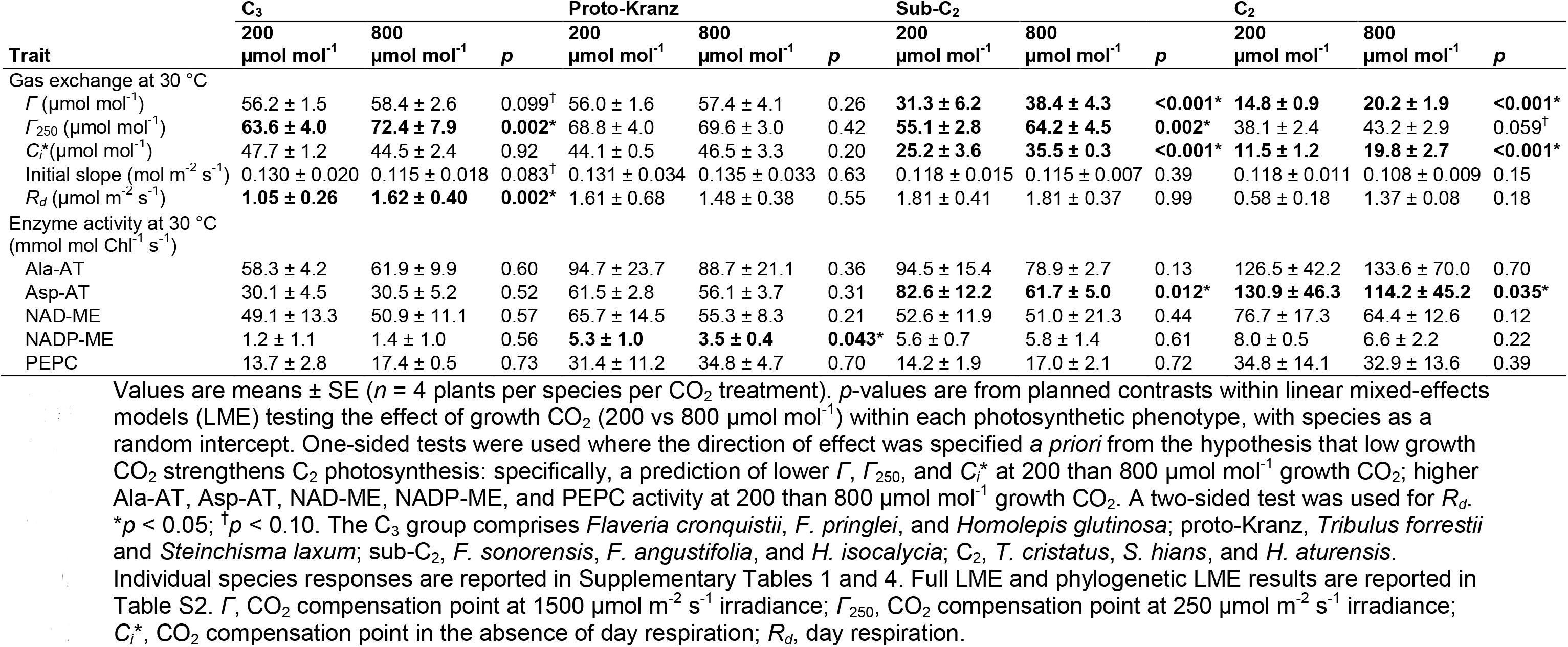
Effects of growth CO_2_ treatment on leaf gas exchange and enzyme activities at 30 °C.

Total leaf Asp-AT activity was 34% higher in sub-C_2_ species and 15% higher in C_2_ species when plants were grown at 200 relative to 800 µmol mol^-1^ CO_2_, while C_3_ and proto-Kranz species showed no significant difference between CO_2_ treatments (Table 2). Activities of PEPC, Ala-AT and NAD-ME did not differ between CO_2_ treatments in any photosynthetic phenotype (Table 2). For NADP-ME, the rate was higher in the proto-Kranz species grown at low than high CO_2_, though this pattern was driven primarily by *S. laxum* (Table 2; Table S4).

### Effects of growth CO_2_ treatment on leaf ultrastructural traits

In sub-C_2_ and C_2_ species grown at 200 µmol mol^-1^ CO_2_, chloroplasts and mitochondria were closely appressed along the centripetal wall of BS cells (Fig. 2a-f; Fig. S5). In these plants grown at 800 µmol mol^-1^ CO_2_, a greater number of chloroplasts were displaced from the vasculature towards the centrifugal pole of the BS cells (Fig. 2a-f; Fig. S5). Mean edge distance from chloroplasts to the nearest mitochondrion neighbour was 40% lower in sub-C_2_ species and 49% lower in C_2_ plants grown at low relative to high CO_2_ (Table 3). Shorter chloroplast-to-mitochondrion edge distances at low relative to high growth CO_2_ were directionally consistent in sub-C_2_ and C_2_ species in all four lineages (Table S3). Bundle sheath vacuoles were more circular in C_2_ species grown at low than high CO_2_, whereas vacuolar circularity in sub-C_2_ species did not differ between treatments (Table S3). C_3_ and proto-Kranz species showed no differences in chloroplast-mitochondrion distance or vacuole circularity between CO_2_ treatments (Table 3).

**Table 3.**
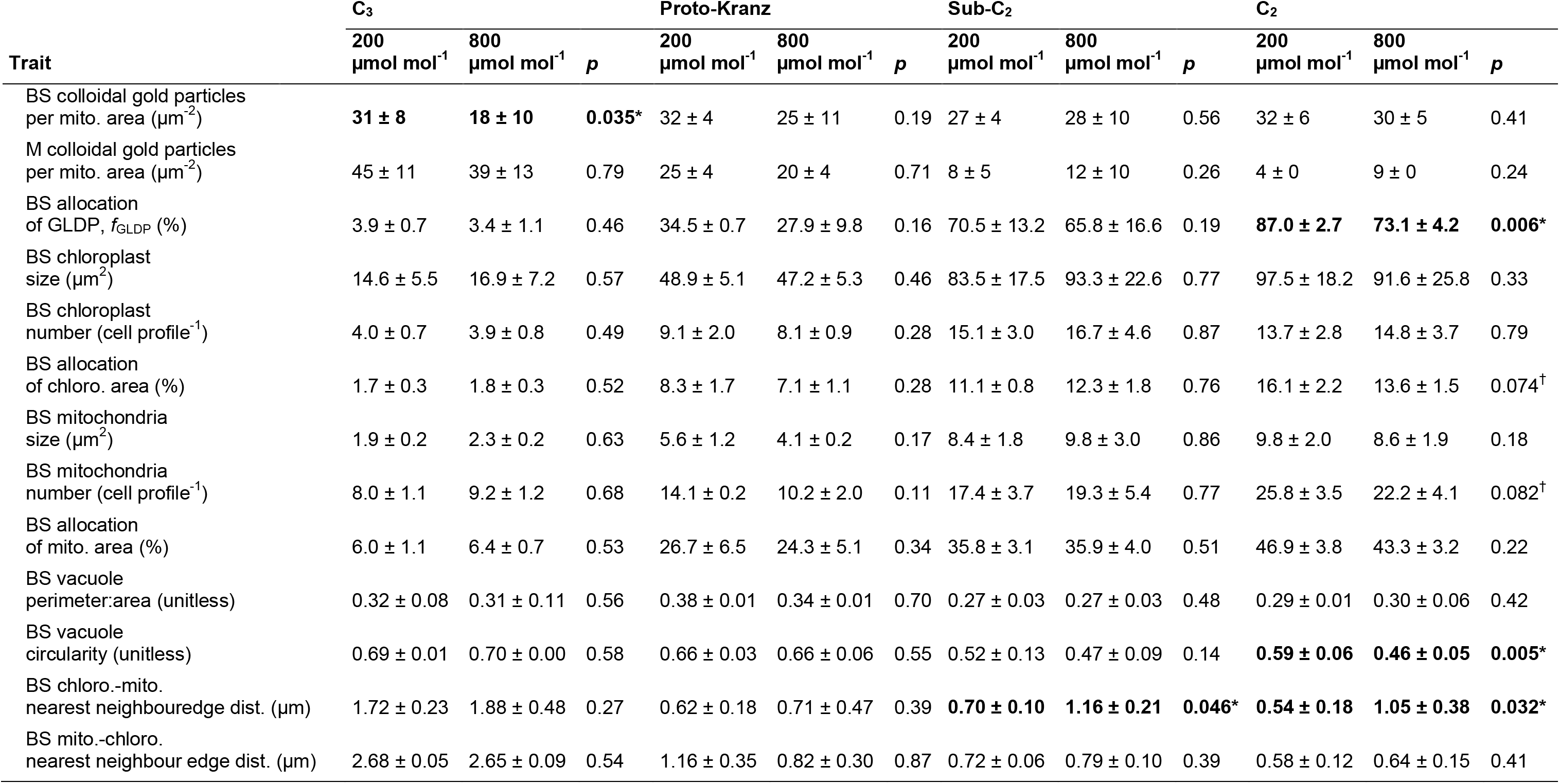
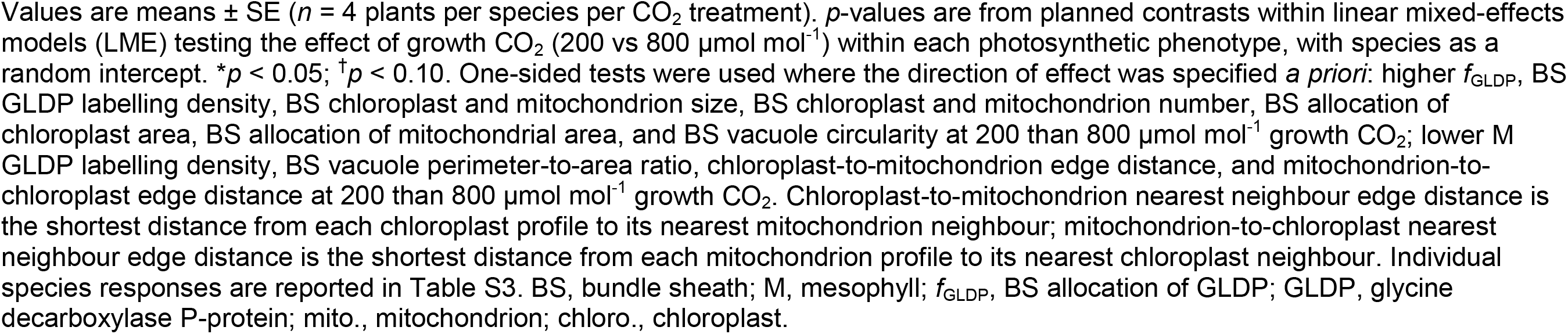
Effects of growth CO_2_ treatment on bundle sheath ultrastructure and GLDP immunolocalization on planar cross-sections.

The allocation of leaf GLDP to the BS, assessed as *f*_GLDP_, did not differ between CO_2_ treatments in C_3_, proto-Kranz, or sub-C_2_ species. Values of *f*_GLDP_ were 3–4% in C_3_, 28–35% in proto-Kranz, and 66–71% in sub-C_2_ species (Table 3; Table S3). The proto-Kranz species *T. forrestii* was an exception, showing higher *f*_GLDP_ and BS labelling density at low CO_2_ relative to high CO_2_ that were not observed in the other proto-Kranz species *S. laxum* (Table S3; Fig. S4e-h). In the C_2_ species, *f*_GLDP_ was 73% in plants grown at high CO_2_ and 87% in those grown at low CO_2_ (Fig. 2g-l; Table 3; Table S3; Fig. S5g-l). In C_2_ species grown at low relative to high CO_2_, respectively, GLDP labelling density in BS mitochondria was 32 and 30 gold particles µm^-2^ while their labelling density in M mitochondria was 4 and 9 gold particles µm^-2^ (Table 3). Neither difference in labelling density was individually significant, but the opposing directions of the BS and M responses produced the significant shift in *f*_GLDP_. The BS allocation of mitochondrial and chloroplast planar area was unchanged between CO_2_ treatments (Table 3). BS cell wall thickness, planar width, and planar perimeter per vein did not differ between growth CO_2_ treatments (data not shown).

### Modelled responses to growth CO_2_ treatment

A model of C_3_-C_4_ intermediate photosynthesis was applied to the gas exchange and ultrastructural data to evaluate how photosynthetic processes in the BS and M tissues responded to the growth CO_2_ treatments (Fig. 3; von Caemmerer, 1989). Sub-C_2_ and C_2_ species grown at 200 µmol mol^-1^ CO_2_ had higher predicted BS assimilation rates (*A_s_*) and higher glycine shuttle rates than plants of the same species grown at 800 µmol mol^-1^ CO_2_ (Table 4).

**Fig. 3.**
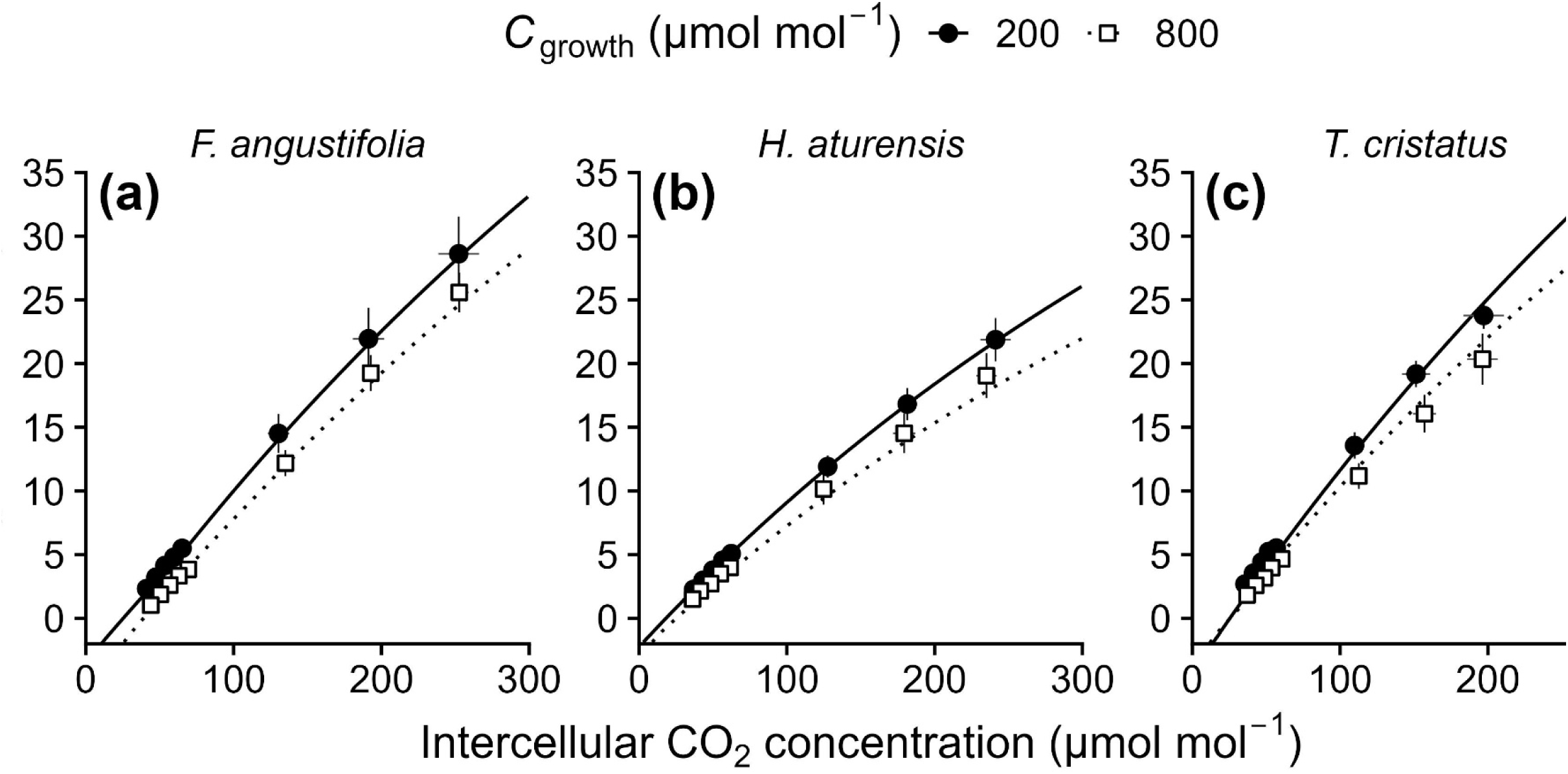
Observed and modelled response of net CO_2_ assimilation rate (*A*) to intercellular CO_2_ concentration (*C_i_*) at 1500 μmol m^-2^ s^-1^ irradiance and 30 °C in three representative species: (**a**) *Flaveria angustifolia* (sub-C_2_), (**b**) *Homolepis aturensis* (C_2_), and (**c**) *Tribulus cristatus* (C_2_). Points represent empirical gas exchange measurements (mean ± SE, n = 4 plants) measured at an irradiance of 1500 μmol m^-2^ s^-1^. Lines represent the mean of model predictions across biological replicates. Solid lines and closed circles are from plants grown at 200 μmol mol^-1^ CO_2_, while dotted lines and open squares are from plants grown at 800 μmol mol^-1^ CO_2_. The model was parameterized using measured ultrastructural parameters (bundle sheath allocation of GLDP and of chloroplast and mitochondrial planar area), while maximum rubisco carboxylation rate (*V_cmax)_*) and bundle sheath conductance (*g_bs_*) were estimated using a Bayesian framework (von Caemmerer, 1989; see *Materials and Methods*). For remaining species, see Fig. S8.

**Table 4.**
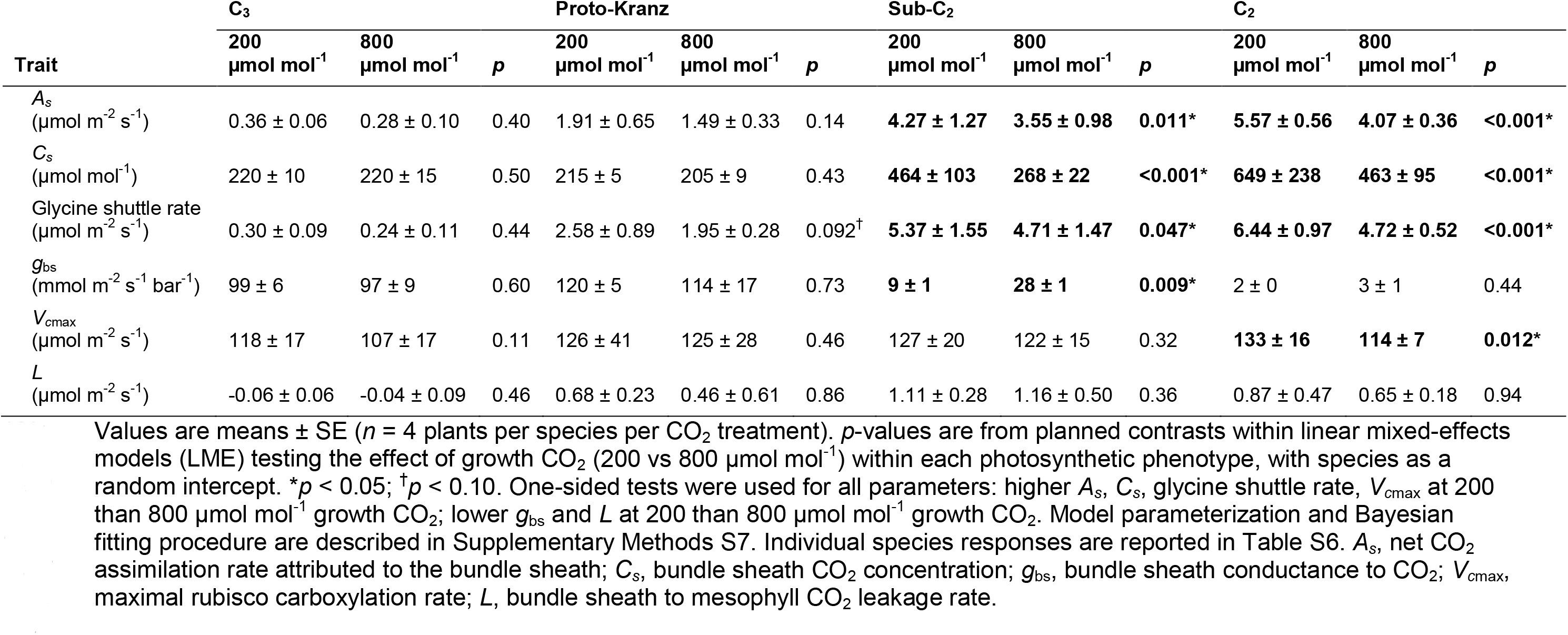
Effects of growth CO_2_ treatment on modelled photosynthetic parameters at 30 °C, irradiance of 1500 μmol m^-2^ s^-1^, and ambient CO_2_ of 400 μmol mol^-1^.

Modelled BS CO_2_ concentration (*C_s_*) was also 73% and 40% higher in the low than the high CO_2_ treatment in sub-C_2_ and C_2_ species, respectively (Table 4). Sub-C_2_ plants were modelled to have a *g*_bs_ of 28 mmol m^-2^ s^-1^ bar^-1^ at high growth CO_2_ and 9 mmol m^-2^ s^-1^ bar^-1^ at low growth CO_2_, a three-fold difference, whereas C_2_ species maintained a low *g*_bs_ of 2–3 mmol m^-2^ s^-1^ bar^-1^ regardless of CO_2_ treatment (Table 4). The modelled rubisco carboxylase activity (*V_c_*_max_) was higher at low relative to high growth CO_2_ only in C_2_ species (Table 4). Asp-AT activity scaled positively with modelled glycine shuttle capacity across species, and a single regression explained both CO_2_ treatments according to AIC (Fig. 4).

**Fig. 4.**
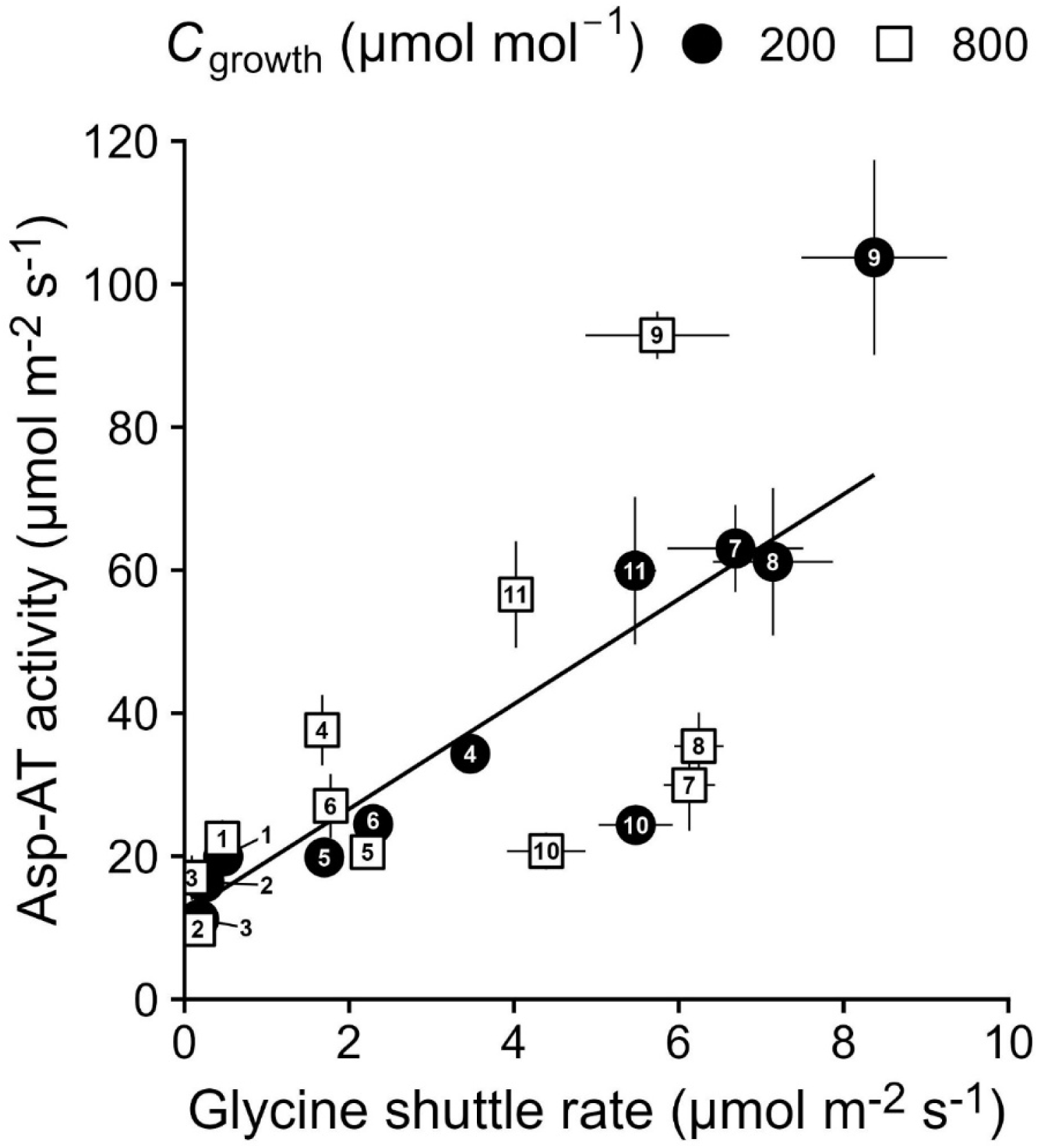
Relationship between modelled mesophyll-to-bundle sheath glycine shuttle rate at an ambient CO_2_ concentration of 400 μmol mol^-1^, 1500 μmol m^-2^ s^-1^ irradiance, and 30 °C versus *in vitro* aspartate aminotransferase activity at 30 °C (mean ± SE, *n* = 4 plants). A linear model was selected using the Akaike Information Criterion. Solid line, ordinary least-squares fit for all data shown: *y* = 11.9 + 7.33*x* (*R*^2^ = 0.59, *p* < 0.001). A phylogenetic generalized least-squares fit (Pagel’s λ) estimated a similar slope for the same relationship (slope = 8.27, *p* < 0.001). Closed circles, plants grown at 200 µmol mol^-1^ CO_2_; open squares, plants grown at 800 µmol mol^-1^ CO_2_. Species are numbered 1, *F. cronquistii*; 2, *H. glutinosa*; 3, *F. pringlei*; 4, *T. forrestii*; 5, *S. laxum*; 6, *H. isocalycia*; 7, *F. sonorensis*; 8, *F. angustifolia*; 9, *T. cristatus*; 10, *H. aturensis*; 11, *S. hians*.

### Acclimation of C_2_ photosynthesis in Flaveria sonorensis leaves

The acclimation responses described above could reflect developmental plasticity during leaf expansion rather than physiological plasticity within cells of mature tissues. To distinguish between developmental and physiological plasticity, plants of *F. sonorensis* (sub-C_2_) grown at 800 µmol mol^-1^ CO_2_ were transferred to 200 µmol mol^-1^ CO_2_ for two days and then returned to 800 µmol mol^-1^ CO_2_ (Fig. 5). Responses of *A* to *C_i_*were measured on leaves that had matured within the initial 800 µmol mol^-1^ growth CO_2_ condition (Fig. 5a-c). Values of Γ did not change significantly between time points (Fig. 5d), but *C_i_*\* halved within two days at low growth CO_2_ (from 41 to 19 µmol mol^-1^) and recovered to 38 µmol mol^-1^ upon return to high growth CO_2_ (Fig. 5e). Values of Γ_250_ declined from 84 to 60 µmol mol^-1^ at low growth CO_2_ and partially recovered to 68 µmol mol^-1^ after return to high CO_2_ (Fig. 5f). The rapid change of *C_i_*\* and Γ_250_ in leaves that had completed development at high CO_2_ indicates that elements of the acclimation response do not require new leaf growth.

**Fig. 5.**
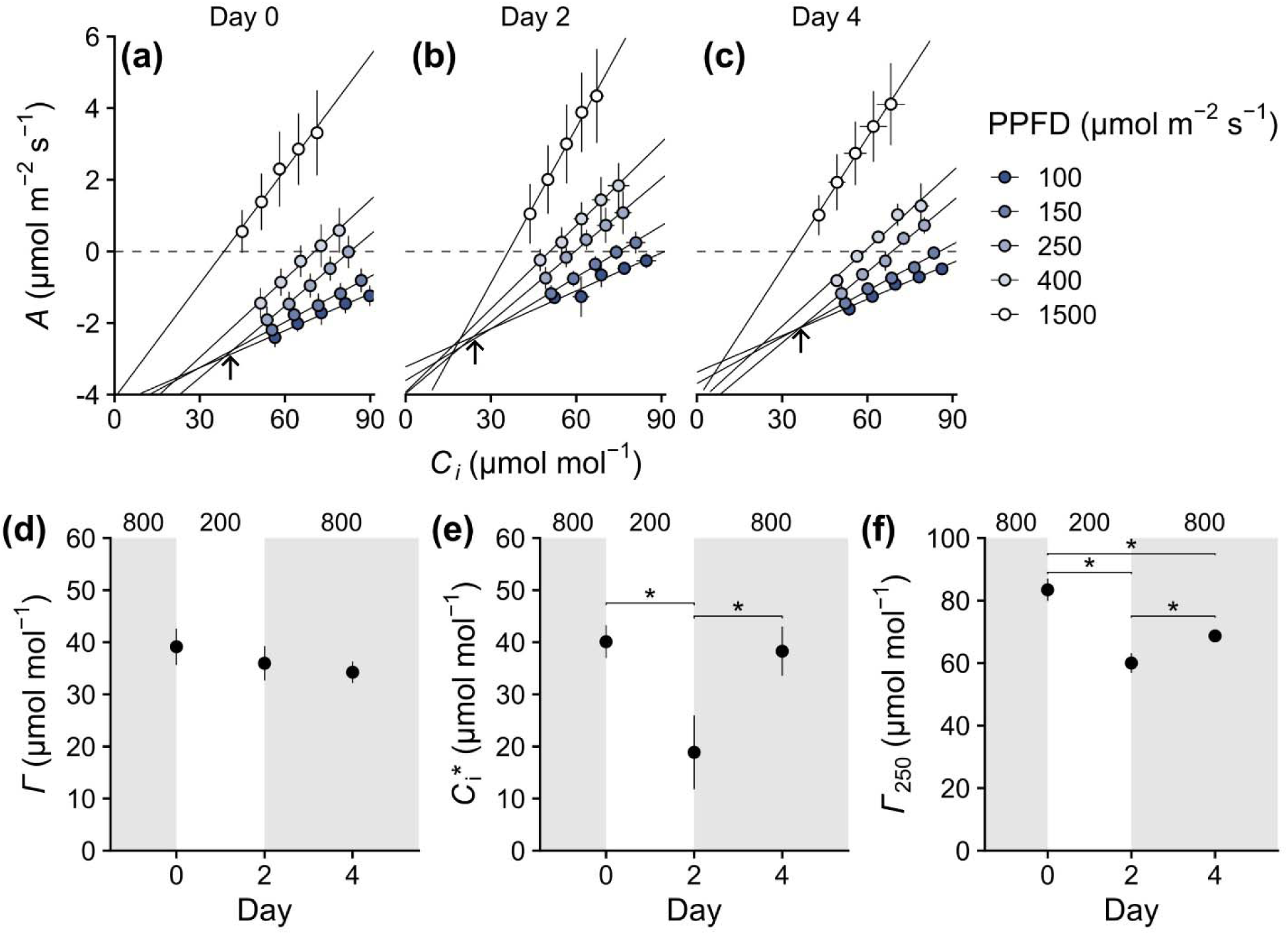
Time series of acclimation of the C_2_ photorespiratory glycine shuttle in mature leaves of *Flaveria sonorensis* following growth in 800 µmol mol^-1^ CO_2_. Plants were acclimated to high CO_2_ (800 µmol mol^-1^) for 7 days prior to the start of the experiment. On Day 0, baseline measurements were taken and plants were transferred to low CO_2_ (200 µmol mol^-1^). On Day 2, plants were returned to high CO_2_ and measured again on Day 4. Gas exchange measurements on all dates were performed on the same plants on leaves which were fully expanded and mature at Day 0. (**a-c**) Response of net CO_2_ assimilation rate (*A*) to intercellular CO_2_ concentration (*C_i_*) at 100, 150, 250, 400, and 1500 μmol m^-2^ s^-1^ irradiance and 30 °C, where arrows indicate the intersection point of lines regressed through the *A*/*C_i_* responses measured at the three lowest irradiance values. (**d**) The CO_2_ compensation point (Γ), (**e**) the CO_2_ compensation point in the absence of day respiration (*C_i_*\*), and (**f**) the CO_2_ compensation point at 250 μmol m^-2^ s^-1^ irradiance (Γ_250_) to changes in growth CO_2_ concentration. Grey shading indicates high CO_2_ conditions; unshaded region indicates the interval at low CO_2_. Asterisks indicate significant pairwise differences (*p* < 0.05, paired one-sided *t*-tests corrected for false discovery rate; Benjamini & Hochberg, 1995). Values are means ± SE (*n* = 4 plants).

### Direction of acclimation responses to growth CO_2_ treatment

Across species and growth CO_2_ treatments, *C_i_*\* declined linearly with increasing *f*_GLDP_ (Fig. 6). Sub-C_2_ and C_2_ species grown at 200 µmol mol^-1^ CO_2_ fell further along this line than the same species grown at 800 µmol mol^-1^ CO_2_, whereas C_3_ and proto-Kranz species clustered at *C_i_*\* values near 45 µmol mol^-1^ regardless of growth CO_2_ (Fig. 6).

**Fig. 6.**
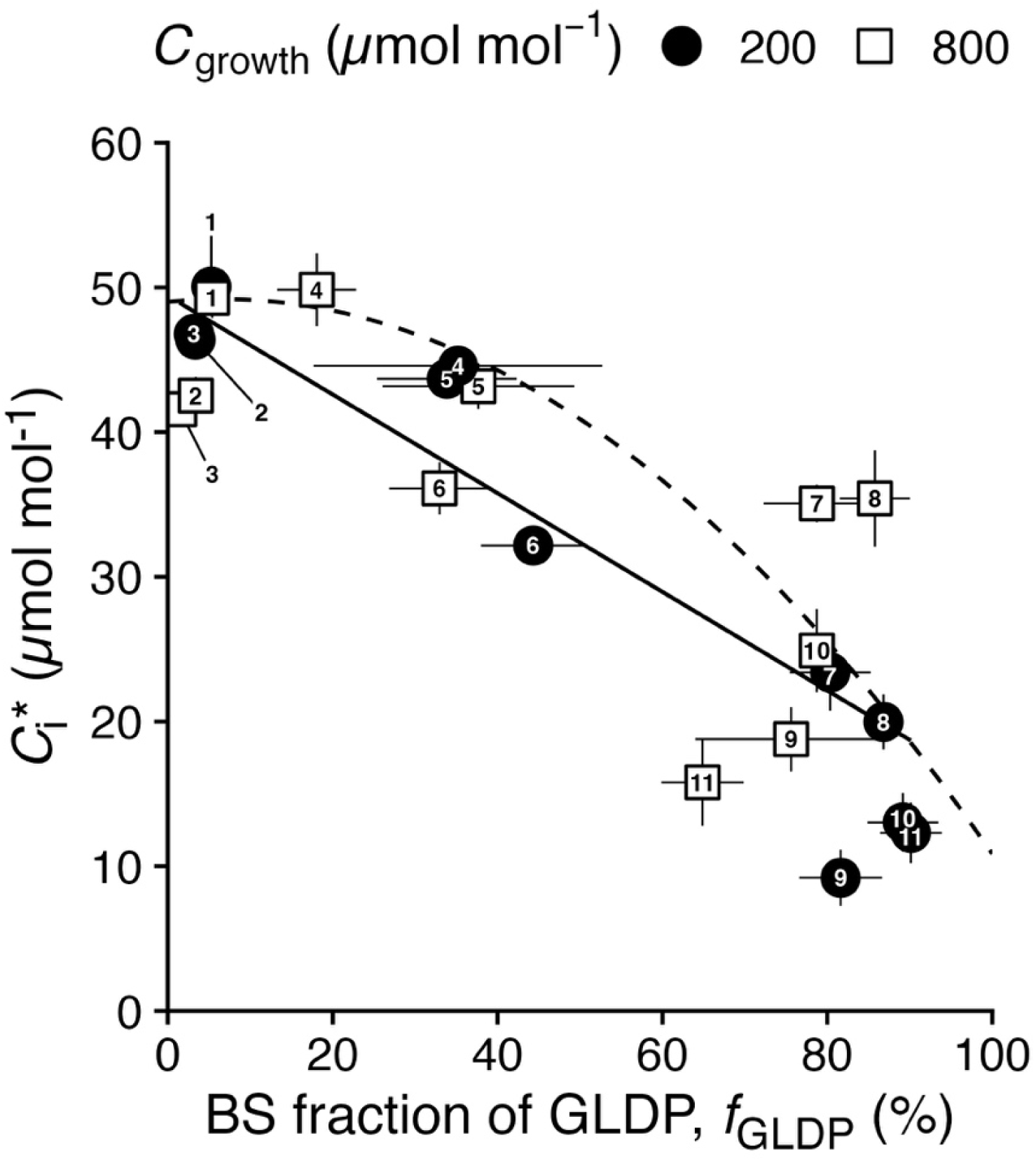
The CO_2_ compensation point in the absence of day respiration (*C_i_*\*) as a function of the bundle sheath (BS) allocation of leaf GLDP immunolabelling (*f*_GLDP_) across the C_3_ to C_2_ continuum in plants grown at 200 and 800 µmol mol^-1^ CO_2_. Data points are species means at each growth CO_2_ (mean ± SE, *n* = 4 plants). Solid line, ordinary least-squares regression: *y* = 49.4 - 0.341*x* (*R*^2^ = 0.73, *p* < 0.001). Phylogenetic generalized least squares (Pagel’s λ) gave a similar slope for the same relationship (-0.39, *p* < 0.001). Dashed line, the relationship reported for *Tribulus* by Leung *et al*. (2024): *y* = 49.0 + 0.0576*x* - 0.00439*x*^2^. Closed circles, 200 µmol mol^-1^; open squares, 800 µmol mol^-1^. Species codes: 1, *F. cronquistii*; 2, *H. glutinosa*; 3, *F. pringlei*; 4, *T. forrestii*; 5, *S. laxum*; 6, *H. isocalycia*; 7, *F. sonorensis*; 8, *F. angustifolia*; 9, *T. cristatus*; 10, *H. aturensis*; 11, *S. hians*.

We then assessed whether low versus high growth CO_2_ responses were random or whether they differed together in similar directions toward a C_2_ phenotype. A PCA was therefore restricted to the 15 traits with significant differences between CO_2_ treatments; a PCA using all traits gave qualitatively similar results (Fig. S10). The PCA produced a dominant axis (PC1, 57% of variance) separating C_3_ and C_2_ phenotypes, with positive loadings on *f*_GLDP_, glycine shuttle rate, *C_s_*, *A_s_*, Asp-AT, and NADP-ME, and negative loadings on Γ, Γ_250_, *C_i_*\*, *g*_bs_, nearest neighbour distances of BS chloroplast to mitochondria, and BS vacuole circularity (Fig. 7a,b). On the second principal component (PC2), eudicot species had more positive values and grass species more negative values, explained by *R_d_*, *V_c_*_max_, and BS labelling density of GLDP (Fig. 7a). Sub-C_2_ and C_2_ species grown at 200 µmol mol^-1^ CO_2_ were shifted towards the C_2_ end of PC1 relative to the same species grown at 800 µmol mol^-1^ CO_2_ (paired one-sided *t*-test; *t* = 3.62, *p* = 0.002; Fig. 7a), whereas C_3_ and proto-Kranz species showed no consistent shifts along PC1 between CO_2_ treatments. Acclimation therefore moved phenotypes along PC1 towards enhanced C_2_ characteristics (Fig. 7a).

**Fig. 7.**
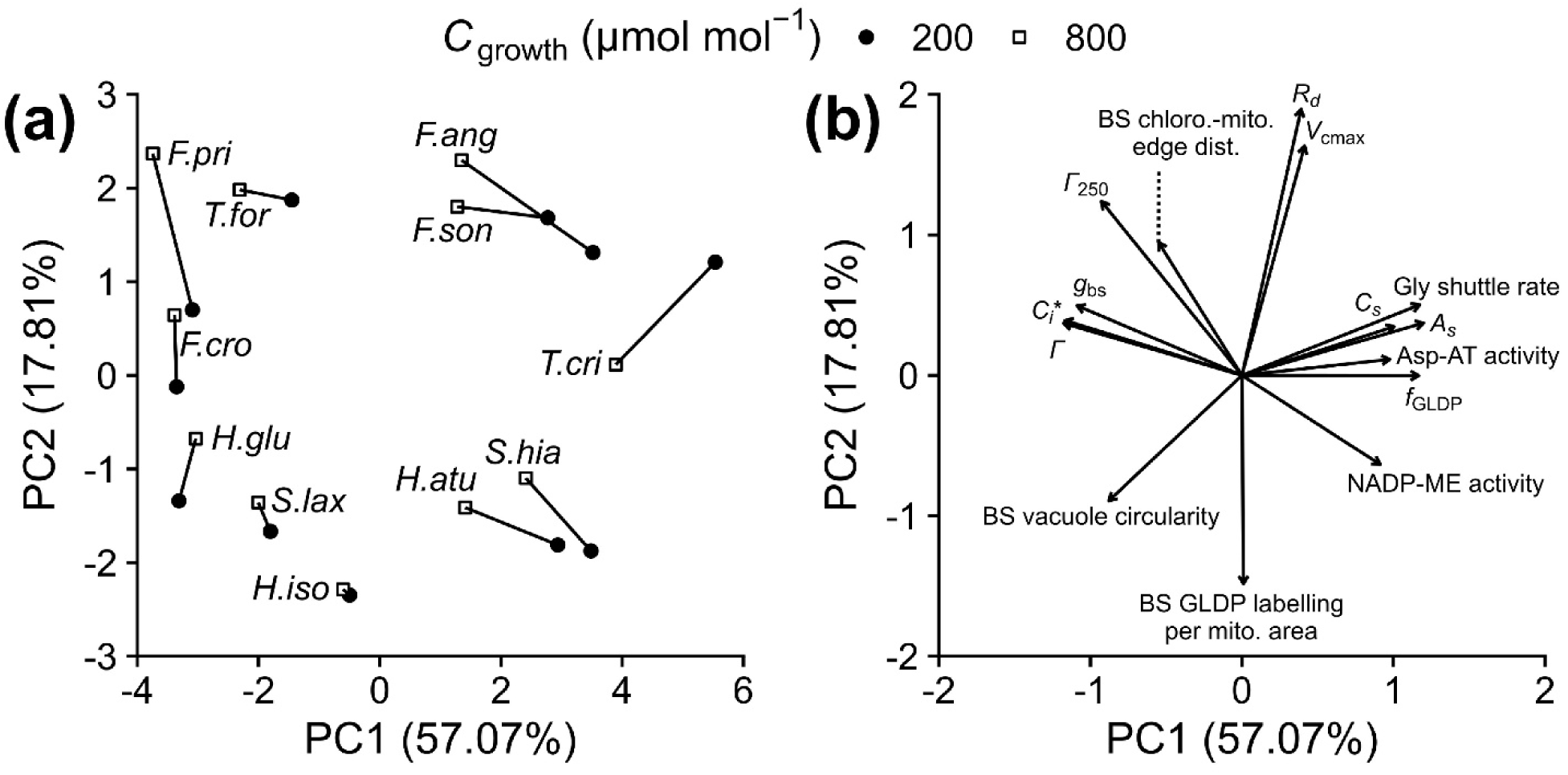
Assessment of leaf phenotypic plasticity across the C_3_ to C_2_ continuum. Principal component analysis (PCA) of 15 physiological, biochemical, and ultrastructural traits measured in plants from two growth CO_2_ concentrations (*C*_growth_). Traits showing significant CO_2_ treatment effects in the LME analysis were included (see Tables 2, 3, and 4). A PCA of the full trait set is shown in Fig. S10. (**a**), Species scores are linked by lines representing the plastic response to low CO_2_. Closed circles, 200 ; open squares, 800 µmol mol^-1^. Acclimation to low CO_2_ shifted phenotypes towards more positive PC1 values (paired one-sided *t*-test; *t* = 3.62, df = 10, *p* = 0.002; mean shift = 0.86 PC1 units). The C_3_ species *H. glutinosa* shifted in the opposite direction. (**b**), Loading vectors indicating that PC1 captures 57% of variance across C_2_-related traits. Negative loadings correspond to higher CO_2_ compensation points (Γ, Γ_250_), CO_2_ compensation point in the absence of day respiration (*C_i_*\*), bundle sheath conductance (*g*_bs_), BS chloroplast-mitochondrion edge distance, and BS vacuole circularity. Positive loadings correspond to higher BS CO_2_ concentration (*C_s_*), aspartate aminotransferase (Asp-AT) activity, glycine (Gly) shuttle rate, BS net CO_2_ assimilation rate (*A_s_*), BS allocation of GLDP (*f*_GLDP_), and NADP-malic enzyme (NADP-ME) activity. PC2 (17% of variance) was correlated with lower BS GLDP labelling per mitochondria area, day respiration rate (*R_d_*), and maximum rubisco carboxylation rate (*V_c_*_max_). Species abbreviations: *F.cro*, *F. cronquistii*; *F.pri*, *F. pringlei*; *H.glu*, *H. glutinosa*; *T.for*, *T. forrestii*; *S.lax*, *S. laxum*; *H.iso*, *H. isocalycia*; *F.son*, *F. sonorensis*; *F.ang*, *F. angustifolia*; *T.cri*, *T. cristatus*; *H.atu*, *H. aturensis*; *S.hia*, *S. hians*.

## Discussion

Enhanced photorespiratory potential caused by a reduction in growth CO_2_ produced clear acclimation responses in and only in photosynthetic phenotypes operating a M-to-BS glycine shuttle. Sub-C_2_ and C_2_ species whose BS tissues had substantial organelle and GDC investment strengthened the M-to-BS glycine shuttle when grown under high photorespiratory conditions, whereas C_3_ and proto-Kranz species did not. Acclimation therefore does not appear to accumulate gradually along the C_3_ to C_2_ continuum, but instead appears once the BS can metabolize glycine rapidly enough to maintain net M-to-BS glycine diffusion. Acclimation potential may thus require ‘potentiation’, where acquisition of specific character states enable a phenotype to respond plastically to altered photorespiratory potential. In the context of C_3_-C_4_ evolutionary intermediates, we hypothesize that this acclimation may push relatively cryptic character states in a population across a phenotypic threshold beyond which selection is intensified. By doing so, acclimation could facilitate evolution of photosynthetic complexity in these lineages, leading to the CO_2_-concentrating mechanisms of C_2_, and subsequently, C_4_ photosynthesis.

### Sub-C_2_ species acclimate to low relative to high growth CO_2_ by repositioning organelles, thereby lowering bundle sheath conductance

In sub-C_2_ species, acclimation to 200 relative to 800 µmol mol^-1^ growth CO_2_ involves modification of BS cell ultrastructure that enhances retention of photorespired CO_2_. Our model of C_2_ photosynthesis predicts lower *g*_bs_, higher *C_s_*, and higher *A_s_* in the plants of the low relative to high growth CO_2_ treatment (Table 4). The closer packing of chloroplasts and mitochondria along the inner BS wall at low relative to high growth CO_2_ can explain, at least in part, the reduced *g*_bs_ predicted for this treatment (Table 3).

Closer packing of chloroplasts against mitochondria increases the length and tortuosity of the diffusion path that CO_2_ must travel to exit the BS, and more importantly places rubisco-rich compartments along the efflux pathway (von Caemmerer & Furbank, 2003). Because rubisco continuously draws down chloroplast CO_2_, closer organelle positioning favours the diffusive influx of photorespired CO_2_ into the chloroplast rather than out of the BS (Evans *et al*., 2009; Tholen & Zhu, 2011). This situation parallels the role of chloroplast coverage of the M cell periphery in refixing photorespired and respired CO_2_ in M cells of C_3_ plants (Busch *et al*., 2013; Peixoto *et al*., 2021). Other structural mechanisms proposed to reduce *g*_bs_, such as increased vacuole size and circularity (Table 3) or thicker BS cell walls (data not shown) were not apparent in sub-C_2_ plants. The stronger glycine shuttle predicted in sub-C_2_ plants was associated with greater leaf Asp-AT activity, but not with greater BS GLDP investment nor with greater activities of C_4_ cycle enzymes such as PEPC, NAD-ME, and NADP-ME (Tables 2, 3, 4). Greater Asp-AT activity could improve ammonia reassimilation in the BS and accelerate nitrogen return to the M tissues (Mallmann et al., 2014), as discussed below for C_2_ species.

### C_2_ species acclimate to low relative to high growth CO_2_ by increasing glycine decarboxylation in the bundle sheath

In C_2_ species, acclimation to declining CO_2_ was evident in both structural and biochemical parameters. Structurally, BS chloroplasts were larger and mitochondria more numerous, which in turn contributed to a greater allocation of leaf GLDP to the BS (higher *f*_GLDP_); vacuole circularity was greater and the distance from chloroplasts to the nearest mitochondrion was shorter in low- than high-CO_2_ grown plants (Table 3). Biochemically, leaf Asp-AT activity and modelled *V_c_*_max_ were greater in low CO_2_-grown C_2_ plants (Tables 2, 4). The structural and biochemical differences were associated with greater glycine shuttle rate, *C_s_*, and *A_s_*, but not with a change in *g*_bs_ (Table 4). C_2_ species maintained a low modelled *g*_bs_ resembling that of C_4_ plants at both CO_2_ treatments (von Caemmerer & Furbank, 2003), indicating that low *g*_bs_ is constitutive in C_2_ species.

Concentrating the large fraction of leaf GLDP in the inner BS of C_2_ species places photorespiratory CO_2_ release in the chloroplast-rich region of the BS, where diffusive efflux is already slowed by the resistance of abundant chloroplasts, a strong CO_2_ sink represented by rubisco, and a large vacuole (Rawsthorne *et al*., 1988b; Hylton *et al*., 1988; Morgan *et al*., 1993; von Caemmerer & Furbank, 2003). The higher *f*_GLDP_ at 200 µmol mol^-1^ CO_2_ was better explained by a greater number of BS mitochondria rather than from denser GLDP labelling within them, since labelling density per mitochondrion differed only slightly between CO_2_ treatments in either cell type (Table 3). Acclimation therefore differs from the developmental pattern observed previously in leaves of C_3_ pea, in which GDC proteins accumulate within pre-existing mitochondria and raise their matrix density (Vauclare *et al*., 1996). Instead, the acclimation pattern resembles the increase in mitochondrial number that accompanies growth at elevated CO_2_ in C_3_ species such as soybean, red maple, and lobolly pine (Griffin *et al*., 2001). The whole-leaf increase in GLDP transcripts observed in intact leaves of *F. sonorensis* grown at 100 relative to 400 µmol mol^-1^ CO_2_ (Wang *et al*., 2026) is also consistent with the proliferation of BS mitochondria at a constant GDC content per organelle. Higher *C_s_* and *A_s_* in sub-C_2_ and C_2_ species at low relative to high growth CO_2_ may increase the demand for cellular energy in the BS, analogous to the higher CO_2_ concentration and *A* of M cells in C_3_ species grown at elevated CO_2_ (Griffin *et al*., 2001).

### C_3_ and proto-Kranz species are largely unresponsive to variation in photorespiratory potential as modulated by growth CO_2_ supply

Growth at 200 versus 800 µmol mol^-1^ CO_2_ generally did not produce significant treatment effects in C_3_ and proto-Kranz species, and where effects were observed, they varied among species within a photosynthetic phenotype. Organelle size, number, and juxtaposition in the BS were unchanged by the growth CO_2_ treatments, as were the measured enzyme activities. C_3_ species had more GLDP labelling per mitochondrial area in both the M and BS at low than high growth CO_2_, but reductions in mitochondrial size and number in the BS offset the higher labelling density, such that GDC allocation to the BS was unchanged by the CO_2_ treatment. Modelled glycine shuttle rates, *g*_bs_, *A_s_*, and *C_s_* were not significantly affected by growth CO_2_ treatment, demonstrating no obvious acclimation in C_2_-like physiology.

The reduction in Γ values in the low CO_2_-grown C_3_ plants would be consistent with stronger glycine shuttling, but could also be explained by the lower *R_d_* relative to *V_c_*_max_ predicted for the C_3_ plants grown in the low relative to high CO_2_ treatments (Brooks & Farquhar, 1985; Sage & Pearcy, 1987). Under the nutrient-rich growth conditions used here, C_3_ species often show little acclimation to declining CO_2_ (Sage & Coleman, 2001), and the acclimation responses of photorespiratory parameters is often small and inconsistent (Campbell *et al*., 2005; Vogan & Sage, 2012; Lundgren *et al*., 2019). Among the proto-Kranz species, *T. forrestii* was modelled to have a greater glycine shuttle rate when grown at low versus high CO_2_, suggesting proto-Kranz species can share some acclimation responses as observed in sub-C_2_ species; however, *S. laxum* showed no change, indicating variable acclimation potential in the proto-Kranz phenotype.

### Aspartate aminotransferase capacity tracks modelled glycine shuttle rates across growth CO_2_ treatments and photosynthetic phenotypes

Common garden studies widely show that C_2_ genotypes have enhanced activity of Asp-AT and Ala-AT or levels of transcripts encoding these proteins, relative to C_3_ and proto-Kranz genotypes, indicating upregulation of aspartate and alanine shuttles that are proposed to sustain C_2_ photosynthesis (Mallmann *et al*., 2014; Dunning *et al*., 2019; Adachi *et al*., 2023; Leung *et al*., 2024; Stata *et al*., 2025). Whether Asp-AT and Ala-AT activities are plastic had not previously been tested. Here, we show that Asp-AT capacity is not fixed but is enhanced under low versus high growth CO_2_. Notably, Asp-AT activity scales with the modelled glycine shuttle rate (Fig. 4). By contrast, Ala-AT activity does not respond to growth CO_2_ nor correlate with the glycine shuttle rate (Table 2). Consistently, Wang *et al*. (2026) reported increased transcript levels of plastid Asp-AT in *F. sonorensis* and *F. linearis* grown at 100 relative to 400 µmol mol^-1^ CO_2_. We hypothesize that Asp-AT activity therefore tracks the demand for nitrogen recycling created by a stronger glycine shuttle, regardless of whether demand arises from acclimation or from evolutionary divergence among species.

Enhancing Asp-AT activity in C_2_ species is proposed to assist the leaf in compensating for any stoichiometric nitrogen imbalance between M and BS tissues caused by M-to-BS glycine shuttling (Mallmann *et al*., 2014). Two glycine molecules are decarboxylated by GDC and serine hydroxymethyltransferase in the BS for every serine returned to the M. The two glycine each import one amino group into the BS. While serine returns one of these amino groups to the M, the other is released as NH_3_ during glycine decarboxylation (Bauwe *et al*., 2010). Sustained glycine synthesis from glyoxylate thus requires an equivalent supply of amino groups returned to the M following glycine decarboxylation (Rawsthorne *et al*., 1988a). For this to occur, NH_3_ released by glycine decarboxylation in the BS is reassimilated into glutamate which can diffuse back to the M; however, rapidly returning amino groups to the M by glutamate diffusion alone would require a steep BS-to-M glutamate concentration gradient. This may be problematic because high BS glutamate concentrations required could feed back negatively onto ammonia reassimilation (Miller & Stadtman, 1972). Another consideration is the chemical lability of 2- oxoglutarate, which may limit its ability to diffuse from the M to the BS. Hydrogen peroxide can oxidatively decarboxylate 2-oxoglutarate, and peroxisomes, the site of its production, are often clustered against the chloroplasts that house the plastidic glutamine synthase (GS) and GOGAT isoforms, and against the mitochondria (Vlessis *et al*., 1990). Failure to reassimilate photorespired ammonia as fast as it is produced could promote progressive ammonia efflux from the leaf, thereby aggravating any nitrogen deficiency of the plant, as well as slowing glycine formation in the M (Kumagai *et al*., 2011). Transamination of glutamate with pyruvate or oxaloacetate in the BS yields alanine or aspartate, respectively, which could diffuse to the M where the reverse reaction regenerates glutamate that would donate the second amino group to glyoxalate during glycine synthesis (Bauwe *et al*., 2010; Mallmann *et al*., 2014). We thus hypothesize that phenotypically plastic engagement of the aspartate and alanine aminotransferases in sub-C_2_ and C_2_ species appears to enable the BS to avoid high glutamate concentrations while sustaining glycine synthesis in the M.

### An anatomical threshold influences plasticity of C_2_ photosynthesis, which may facilitate the evolution of C_4_ photosynthesis

Values of *f*_GLDP_ and *C_i_*\* are inversely related across the C_3_ to C_2_ continuum at 30 °C (Fig. 6), as reported previously across three lineages (Leung *et al*., 2024). The low variation in *C_i_*\* below an *f*_GLDP_ of ∼33% shown in Fig. 6 indicates a threshold of BS investment of leaf GLDP is required for substantial acclimation. This threshold we propose is around an *f*_GLDP_ value of above 33%, above which greater *f*_GLDP_ corresponds to declines in *C_i_*\*. The decline in *C_i_*\* in this case reflects the reduction in rubisco oxygenation relative to carboxylation in the whole leaf, caused by the greater rubisco efficiency at the elevated CO_2_ concentrations within the BS chloroplasts (von Caemmerer, 1989; Sage *et al*., 2013). The PCA of 15 traits also supports this interpretation; acclimation responses are largest and most consistent in terms of their direction of change above a PC1 value of near zero yet are small and inconsistent below a PC1 of zero (Fig. 7a). A PC1 value of zero corresponds to an *f*_GLDP_ of 47%, a *C_i_*\* of 33 µmol mol^-1^, a *g*_bs_ of 54 mmol m^-2^ s^-1^ bar^-1^, and a modelled glycine shuttle rate of 3.4 µmol m^-2^ s^-1^, values that fall between the means of proto-Kranz and sub-C_2_ phenotype (Table 1).

The absence of comparable growth CO_2_ responses in leaves of C_3_ and proto-Kranz species supports a hypothesis that a threshold of BS potentiation is required before M-to-BS glycine shuttling can acclimate to photorespiratory conditions. Where BS tissues contain few mitochondria and little GDC, as in the C_3_ and proto-Kranz species examined here, the glycine sink potential of the BS may be too low to influence leaf glycine metabolism. Conversely, where the BS contains sufficient mitochondria, GDC, and rubisco, rapid metabolism of glycine spillover from the M could maintain a steep glycine gradient along which excess glycine diffuses into the BS. Glycine flow into the BS during photorespiratory surges could in turn stimulate induction of GDC expression and mitochondrial biogenesis and reinforce the acclimation response (Griffin *et al*., 2001; Timm *et al*., 2013). Once BS cells become sufficiently organelle-rich and metabolically active, high photorespiratory conditions may then induce latent traits that strengthen C_2_ character states.

Where enablers are quantitative traits, crossing specific thresholds may facilitate the evolution of another trait (Blount *et al*., 2008; Edwards, 2023). We hypothesize that if a threshold governs the ability of C_2_ photosynthesis to acclimate to high photorespiration, certain C_2_ characteristics may remain cryptic or latent until the necessary environmental cue(s) and leaf phenotypic context coincide. The transition from proto-Kranz to sub-C_2_ phenotypes may represent such thresholds which potentiate acclimation and subsequent evolution (West-Eberhard *et al*., 2011; Edwards, 2023; Leung *et al*., 2024). If so, acclimation may affect evolutionary potential in the opposite manner that is often presumed in plant physiological traits. Commonly, acclimation may constrain rather than promote evolutionary change. By maintaining fitness near its optimum across environments, acclimation shields the underlying genotype from selection and thus weakens selection on more derived character states (Huey *et al*., 2003; Ghalambor *et al*., 2007). In the case of certain traits such as C_2_ photosynthesis, acclimation may have a facilitating effect on selection; the low versus high growth CO_2_ increased the phenotypic disparity between the C_3_ to C_2_ phenotypes, which could increase opportunity for directional selection (Fig. 7a; Paaby & Rockman, 2014). Increased phenotypic variation could also expand the realized ecological niche of a population, likewise increasing opportunity for selection (Ghalambor *et al*., 2007; Lundgren *et al*., 2014). Plasticity could then facilitate, rather than impede, the repeated origin of C_2_ and ultimately C_4_ photosynthesis (West-Eberhard *et al*., 2011).

## Conclusions

Leaf photorespiratory fluxes in the hot, seasonally dry habitats occupied by C_2_ species and their close C_3_ relatives vary strongly over diel and seasonal timescales as temperature, salinity, water availability, and intercellular CO_2_ concentration vary (Huxman *et al*., 2004; Sage *et al*., 2018).

An enhanced C_2_ phenotype is adaptive under strong photorespiratory conditions, as evidenced by higher photosynthetic rates and temperature optima in C_2_ species than C_3_ species (Schuster & Monson, 1990; Monson & Jaeger, 1991; Vogan & Sage, 2012; Leung *et al*., 2024), and by faster growth at high temperature in C_2_ genotypes with lower Γ relative to those with higher Γ (Teese, 1995). Selection in ancestral populations may therefore have acted not on phenotypes expressed under benign conditions but on phenotypes induced during high photorespiratory conditions. Where C_2_ photosynthesis is both adaptive and plastic, acclimation could facilitate evolution through what is termed genetic assimilation, whereby a plastic trait becomes constitutive and is expressed without the environmental cue (Waddington, 1953). The difference in *g*_bs_ between growth CO_2_ treatments is three-fold in sub-C_2_ species and is near-absent in C_2_ species (Table 4), as would occur if an ancestrally plastic *g*_bs_ became constitutively expressed. Plasticity may thus assist genotypes moving across troughs in the C_4_ evolutionary fitness landscape (Sage, 2004; Heckmann, 2016). The frequent evolution of one of the most complex traits in plants may be explained, in part, by plasticity-facilitated evolution.

## Acknowledgements

The authors thank Alice DesRoches and Thomas Gludovacz for assistance at the growth facilities, Kenana Al Kakouni and Audrey Chong for assistance at the imaging facilities, USDA-ARS GRIN, Corey Stinson, Florian Busch, Joyce Pereira Alvarenga, and Matt Stata for collecting seeds used in this study, and Chandra Bellasio, Ingo Ensminger, Nicholas Levis, Matthew Osmond, and Arthur Weis for helpful discussions. This research was supported by Queen Elizabeth II/Charles Eckenwalder Scholarships to A.L., Australian Research Council Discovery Grant DP130102243 to M.L., and Natural Sciences and Engineering Research Council Discovery Grants RGPIN 2020 05925 and RGPIN 2017 06476 to T.L.S and R.F.S.

## Author contributions

A.L. and R.F.S. designed the research. M.L., T.L.S, and R.F.S. consulted on experimental setup and methods. M.L. generated the antibodies. A.L. conducted the research and wrote the manuscript with input from all other coauthors. Artificial Intelligence-generated content was not used in the preparation of any portion of this manuscript nor analysis of the results.

## Data availability

The data and code associated with this article are deposited on GitHub: https://github.com/aleungplants/c2-low-co2-acclimation.

