## Supplementary material for "Acclimation to enhanced photorespiratory conditions increases the effectiveness of C_2_ photosynthesis in C_3_-C_4_ intermediates": Methods S1

### Supplementary Methods

**Supplementary Methods S1. Growth chamber CO_2_ control and fertilization.**

Plants were propagated from cuttings (*Flaveria*, *Homolepis*) or seeds (*Tribulus*, *Steinchisma*) and grown in 9 L pots in a sandy-loam soil (Leung *et al.*, 2024). Pots were watered every other day and fertilized weekly with a 1:1 mix of Classic (20-20-20) and Acid (21-7-7) fertilizers (0.875 g L^-1^ each; Plant Products), supplemented with 0.025 g L^-1^ ferric ethylenediamine-N,N'-bis(2-hydroxyphenylacetic acid) (Fe-EDDHA) and 0.15 g L^-1^ each of MgSO_4_ and CaNO_3_.

Plants were grown at ambient CO_2_ (approximately 420 µmol mol^-1^) in a greenhouse for at least four weeks before transfer to growth CO_2_ concentrations of 200 or 800 µmol mol^-1^. Within the GC20 growth cabinets (BioChambers; <https://www.biochambers.com/>), 30 min dawn and dusk periods were imposed, during which temperature and PPFD were ramped between day and night settings. Growth CO_2_ was held within 20 µmol mol^-1^ of the set-point using WMA-4 infrared gas analyzers (PP Systems; <https://ppsystems.com/>), which actuated a solenoid valve connected to CO_2_ cylinders to raise CO_2_, or fans drawing chamber air through soda lime scrubbers to lower CO_2_ (McCann & Sage, 2022). The flowrate of the CO_2_ gas was adjusted to ensure recovery of CO_2_ concentrations to the setpoint within 15 min of opening the doors of the growth chambers. Plants were acclimated to the growth CO_2_ concentration for at least 10 days before measurements began.

**Supplementary Methods S2. Gas exchange measurement sequences.**

Gas exchange was measured with two LI-6400XT systems fitted with 6400-02B LED light sources (Li-Cor Biosciences; <https://www.licor.com/>) at a leaf temperature of 30 °C, following previous protocols (Leung *et al.*, 2024). Responses of gas exchange to intercellular CO_2_ concentration (*C_i_*) were measured at 1500 µmol m^-2^ s^-1^ PPFD at ambient CO_2_ concentration (*C_a_*) setpoints of 400, 300, 200, 90, 80, 70, 60, 50, 400, 400, 500, 600, 800, 1200, 1500, and 400 µmol mol^-1^, applied in that order. After each change in *C_a_*, gas exchange was allowed to stabilize for 90 s before matching sample and reference gas analyzer cells and allowed to stabilize for another 90 s before logging a data point. Each data point averaged the last 30 s of data.

The CO_2_ compensation point in the absence of day respiration (*C_i_**) and day respiration (*R_d_*) were determined by the Laisk method (Laisk, 1977; Brooks & Farquhar, 1985). After each *A*/*C_i_* response was measured at 1500 µmol m^-2^ s^-1^ PPFD, *A*/*C_i_* responses were measured at *C_a_* set-points of 400, 90, 80, 70, 60, 50, and 400 µmol mol^-1^ at each of four PPFDs (400, 250, 150, and 100 µmol m^-2^ s^-1^), applied in that order. Gas exchange was allowed to stabilize for 10 min after switching to 400 µmol m^-2^ s^-1^ PPFD and 5 min after each switch to other PPFDs. Following Walker & Ort (2015), the *y*-intercepts of these responses were regressed against their initial slopes; the *C_i_* at the common intersection equals *C_i_**, and the negative of the corresponding *A* equals *R_d_*. For C_3_ and proto-Kranz species all five irradiances were used; for sub-C_2_ species the four lowest were used, and for C_2_ species the three lowest, because photorespiratory refixation causes the higher-irradiance responses to diverge (Fig. S2).

**Supplementary Methods S3. Enzyme assay procedure.**

Leaves were sampled from plants in the growth chambers 2 to 4 h into the photoperiod, plunged into liquid N_2_, and stored at -80 °C until they were assayed. Total leaf-level enzyme activities were measured at 30 °C on a diode array spectrophotometer (Hewlett-Packard 8452A) by coupling NADPH or NADH oxidation or reduction to absorbance at 340 nm (Adachi *et al.*, 2023; Leung *et al.*, 2024).

Extracts were made at pH 8.0 in a previously described buffer (Adachi *et al.*, 2023) with the addition of 2 mM MnCl_2_, 0.08 mM pyridoxal phosphate, and 1% (v/v) protease inhibitor cocktail (Sigma P9599) and assayed within 15 min of extraction. PEPC, NADP-ME, Asp-AT, and Ala-AT were assayed as previously described (Adachi *et al.*, 2023), except that Asp-AT was assayed at pH 8.0 (Ashton *et al.*, 1990).

NAD-ME was assayed at pH 7.2 in 25 mM 4-(2-hydroxyethyl)-1-piperazineethanesulfonic acid (HEPES), 0.2 mM ethylenediaminetetraacetic acid (EDTA), 5 mM dithiothreitol, 0.1 mM coenzyme A, 1 unit mL^-1^ malate dehydrogenase, 5 mM *L*-malate, 2.5 mM MnCl_2_, 0.025 mM NADH, and 2.5 mM NAD, with the reaction initiated by 0.3 mM coenzyme A and 6 mM MnCl_2_ (Hatch *et al.*, 1982; Muhaidat *et al.*, 2011).

Chlorophyll a and b molar concentrations were determined in 80% (v/v, aqueous) acetone (Porra *et al.*, 1989). Chlorophyll a:b ratios are provided in Table S5 to facilitate conversion to chlorophyll amounts on a mass basis.

**Supplementary Methods S4. Immunolocalization of GLDP.**

Tissue was fixed in 1% (v/v) glutaraldehyde and 1% (w/v) paraformaldehyde in 0.1 M sodium cacodylate buffer, dehydrated in a graded ethanol series, and embedded in LR White, following previous procedures (Khoshravesh et al., 2017, 2020; Leung et al., 2024). Leaf segments of approximately 1 to 2 mm^2^ were sampled from the middle of the leaf, between the midrib and the margin.

For immunolabelling, 90 nm cross-sections were collected on formvar-coated nickel grids. Sections were incubated in 50 mM glycine for 15 min, 0.5% (w/v) bovine serum albumin (BSA) for 15 min, and 1:100 anti-GLDP P-protein primary antibody (Khoshravesh et al., 2017, 2020; Leung et al., 2024) in 0.5% BSA for 2 h. Sections were washed three times for 5 min in tris(hydroxymethyl)aminomethane-buffered saline (TBS; 12.5 mM Tris-HCl, 37.5 mM NaCl, pH 8.0), labelled with 1:20 anti-rabbit IgG secondary antibody conjugated to 18 nm gold particles (Jackson ImmunoResearch; <https://www.jacksonimmuno.com/>) in TBS for 1 h, and washed three more times for 5 min in TBS. All reagents were buffered in TBS except glycine, which was in deionized water. Sections were imaged at 80 kV on an HT7700 transmission electron microscope (Hitachi; <https://www.hitachi-hightech.com/>).

**Supplementary Methods S5. Quantification of ultrastructure and GLDP immunolabelling.**

For each sampled leaf, ultrastructure and immunolabelling parameters were quantified using ImageJ v1.53 (Schneider et al., 2012) from planar images of five BS and five M cells, and the mean of the five cells gave the value for the biological replicate, following previous procedures (Wang *et al.*, 2017; Khoshravesh *et al.*, 2020; Leung *et al.*, 2024).

Chloroplast, mitochondrion, and cell areas were traced, and organelle coverage was calculated as total planar chloroplast or mitochondrial area per cell area. Light micrographs were traced to determine the fraction of leaf chlorenchyma area occupied by BS and M tissues, where the BS fraction equals BS tissue area divided by the sum of BS and M tissue area.

For GLDP abundance, gold particles were counted on mitochondrial profiles in five cells each of BS and M tissue per replicate, from at least three mitochondria per M cell and at least five mitochondria per BS cell (typically 8 to 15 mitochondria in C_2_ species). Labelling density was calculated as the number of gold particles per planar mitochondrial area, and the BS allocation of leaf GLDP (*f*_GLDP_) was extrapolated as a function of labelling density, mitochondrial coverage, and tissue fraction of total chlorenchyma area (Khoshravesh *et al.*, 2020; Leung *et al.*, 2024).

BS vacuole shape was quantified from the same cross-sections as the perimeter-to-area ratio, circularity (4π area / perimeter^2^), and solidity (area / convex area) of the central vacuole. Chloroplast-mitochondrion and mitochondrion-chloroplast nearest neighbour distances were quantified using the custom ImageJ procedures described in Supplementary Methods S6.

**Supplementary Methods S6.** **Determination of chloroplast-mitochondrion and mitochondrion-chloroplast distances. See Fig. S6 for a schematic demonstrating the quantification of edge-to-edge distances.**

Transmission electron micrographs corresponding to each of five bundle sheath (BS) cells were each saved in a folder. The folder was opened as a stack in ImageJ.

Chloroplasts and mitochondria were traced and saved as Regions of Interest in ImageJ. Specific colors for each organelle type enabled systematic quantification of positioning using an ImageJ macro.

For each chloroplast, the distance between its edge and all mitochondria within each image slice was calculated. To capture all neighbouring mitochondria, each image had been taken at a sufficiently low magnification. The nearest neighbour distance was extracted for each chloroplast, and the mean of all chloroplasts in the centripetal half of the BS cell was used. To define the centripetal half, the BS cell was first halved by drawing a line within the cell walls radially from the vein. The centripetal and centrifugal halves were defined by a line at the halfway point of and perpendicular to the radial line. The mean of all five cells was used as the value for the biological replicate.

Two types of nearest neighbour distances were calculated: (i) distances between organelle centroids, where the centroid is the mean position of all points in the shape; and (ii) the shortest distances between organelle edges.

The above procedure was repeated for mitochondrion-chloroplast distances, which was implemented in ImageJ macro to facilitate quantification in a reproducible manner (downloadable from *Data Availability*).

**Supplementary Methods S7.** Photosynthesis model description.

A steady-state model of C_2_ photosynthesis was used to analyze the *A*/*C*_i_ responses measured at a PPFD of 1500 µmol m^-2^ s^-1^ (von Caemmerer, 1989, 2013). This model can be used to examine a continuum of C_3_ to C_2_ gas exchange responses. The model partitions leaf-level *A* into that of BS and M tissues (*A*_s_ and *A*_m_, respectively), such that

$$\begin{aligned} {A=A}_{s}+A_{m}, \#\left( 1 \right) \end{aligned}$$

$$\begin{aligned} {\mathrm{where} A}_{s}=V_{cs}-{0.5V}_{os}-R_{ds}, A_{m}=V_{cm}-{0.5V}_{om}-R_{dm},\#\left( 2,3 \right) \end{aligned}$$

and *V*_cs_, *V*_os_, and *R*_ds_ refer to the BS rubisco carboxylation rate, rubisco oxygenation rate, and day respiration, respectively, while *V*_cm_, *V*_om_, and *R*_dm_ are the corresponding values for the M tissue.

The glycine shuttle rate was defined as a fraction, α, of photorespiratory glycine produced in the M tissues that is decarboxylated in the BS tissues (0.5*V_o_*_m_):

$${glycine shuttle rate= 0.5\alpha V}_{om}$$

The CO_2_ leakage rate out of the BS tissues into M tissues (*L*) had two equivalent representations: (i) a balance between CO_2_ released from the C_2_ glycine shuttle with BS CO_2_ assimilation rate,

$$\begin{aligned} {L={0.5\alpha V}_{om}-A}_{s}, \#\left( 4 \right) \end{aligned}$$

and (ii) an Ohm’s-law formulation based on the CO_2_ gradient between BS and M:

$$\begin{aligned} {L=g}_{bs}\left( C_{s}-C_{m} \right),\#\left( 5 \right) \end{aligned}$$

where *g*_bs_ is the BS conductance, and *C*_s_ and *C*_m_ are the BS and M CO_2_ concentrations, respectively.

The M CO_2_ concentration was calculated from the *C*_i_ and a mesophyll conductance (*g*_m_):

$$\begin{aligned} C_{m}=C_{i}-\frac{A}{g_{m}}\#\left( 6 \right) \end{aligned}$$

Rubisco carboxylation rate in each tissue followed a Michaelis-Menten formulation, assuming CO_2_ concentration is the limiting substrate (Farquhar *et al.*, 1980).

$$\begin{aligned} {V_{cm}=V}_{m\max}\frac{C_{m}}{C_{m}{+K}_{c}\left( 1+\frac{O_{m}}{K_{o}} \right)}, {V_{cs}=V}_{s\max}\frac{C_{s}}{C_{s}{+K}_{c}\left( 1+\frac{O_{s}}{K_{o}} \right)},\#\left( 7,8 \right) \end{aligned}$$

where *K_c_* and *K_o_* are the Michaelis-Menten constants for CO_2_ and O_2_, respectively, and *O_m_* and *O_s_* are the O_2_ concentrations in the M and BS tissues, respectively.

Rubisco oxygenation rates in the M and BS tissues are

$$\begin{aligned} {V_{om}=2V}_{cm}\left( \frac{0.5}{S_{c/o}} \right)\left( \frac{O_{m}}{C_{m}} \right), {V_{os}=2V}_{cs}\left( \frac{0.5}{S_{c/o}} \right)\left( \frac{O_{s}}{C_{s}} \right),\#\left( 9,10 \right) \end{aligned}$$

where *S_c/o_* is the specificity factor of rubisco.

The M O_2_ concentration was assumed to be 210,000 µmol mol^-1^, and the BS O_2_ concentration was scaled to *A_s_* and assumed to diffuse out of the BS tissues at a rate proportional to *g*_bs_ (corrected with the relative diffusivities of O_2_ and CO_2_ with the 0.047 coefficient).

$$\begin{aligned} {O_{s}=O}_{m}+\frac{A_{s}}{0.047g_{bs}}\#\left( 11 \right) \end{aligned}$$

*V_m_*_max_ and *V_s_*_max_ were partitioned from the total maximal rubisco carboxylase activity (*V_c_*_max_) using the allocation of rubisco in the BS tissues (*f*_rubisco_):

$$\begin{aligned} V_{m\max}=\left( 1-f_{\mathrm{rubisco}} \right)V_{c\max}, V_{s\max}=f_{\mathrm{rubisco}}V_{c\max}\#\left( 12 \right) \end{aligned}$$

The leaf-level day respiration (*R*_d_) was partitioned among M, BS, and heterotrophic tissues using a fraction of day respiration in the heterotrophic tissues (*f*_Rdh_) and in the BS tissues (*f*_Rd_):

$$\begin{aligned} R_{dh}=f_{\mathrm{Rdh}}R_{d}, R_{dm}=\left( 1-f_{\mathrm{Rd}} \right)\left( R_{d}-R_{dh} \right), R_{ds}=f_{\mathrm{Rd}}\left( R_{d}-R_{dh} \right)\#\left( 13-15 \right) \end{aligned}$$

All intermediate variables were solved recursively by symbolic substitution using `sympy` in Python (Meurer *et al.*, 2017). Specifically, equations (13–15), along with the expressions for rubisco carboxylation and oxygenation (7–10), were substituted into the BS carbon balance (4) and Ohm’s-law expression (5) to solve for the bundle sheath CO_2_ concentration (*C*_s_). The positive solution of the quadratic equation was selected. The resulting *C*_s_ was substituted into the BS rubisco carboxylation and oxygenation rates (*V_c_*_s_, *V_o_*_s_) and BS O_2_ concentration (11), which in turn allowed evaluation of *A_s_*, *V_o_*_m_, and *A_m_*. Finally, all expressions were substituted into the leaf-level assimilation equation (1) to generate the net photosynthesis function, *A*(*C_m_*, *V_c_*_max_, *g_bs_*).

#### Model assumptions

Multiple assumptions were made to empirically parameterize the model using data collected using TEM. The allocation of GLDP to BS tissues was assumed to be the α, implying GDC operation in the M at its maximal turnover rate (Bellasio, 2017). The allocation of planar mitochondria area to the BS tissues was assumed to equal *f*_Rd_, assuming uniform respiration rates between mitochondria. The allocation of chloroplasts to the BS tissues was assumed to determine *f*_rubisco_, because the rubisco immunolabelling density in M and BS chloroplasts does not differ substantially between C_3_ and C_3_-C_4_ intermediate species (Bauwe, 1984; Ueno, 2011; Stata, 2023). The rubisco specificity factor and Michaelis-Menten kinetics were assumed to be those of a spinach rubisco, set as to an *in vitro* liquid-phase value of 76 mol mol^-1^ at 30°C (Jordan & Ogren, 1984) and converted to a gas-phase equivalent of 2015 mol mol^-1^ using CO_2_ and O_2_ solubilities (0.0334 and 0.00126 mol L^-1^ bar^-1^, respectively) (von Caemmerer, 2000). A value of *f_Rd_*_h_ = 0.1 was assigned arbitrarily to account for respiration in heterotrophic tissues such as the leaf vasculature (Tcherkez *et al.*, 2017). The mesophyll conductance (*g_m_*) was set to 0.6 mol m^-2^ s^-1^ bar^-1^, a realistic value at 30 °C (von Caemmerer & Evans, 2015). In the absence of actual rubisco kinetics and carbon isotope discrimination data, assigning arbitrary but biologically plausible values was considered appropriate because the primary objective was to evaluate relative CO_2_ acclimation effects.

To evaluate the consistency of modelled parameters under various assumed values of *g_m_* and *f*_Rdh_, the model was fitted across all combinations of *g_m_* = {0.3, 0.6, 0.9} mol m^-2^ s^-1^ bar^-1^ and *f*_Rdh_ = {0.0, 0.1, 0.2} (9 parameter combinations in total). While absolute values of modelled parameters varied with *g_m_* and *f*_Rdh_, the direction and relative magnitude of CO_2_ treatment effects were consistent across all parameter combinations (Fig. S7).

#### Model fitting

Model parameters *V_c_*_max_ and *g_bs_* were estimated by fitting the C_2_ photosynthesis model to *A*/*C*_i_ responses, using a Bayesian framework implemented in `emcee` in Python (Foreman-Mackey *et al.*, 2013). The model allowed for fitting of non-C_2_ *A*/*C_i_* responses by varying the empirically determined *f*_GLDP_, *f*_rubisco_, and *f_Rd_*_s_. A Bayesian approach was preferred to constrain fitted values to a realistic range. The model was fitted to points measured at *C*_a_

≤ 400 µmol mol^-1^, because the von Caemmerer (1989) formulation did not include RuBP regeneration-limited processes. Uniform priors, that is, probability distributions that assign equal probability to all possible values within a specified range, were imposed over biologically plausible ranges of 25 ≤ *V_c_*_max_ ≤ 250 µmol m^-2^ s^-1^ and 0 < *g_bs_* ≤ 0.2 mol m^-2^ s^-1^. The posterior distribution was sampled for 2500 iterations, with the first 500 steps discarded as burn-in, and median posterior values were used. The leakage rate (*L*), glycine shuttle rate (0.5*αV_o_*_m_) and BS net CO_2_ assimilation rate (*A*_s_) at *C*_a_ = 400 µmol mol^-1^ were then calculated by substituting the parameter estimates (median of the Bayesian posterior samples) and the corresponding *C*_m_ into the model expressions. Example model fits are shown for individual plants in Fig. S9.
