## Supplementary material for "Acclimation to enhanced photorespiratory conditions increases the effectiveness of C_2_ photosynthesis in C_3_-C_4_ intermediates": Table S1

### Supplementary Tables and Figures

**Table S1**. Gas exchange parameters in C_3_, proto-Kranz (PK), sub-C_2_, and C_2_ species grown at 200 and 800 µmol mol^-1^ CO_2_.

| Species | *Γ* (µmol mol^-1^) ^a^ | | *C*_i_* (µmol mol^-1^) ^a^ | | Initial slope (mol m^-2^ s^-1^) ^a^ | | *R*_d_ (µmol m^-2^ s^-1^) ^b^ | |
| --- | --- | --- | --- | --- | --- | --- | --- | --- |
|  | 200 | 800 | 200 | 800 |  |  | 200 | 800 |
| C_3_ |  |  |  |  |  |  |  |  |
| *F. cronquistii* | 58.4 ± 0.6 | 61.5 ± 2.1 | 50.1 ± 1.7 | 49.3 ± 2.0 | 0.152 ± 0.019 | 0.146 ± 0.009 | 1.15 ± 0.12 | 1.76 ± 0.13 |
| *H. glutinosa* | 53.4 ± 0.9 | 53.2 ± 0.8 | 46.4 ± 1.1 | 42.5 ± 1.8 | 0.089 ± 0.004 | 0.085 ± 0.005 | 0.57 ± 0.06 | 0.86 ± 0.17 |
| *F. pringlei* | **56.8 ± 0.9*** | **60.5 ± 1.2*** | 46.8 ± 1.1 | 41.6 ± 1.1 | 0.148 ± 0.018 | 0.114 ± 0.007 | **1.44 ± 0.06*** | **2.23 ± 0.35*** |
| Proto-Kranz |  |  |  |  |  |  |  |  |
| *T. forrestii* | 57.7 ± 2.4 | 61.5 ± 3.5 | 44.6 ± 0.5 | 49.8 ± 3.6 | 0.165 ± 0.010 | 0.168 ± 0.009 | 2.28 ± 0.41 | 1.86 ± 0.26 |
| *S. laxum* | 54.4 ± 1.2 | 53.3 ± 1.3 | 43.7 ± 1.7 | 43.2 ± 2.2 | 0.096 ± 0.014 | 0.102 ± 0.005 | 0.93 ± 0.10 | 1.09 ± 0.21 |
| Sub-C_2_ |  |  |  |  |  |  |  |  |
| *H. isocalycia* | **43.2 ± 0.9*** | **47.1 ± 1.6*** | 32.2 ± 0.6 | 36.1 ± 2.6 | 0.088 ± 0.006 | 0.106 ± 0.013 | **1.00 ± 0.06*** | **1.11 ± 0.15*** |
| *F. sonorensis* | 28.3 ± 2.2 | 34.3 ± 2.4 | **23.4 ± 3.7*** | **35.1 ± 1.8*** | 0.136 ± 0.019 | 0.128 ± 0.009 | 2.38 ± 0.12 | 1.94 ± 0.14 |
| *F. angustifolia* | **22.4 ± 3.3*** | **33.9 ± 1.5*** | **20.0 ± 2.7*** | **35.4 ± 4.7*** | 0.130 ± 0.013 | 0.112 ± 0.010 | **2.04 ± 0.11*** | **2.37 ± 0.45*** |
| C_2_ |  |  |  |  |  |  |  |  |
| *T. cristatus* | **15.3 ± 1.3*** | **22.9 ± 3.0*** | **9.2 ± 2.7*** | **18.8 ± 3.1*** | 0.139 ± 0.016 | 0.127 ± 0.023 | **1.93 ± 0.21*** | **1.42 ± 0.21*** |
| *H. aturensis* | 16.0 ± 1.0 | 21.2 ± 3.5 | **13.0 ± 2.9*** | **24.9 ± 4.1*** | 0.114 ± 0.008 | 0.098 ± 0.007 | 1.31 ± 0.10 | 1.21 ± 0.12 |
| *S. hians* | 13.1 ± 0.8 | 16.4 ± 1.8 | 12.3 ± 3.0 | 15.8 ± 4.3 | 0.102 ± 0.006 | 0.099 ± 0.002 | 1.50 ± 0.20 | 1.48 ± 0.27 |

Values represent means ± SE, based on *n* = 4 plants per CO_2_ treatment. The sub-headers ‘200’ and ‘800’ indicate the growth CO_2_ concentration in µmol mol^-1^. Asterisks (*) denote a statistically significant difference between 200 and 800 µmol mol^-1^ CO_2_ treatments (*p* < 0.05). Superscripts on column headers indicate test type: ^a^ One-tailed *t*-test for lower value in 200 µmol mol^-1^ CO_2_-grown plants; ^b^ two-tailed *t*-test.

Abbreviations: *Γ*, CO_2_ compensation point; *C*_i_*, CO_2_ compensation point in the absence of day respiration; *R*_d_, day respiration.

**Table S2.** Comparison of linear mixed-effects (LME) and phylogenetic linear mixed-effects (PLME) model contrasts for CO_2_ treatment effects within each photosynthetic phenotype.

| **Trait** | **Phot. type** | **Hypothesis** | **LME est.** | **LME SE** | **LME *t*** | **LME *p*** | **PLME est.** | **PLME SE** | **PLME *t*** | **PLME *p*** | **Agreement** |
| --- | --- | --- | --- | --- | --- | --- | --- | --- | --- | --- | --- |
| Gas exchange |  |  |  |  |  |  |  |  |  |  |  |
| *Γ* (µmol mol^-1^) | C_3_ | lower at 200 | -2.192 | 1.686 | -1.3 | 0.099 | -2.192 | 1.702 | -1.29 | 0.119 | Yes |
|  | PK | lower at 200 | -1.341 | 2.064 | -0.65 | 0.259 | -1.341 | 2.084 | -0.64 | 0.270 | Yes |
|  | sub-C_2_ | lower at 200 | **-7.111** | **1.686** | **-4.22** | **< 0.001** | **-7.111** | **1.702** | **-4.18** | **0.002** | Yes |
|  | C_2_ | lower at 200 | **-5.557** | **1.62** | **-3.43** | **< 0.001** | **-5.387** | **1.702** | **-3.17** | **0.008** | Yes |
| *C_i_** (µmol mol^-1^) | C_3_ | lower at 200 | 3.285 | 2.302 | 1.43 | 0.921 | 3.285 | 2.335 | 1.41 | 0.899 | Yes |
|  | PK | lower at 200 | -2.369 | 2.82 | -0.84 | 0.202 | -2.369 | 2.86 | -0.83 | 0.217 | Yes |
|  | sub-C_2_ | lower at 200 | **-10.357** | **2.302** | **-4.5** | **< 0.001** | **-10.357** | **2.335** | **-4.44** | **0.002** | Yes |
|  | C_2_ | lower at 200 | **-8.412** | **2.212** | **-3.8** | **< 0.001** | **-8.315** | **2.335** | **-3.56** | **0.005** | Yes |
| *Γ* at 250 µmol m^-2^ s^-1^ irradiance, *Γ*_250_ | C_3_ | lower at 200 | **-8.831** | **3.005** | **-2.94** | **0.002** | **-8.831** | **3.899** | **-2.27** | **0.029** | Yes |
| (µmol mol^-1^) | PK | lower at 200 | -0.774 | 3.68 | -0.21 | 0.417 | -0.774 | 4.775 | -0.16 | 0.438 | Yes |
|  | sub-C_2_ | lower at 200 | **-9.125** | **3.005** | **-3.04** | **0.002** | **-9.125** | **3.899** | **-2.34** | **0.026** | Yes |
|  | C_2_ | lower at 200 | -4.558 | 2.887 | -1.58 | 0.059 | -5.068 | 3.899 | -1.3 | 0.117 | Yes |
| *R_d_* (µmol m^-2^ s^-1^) | C_3_ | two-sided | **0.565** | **0.18** | **3.15** | **0.002** | **0.565** | **0.189** | **2.99** | **0.020** | Yes |
|  | PK | two-sided | -0.131 | 0.22 | -0.59 | 0.554 | -0.131 | 0.231 | -0.57 | 0.589 | Yes |
|  | sub-C_2_ | two-sided | -0.003 | 0.18 | -0.02 | 0.988 | -0.003 | 0.189 | -0.01 | 0.989 | Yes |
|  | C_2_ | two-sided | -0.235 | 0.173 | -1.36 | 0.178 | -0.212 | 0.189 | -1.12 | 0.299 | Yes |
| Ultrastructure |  |  |  |  |  |  |  |  |  |  |  |
| BS allocation of GLDP, *f*_GLDP_ (%) | C_3_ | greater at 200 | 0.548 | 5.382 | 0.1 | 0.460 | 0.548 | 4.847 | 0.11 | 0.457 | Yes |
|  | PK | greater at 200 | 6.652 | 6.592 | 1.01 | 0.158 | 6.652 | 5.936 | 1.12 | 0.150 | Yes |
|  | sub-C_2_ | greater at 200 | 4.678 | 5.382 | 0.87 | 0.194 | 4.678 | 4.847 | 0.97 | 0.183 | Yes |
|  | C_2_ | greater at 200 | **13.908** | **5.382** | **2.58** | **0.006** | **13.908** | **4.847** | **2.87** | **0.012** | Yes |
| BS colloidal gold particles per mito. area (µm^-2^) | C_3_ | greater at 200 | **12.418** | **6.735** | **1.84** | **0.035** | **12.418** | **4.451** | **2.79** | **0.013** | Yes |
|  | PK | greater at 200 | 7.28 | 8.249 | 0.88 | 0.190 | 7.28 | 5.452 | 1.34 | 0.112 | Yes |
|  | sub-C_2_ | greater at 200 | -1.057 | 6.735 | -0.16 | 0.562 | -1.057 | 4.451 | -0.24 | 0.590 | Yes |
|  | C_2_ | greater at 200 | 1.596 | 6.735 | 0.24 | 0.407 | 1.596 | 4.451 | 0.36 | 0.365 | Yes |
| M colloidal gold particles per mito. area (µm^-2^) | C_3_ | lower at 200 | 5.644 | 7.101 | 0.79 | 0.785 | 5.644 | 4.915 | 1.15 | 0.856 | Yes |
|  | PK | lower at 200 | 4.818 | 8.697 | 0.55 | 0.709 | 4.818 | 6.02 | 0.8 | 0.775 | Yes |
|  | sub-C_2_ | lower at 200 | -4.699 | 7.101 | -0.66 | 0.255 | -4.699 | 4.915 | -0.96 | 0.185 | Yes |
|  | C_2_ | lower at 200 | -4.937 | 7.101 | -0.7 | 0.245 | -4.937 | 4.915 | -1 | 0.174 | Yes |
| BS chloroplast size (µm^2^) | C_3_ | greater at 200 | -2.308 | 13.351 | -0.17 | 0.568 | -2.308 | 11.771 | -0.2 | 0.575 | Yes |
|  | PK | greater at 200 | 1.642 | 16.351 | 0.1 | 0.460 | 1.642 | 14.416 | 0.11 | 0.456 | Yes |
|  | sub-C_2_ | greater at 200 | -9.82 | 13.351 | -0.74 | 0.768 | -9.82 | 11.771 | -0.83 | 0.784 | Yes |
|  | C_2_ | greater at 200 | 5.939 | 13.351 | 0.44 | 0.329 | 5.939 | 11.771 | 0.5 | 0.315 | Yes |
| BS chloroplast number (cell profile^-1^) | C_3_ | greater at 200 | 0.033 | 1.407 | 0.02 | 0.491 | 0.033 | 1.453 | 0.02 | 0.491 | Yes |
|  | PK | greater at 200 | 1.025 | 1.723 | 0.59 | 0.277 | 1.025 | 1.779 | 0.58 | 0.291 | Yes |
|  | sub-C_2_ | greater at 200 | -1.579 | 1.407 | -1.12 | 0.867 | -1.579 | 1.453 | -1.09 | 0.844 | Yes |
|  | C_2_ | greater at 200 | -1.133 | 1.407 | -0.81 | 0.788 | -1.133 | 1.453 | -0.78 | 0.770 | Yes |
| BS allocation of chloro. area (%) | C_3_ | greater at 200 | -0.07 | 1.687 | -0.04 | 0.517 | -0.07 | 1.587 | -0.04 | 0.517 | Yes |
|  | PK | greater at 200 | 1.239 | 2.066 | 0.6 | 0.275 | 1.239 | 1.944 | 0.64 | 0.272 | Yes |
|  | sub-C_2_ | greater at 200 | -1.215 | 1.687 | -0.72 | 0.763 | -1.215 | 1.587 | -0.77 | 0.766 | Yes |
|  | C_2_ | greater at 200 | 2.468 | 1.687 | 1.46 | 0.074 | 2.468 | 1.587 | 1.55 | 0.082 | Yes |
| BS mitochondria size (µm^2^) | C_3_ | greater at 200 | -0.435 | 1.278 | -0.34 | 0.633 | -0.435 | 0.894 | -0.49 | 0.679 | Yes |
|  | PK | greater at 200 | 1.482 | 1.565 | 0.95 | 0.173 | 1.482 | 1.094 | 1.35 | 0.109 | Yes |
|  | sub-C_2_ | greater at 200 | -1.404 | 1.278 | -1.1 | 0.862 | -1.404 | 0.894 | -1.57 | 0.920 | Yes |
|  | C_2_ | greater at 200 | 1.185 | 1.278 | 0.93 | 0.178 | 1.185 | 0.894 | 1.33 | 0.113 | Yes |
| BS mitochondria number (cell profile^-1^) | C_3_ | greater at 200 | -1.25 | 2.59 | -0.48 | 0.685 | -1.25 | 1.253 | -1 | 0.824 | Yes |
|  | PK | greater at 200 | 3.9 | 3.173 | 1.23 | 0.111 | **3.9** | **1.535** | **2.54** | **0.019** | No |
|  | sub-C_2_ | greater at 200 | -1.958 | 2.59 | -0.76 | 0.774 | -1.958 | 1.253 | -1.56 | 0.919 | Yes |
|  | C_2_ | greater at 200 | 3.633 | 2.59 | 1.4 | 0.082 | **3.633** | **1.253** | **2.9** | **0.012** | No |
| BS allocation of mito. area (%) | C_3_ | greater at 200 | -0.342 | 4.579 | -0.07 | 0.530 | -0.342 | 3.522 | -0.1 | 0.537 | Yes |
|  | PK | greater at 200 | 2.354 | 5.609 | 0.42 | 0.338 | 2.354 | 4.313 | 0.55 | 0.301 | Yes |
|  | sub-C_2_ | greater at 200 | -0.074 | 4.579 | -0.02 | 0.506 | -0.074 | 3.522 | -0.02 | 0.508 | Yes |
|  | C_2_ | greater at 200 | 4.7 | 4.579 | 0.79 | 0.215 | 3.627 | 3.522 | 1.03 | 0.169 | Yes |
| BS vacuole perimeter:area (unitless) | C_3_ | lower at 200 | 0.009 | 0.056 | 0.16 | 0.562 | 0.009 | 0.073 | 0.12 | 0.546 | Yes |
|  | PK | lower at 200 | 0.035 | 0.069 | 0.51 | 0.695 | 0.035 | 0.089 | 0.4 | 0.648 | Yes |
|  | sub-C_2_ | lower at 200 | -0.003 | 0.056 | -0.05 | 0.479 | -0.003 | 0.073 | -0.04 | 0.484 | Yes |
|  | C_2_ | lower at 200 | -0.012 | 0.056 | -0.21 | 0.416 | -0.012 | 0.073 | -0.16 | 0.437 | Yes |
| BS vacuole circularity (unitless) | C_3_ | greater at 200 | -0.01 | 0.05 | -0.21 | 0.582 | -0.01 | 0.057 | -0.18 | 0.570 | Yes |
|  | PK | greater at 200 | -0.008 | 0.061 | -0.13 | 0.550 | -0.008 | 0.07 | -0.11 | 0.542 | Yes |
|  | sub-C_2_ | greater at 200 | 0.054 | 0.05 | 1.09 | 0.141 | 0.054 | 0.057 | 0.94 | 0.188 | Yes |
|  | C_2_ | greater at 200 | **0.131** | **0.05** | **2.62** | **0.005** | **0.131** | **0.057** | **2.28** | **0.028** | Yes |
| BS chloro.-mito. nearest neighbour edge dist. (µm) | C_3_ | lower at 200 | -0.168 | 0.27 | -0.62 | 0.267 | -0.168 | 0.267 | -0.63 | 0.275 | Yes |
|  | PK | lower at 200 | -0.092 | 0.33 | -0.28 | 0.391 | -0.092 | 0.327 | -0.28 | 0.394 | Yes |
|  | sub-C_2_ | lower at 200 | **-0.459** | **0.27** | **-1.7** | **0.046** | -0.459 | 0.267 | -1.72 | 0.065 | No |
|  | C_2_ | lower at 200 | **-0.509** | **0.27** | **-1.89** | **0.032** | **-0.509** | **0.267** | **-1.9** | **0.049** | Yes |
| BS mito.-chloro. nearest neighbour edge dist. (µm) | C_3_ | lower at 200 | 0.027 | 0.25 | 0.11 | 0.544 | 0.027 | 0.112 | 0.25 | 0.593 | Yes |
|  | PK | lower at 200 | 0.345 | 0.306 | 1.13 | 0.868 | 0.345 | 0.137 | 2.52 | 0.980 | Yes |
|  | sub-C_2_ | lower at 200 | -0.069 | 0.25 | -0.28 | 0.392 | -0.069 | 0.112 | -0.62 | 0.279 | Yes |
|  | C_2_ | lower at 200 | -0.056 | 0.25 | -0.22 | 0.412 | -0.056 | 0.112 | -0.5 | 0.317 | Yes |
| Enzyme activities (µmol mg Chl^-1^ h^-1^) |  |  |  |  |  |  |  |  |  |  |  |
| Asp-AT | C_3_ | greater at 200 | 7.981 | 36.698 | 0.22 | 0.414 | 7.981 | 41.047 | 0.19 | 0.426 | Yes |
|  | PK | greater at 200 | 21.709 | 44.946 | 0.48 | 0.315 | 21.709 | 50.272 | 0.43 | 0.339 | Yes |
|  | sub-C_2_ | greater at 200 | **83.93** | **36.698** | **2.29** | **0.013** | **83.93** | **41.047** | **2.04** | **0.040** | Yes |
|  | C_2_ | greater at 200 | **66.963** | **36.698** | **1.82** | **0.036** | 66.963 | 41.047 | 1.63 | 0.073 | No |
| Ala-AT | C_3_ | greater at 200 | 2.515 | 55.038 | 0.05 | 0.482 | 2.515 | 74.365 | 0.03 | 0.487 | Yes |
|  | PK | greater at 200 | 23.923 | 67.407 | 0.35 | 0.362 | 23.923 | 91.079 | 0.26 | 0.400 | Yes |
|  | sub-C_2_ | greater at 200 | 62.684 | 55.038 | 1.14 | 0.129 | 62.684 | 74.365 | 0.84 | 0.214 | Yes |
|  | C_2_ | greater at 200 | -28.48 | 55.038 | -0.52 | 0.697 | -28.48 | 74.365 | -0.38 | 0.643 | Yes |
| NAD-ME | C_3_ | greater at 200 | 20.827 | 46.101 | 0.45 | 0.326 | 20.827 | 57.278 | 0.36 | 0.363 | Yes |
|  | PK | greater at 200 | 41.784 | 56.462 | 0.74 | 0.231 | 41.784 | 70.151 | 0.6 | 0.285 | Yes |
|  | sub-C_2_ | greater at 200 | 6.414 | 46.101 | 0.14 | 0.445 | 6.414 | 57.278 | 0.11 | 0.457 | Yes |
|  | C_2_ | greater at 200 | 49.493 | 46.101 | 1.07 | 0.143 | 49.493 | 57.278 | 0.86 | 0.208 | Yes |
| NADP-ME | C_3_ | greater at 200 | -0.368 | 3.634 | -0.1 | 0.540 | -0.44 | 4.04 | -0.11 | 0.542 | Yes |
|  | PK | greater at 200 | **7.461** | **4.262** | **1.75** | **0.043** | 7.461 | 4.948 | 1.51 | 0.088 | No |
|  | sub-C_2_ | greater at 200 | -1.05 | 3.634 | -0.29 | 0.613 | -0.946 | 4.04 | -0.23 | 0.589 | Yes |
|  | C_2_ | greater at 200 | 3.559 | 4.556 | 0.78 | 0.219 | 5.7 | 4.04 | 1.41 | 0.101 | Yes |
| PEPC | C_3_ | greater at 200 | -9.793 | 21.955 | -0.45 | 0.671 | -10.374 | 21.948 | -0.47 | 0.675 | Yes |
|  | PK | greater at 200 | -13.773 | 25.744 | -0.53 | 0.703 | -13.773 | 26.881 | -0.51 | 0.688 | Yes |
|  | sub-C_2_ | greater at 200 | -12.719 | 21.955 | -0.58 | 0.718 | -11.14 | 21.948 | -0.51 | 0.686 | Yes |
|  | C_2_ | greater at 200 | 7.707 | 27.521 | 0.28 | 0.390 | 7.49 | 21.948 | 0.34 | 0.371 | Yes |
| Modelled parameters |  |  |  |  |  |  |  |  |  |  |  |
| *A_s_* (µmol m^-2^ s^-1^) | C_3_ | greater at 200 | 0.08 | 0.307 | 0.26 | 0.398 | 0.08 | 0.234 | 0.34 | 0.371 | Yes |
|  | PK | greater at 200 | 0.416 | 0.376 | 1.11 | 0.136 | 0.416 | 0.287 | 1.45 | 0.095 | Yes |
|  | sub-C_2_ | greater at 200 | 0.718 | **0.307** | **2.34** | **0.011** | **0.718** | **0.234** | **3.07** | **0.009** | Yes |
|  | C_2_ | greater at 200 | **1.532** | **0.295** | **5.19** | **< 0.001** | **1.502** | **0.234** | **6.42** | **< 0.001** | Yes |
| *C_s_* (µmol mol^-1^) | C_3_ | greater at 200 | -0.101 | 44.706 | 0 | 0.501 | -0.101 | 92.34 | 0 | 0.500 | Yes |
|  | PK | greater at 200 | 10.065 | 54.753 | 0.18 | 0.427 | 10.065 | 113.093 | 0.09 | 0.466 | Yes |
|  | sub-C_2_ | greater at 200 | **195.365** | **44.706** | **4.37** | **< 0.001** | **195.365** | **92.34** | **2.12** | **0.036** | Yes |
|  | C_2_ | greater at 200 | **204.34** | **42.952** | **4.76** | **< 0.001** | **185.654** | **92.34** | **2.01** | **0.042** | Yes |
| Glycine shuttle rate (µmol m^-2^ s^-1^) | C_3_ | greater at 200 | 0.061 | 0.387 | 0.16 | 0.438 | 0.061 | 0.442 | 0.14 | 0.447 | Yes |
|  | PK | greater at 200 | 0.636 | 0.474 | 1.34 | 0.092 | 0.636 | 0.542 | 1.17 | 0.139 | Yes |
|  | sub-C_2_ | greater at 200 | 0.659 | 0.387 | 1.7 | **0.047** | 0.659 | 0.442 | 1.49 | 0.090 | No |
|  | C_2_ | greater at 200 | **1.792** | **0.372** | **4.82** | **< 0.001** | **1.721** | **0.442** | **3.89** | **0.003** | Yes |
| *g*_bs_ (mmol m^-2^ s^-1^ bar^-1^) | C_3_ | lower at 200 | 1.958 | 7.923 | 0.25 | 0.597 | 1.958 | 4.206 | 0.47 | 0.672 | Yes |
|  | PK | lower at 200 | 5.866 | 9.703 | 0.6 | 0.726 | 5.866 | 5.151 | 1.14 | 0.854 | Yes |
|  | sub-C_2_ | lower at 200 | -**19.031** | **7.923** | **-2.4** | **0.009** | **-19.031** | **4.206** | **-4.52** | **0.001** | Yes |
|  | C_2_ | lower at 200 | -1.072 | 7.612 | -0.14 | 0.444 | -1.086 | 4.206 | -0.26 | 0.402 | Yes |
| *V_c_*_max_ (µmol m^-2^ s^-1^) | C_3_ | greater at 200 | 11.656 | 9.392 | 1.24 | 0.109 | 11.656 | 8.115 | 1.44 | 0.097 | Yes |
|  | PK | greater at 200 | 1.236 | 11.503 | 0.11 | 0.457 | 1.236 | 9.938 | 0.12 | 0.452 | Yes |
|  | sub-C_2_ | greater at 200 | 4.485 | 9.392 | 0.48 | 0.317 | 4.485 | 8.115 | 0.55 | 0.299 | Yes |
|  | C_2_ | greater at 200 | **20.683** | **9.024** | **2.29** | **0.012** | **19.276** | **8.115** | **2.38** | **0.025** | Yes |
| Leakage rate, *L* (µmol m^-2^ s^-1^) | C_3_ | lower at 200 | -0.019 | 0.168 | -0.11 | 0.455 | -0.019 | 0.315 | -0.06 | 0.477 | Yes |
|  | PK | lower at 200 | 0.22 | 0.206 | 1.07 | 0.855 | 0.22 | 0.386 | 0.57 | 0.707 | Yes |
|  | sub-C_2_ | lower at 200 | -0.059 | 0.168 | -0.35 | 0.363 | -0.059 | 0.315 | -0.19 | 0.428 | Yes |
|  | C_2_ | lower at 200 | 0.26 | 0.162 | 1.61 | 0.944 | 0.22 | 0.315 | 0.7 | 0.746 | Yes |

Each row reports the estimated contrast (200 vs. 800 µmol mol^-1^) for a given trait within a photosynthetic phenotype, with one-sided *p*-values where the direction of effect was predicted *a priori*. LME models included species as a random intercept with plant-level replication (*n* = 4 per species per CO_2_ treatment). PLME models incorporated a Brownian motion phylogenetic correlation structure. LME and PLME agreed in 106 of 112 comparisons (95%). Of the six disagreements, four reflect LME significance with PLME marginally non-significant (0.06 < *p* < 0.09), and two (BS mitochondrion number in proto-Kranz and C_2_) reflect PLME significance with LME non-significant. Bold *p*-values indicate *p* < 0.05. Abbreviations: est., estimate; SE, standard error; *Γ*, CO_2_ compensation point; *C_i_**, CO_2_ compensation point in the absence of day respiration; *Γ*_250_, CO_2_ compensation point measured at 250 µmol m^-2^ s^-1^ irradiance; BS, bundle sheath; M, mesophyll; GLDP, glycine decarboxylase P-protein; chloro., chloroplast; mito., mitochondrion; Asp-AT, aspartate aminotransferase; Ala-AT, alanine aminotransferase; NAD-ME, NAD-malic enzyme; NADP-ME, NADP-malic enzyme; PEPC, phosphoenolpyruvate carboxylase; *A_s_*, BS net CO_2_ assimilation rate; *C_s_*, BS chloroplastic CO_2_ concentration; *g*_bs_, BS conductance to CO_2_; *V_c_*_max_, maximum rubisco carboxylation rate; *L*, CO_2_ leakage rate from BS to M; PK, proto-Kranz; Phot. type, photosynthetic phenotype.

**Table S3.** Ultrastructural parameters in C_3_, proto-Kranz, sub-C_2_, and C_2_ species grown at 200 and 800 µmol mol^-1^ CO_2_.

| Species | BS allocation of  planar chloro. area (%) ^a^ | | BS allocation of  planar mito. area (%) ^a^ | | Chloro.-mito.  edge distance (µm) ^b^ | | BS allocation of GLDP (%) ^a^ | | BS colloidal gold  particle density (µm^-2^) ^a^ | |
| --- | --- | --- | --- | --- | --- | --- | --- | --- | --- | --- |
|  | 200 | 800 | 200 | 800 | 200 | 800 | 200 | 800 | 200 | 800 |
| C_3_ |  |  |  |  |  |  |  |  |  |  |
| *F. cronquistii* | 2.4 ± 0.9 | 1.8 ± 0.3 | 12.2 ± 2.4 | 8.5 ± 0.2 | 1.35 ± 0.70 | 1.11 ± 0.48 | 8.4 ± 1.3 | 6.0 ± 0.3 | 53 ± 7 | 39 ± 5 |
| *H. glutinosa* | 2.2 ± 0.1 | 2.9 ± 0.4 | 4.8 ± 0.9 | 6.1 ± 0.4 | 2.18 ± 0.73 | 1.79 ± 0.28 | 3.3 ± 0.2 | 3.4 ± 0.6 | 26 ± 9 | 8 ± 1 |
| *F. pringlei* | 1.1 ± 0.3 | 0.7 ± 0.1 | 5.4 ± 0.6 | 5.5 ± 2.2 | 1.48 ± 0.15 | 2.75 ± 0.53 | 3.2 ± 0.8 | 1.5 ± 0.8 | 20 ± 9 | 8 ± 2 |
| Proto-Kranz |  |  |  |  |  |  |  |  |  |  |
| *T. forrestii* | 11.1 ± 3.2 | 7.5 ± 1.3 | 36.0 ± 12.7 | 19.8 ± 5.6 | 0.51 ± 0.19 | 1.18 ± 0.57 | 45.3 ± 17.5 | 14.2 ± 3.5 | **26 ± 1*** | **13 ± 1*** |
| *S. laxum* | 7.9 ± 1.1 | 6.7 ± 1.8 | 25.3 ± 5.1 | 28.9 ± 8.7 | 0.44 ± 0.20 | 0.24 ± 0.04 | 33.8 ± 8.5 | 37.7 ± 11.6 | 36 ± 17 | 36 ± 20 |
| Sub-C_2_ |  |  |  |  |  |  |  |  |  |  |
| *H. isocalycia* | 9.1 ± 1.3 | 9.2 ± 1.6 | 29.0 ± 5.4 | 26.7 ± 4.6 | 0.68 ± 0.03 | 0.75 ± 0.29 | 44.3 ± 6.3 | 32.9 ± 6.1 | 34 ± 5 | 48 ± 19 |
| *F. sonorensis* | 11.8 ± 2.0 | 13.1 ± 3.8 | 32.6 ± 4.2 | 32.0 ± 6.1 | **0.74 ± 0.12*** | **1.26 ± 0.14*** | 80.4 ± 4.9 | 78.7 ± 6.5 | 24 ± 3 | 19 ± 3 |
| *F. angustifolia* | 12.5 ± 0.5 | 14.7 ± 3.5 | 45.7 ± 2.7 | 48.9 ± 5.5 | **1.11 ± 0.12*** | **1.46 ± 0.08*** | 86.8 ± 2.2 | 85.8 ± 4.3 | 22 ± 4 | 16 ± 5 |
| C_2_ |  |  |  |  |  |  |  |  |  |  |
| *T. cristatus* | 9.4 ± 3.2 | 9.3 ± 3.4 | 41.9 ± 8.8 | 38.1 ± 13.8 | 1.02 ± 0.08 | 1.26 ± 0.46 | 81.6 ± 5.0 | 65.4 ± 21.8 | 20 ± 2 | 20 ± 3 |
| *H. aturensis* | **23.9 ± 1.7*** | **16.6 ± 2.9*** | 51.0 ± 5.9 | 44.6 ± 4.5 | **0.55 ± 0.09*** | **1.12 ± 0.19*** | **89.2 ± 4.3*** | **78.7 ± 2.1*** | 40 ± 3 | 38 ± 9 |
| *S. hians* | 15.3 ± 1.7 | 12.0 ± 3.1 | 44.4 ± 5.3 | 35.2 ± 5.3 | 0.29 ± 0.09 | 0.38 ± 0.14 | **86.9 ± 2.4*** | **64.9 ± 5.0*** | 36 ± 3 | 32 ± 1 |

Values represent means ± SE, based on *n* = 4 plants per CO_2_ treatment. The sub-headers ‘200’ and ‘800’ indicate the growth CO_2_ concentration in µmol mol^-1^. Asterisks (*) denote a statistically significant difference between 200 and 800 µmol mol^-1^ CO_2_ treatments (*p* < 0.05). Superscripts on column headers indicate test type: ^a^ One-tailed *t*-test for greater values in 200 µmol mol^-1^ CO_2_-grown plants; ^b^ one-tailed *t*-test for lower values in 200 µmol mol^-1^ CO_2_-grown plants. Abbreviations: chloro., chloroplast; mito., mitochondria; BS, bundle sheath; GLDP, glycine decarboxylase P-protein.

**Table S4**. *In vitro* activities relative to chlorophyll concentration of aspartate aminotransferase (Asp-AT), alanine aminotransferase (Ala-AT), NAD-malic enzyme (NAD-ME), NADP-malic enzyme (NADP-ME), and phosphoenolpyruvate carboxylase (PEPC).

| Species | Ala-AT  (µmol mg Chl^-1^ h^-1^) | | Asp-AT  (µmol mg Chl^-1^ h^-1^) | | NAD-ME  (µmol mg Chl^-1^ h^-1^) | | NADP-ME  (µmol mg Chl^-1^ h^-1^) | | PEPC  (µmol mg Chl^-1^ h^-1^) | |
| --- | --- | --- | --- | --- | --- | --- | --- | --- | --- | --- |
|  | 200 | 800 | 200 | 800 | 200 | 800 | 200 | 800 | 200 | 800 |
| C_3_ |  |  |  |  |  |  |  |  |  |  |
| *F. cronquistii* | 62.3 ± 8.8 | 55.1 ± 13.3 | 36.2 ± 4.5 | 40.2 ± 5.9 | 62.3 ± 4.2 | 72.0 ± 9.1 | -0.5 ± 0.2 | -0.4 ± 0.7 | 18.7 ± 3.8 | 18.1 ± 4.1 |
| *H. glutinosa* | 50.0 ± 5.7 | 49.3 ± 5.3 | 21.3 ± 2.4 | 22.2 ± 4.1 | 22.5 ± 1.2 | 34.2 ± 3.7 | 3.2 ± 0.5 | 3.2 ± 1.0 | 13.5 ± 1.9 | 17.6 ± 2.3 |
| *F. pringlei* | 62.7 ± 5.8 | 81.4 ± 6.3 | 33.0 ± 3.7 | 29.0 ± 6.6 | 62.5 ± 14.3 | 46.5 ± 8.4 | 1.0 ± 0.4 | 1.4 ± 0.9 | 8.9 ± 1.8 | 16.4 ± 4.3 |
| Proto-Kranz |  |  |  |  |  |  |  |  |  |  |
| *T. forrestii* | 118.4 ± 41.7 | 109.8 ± 21.6 | 58.7 ± 12.3 | 59.7 ± 11.8 | 80.2 ± 21.9 | 63.6 ± 10.2 | 4.3 ± 0.7 | 3.9 ± 0.2 | 42.5 ± 10.6 | 30.1 ± 2.2 |
| *S. laxum* | 70.9 ± 12.8 | 67.6 ± 14.2 | 64.3 ± 10.3 | 52.4 ± 3.8 | 51.2 ± 3.7 | 47.0 ± 11.1 | **6.4 ± 0.9*** | **3.0 ± 1.1*** | 20.2 ± 1.9 | 39.5 ± 19.0 |
| Sub-C_2_ |  |  |  |  |  |  |  |  |  |  |
| *H. isocalycia* | 63.8 ± 7.3 | 73.5 ± 9.3 | 58.3 ± 3.6 | 71.5 ± 4.6 | 76.3 ± 17.5 | 93.5 ± 10.6 | 6.5 ± 0.6 | 8.2 ± 1.5 | 10.5 ± 1.7 | 21.1 ± 2.8 |
| *F. sonorensis* | 106.9 ± 15.3 | 82.1 ± 21.0 | **94.0 ± 10.3*** | **55.4 ± 15.9*** | 41.0 ± 5.9 | 26.3 ± 8.7 | 4.2 ± 1.1 | 3.3 ± 0.1 | 14.9 ± 2.5 | 14.2 ± 1.3 |
| *F. angustifolia* | 112.8 ± 19.4 | 81.1 ± 5.2 | 95.5 ± 16.3 | 58.2 ± 10.3 | 40.5 ± 13.8 | 33.2 ± 6.5 | 6.1 ± 2.0 | 6.1 ± 2.1 | 17.1 ± 4.5 | 15.5 ± 2.6 |
| C_2_ |  |  |  |  |  |  |  |  |  |  |
| *T. cristatus* | 201.1 ± 22.4 | 270.9 ± 19.5 | 205.4 ± 30.7 | 193.6 ± 10.2 | 84.4 ± 29.3 | 89.6 ± 8.5 | 8.5 ± 2.3 | 10.9 ± 0.3 | 62.0 ± 8.6 | 59.7 ± 9.5 |
| *H. aturensis* | 55.0 ± 3.8 | 41.5 ± 9.7 | 45.9 ± 3.2 | 36.9 ± 4.8 | 43.6 ± 9.3 | 51.5 ± 9.8 | 8.6 ± 2.7 | 5.6 ± 0.2 | 14.9 ± 3.2 | 16.2 ± 1.4 |
| *S. hians* | **123.4 ± 10.7*** | **88.4 ± 6.1*** | **141.3 ± 9.8*** | **112.0 ± 6.5*** | **102.1 ± 9.7*** | **52.0 ± 10.5*** | 7.0 ± 1.8 | 3.4 ± 1.1 | 27.4 ± 0.5 | 22.8 ± 5.9 |

Values represent means ± SE, based on *n* = 4 plants per CO_2_ treatment. The sub-headers ‘200’ and ‘800’ indicate the growth CO_2_ concentration in µmol mol^-1^. Asterisks (*) denote a statistically significant difference between 200 and 800 µmol mol^-1^ CO_2_ treatments, using one-tailed *t*-tests for greater activities in 200 µmol mol^-1^ CO_2_-grown plants (*p* < 0.05). Activities expressed on a leaf area basis and chlorophyll a/b ratios are reported in Table S5.

**Table S5.** *In vitro* activities relative to leaf area of aspartate aminotransferase (Asp-AT), alanine aminotransferase (Ala-AT), NAD-malic enzyme (NAD-ME), NADP-malic enzyme (NADP-ME), and phosphoenolpyruvate carboxylase (PEPC).

| Species | Ala-AT  (µmol m^-2^ s^-1^) | | Asp-AT  (µmol m^-2^ s^-1^) | | NAD-ME  (µmol m^-2^ s^-1^) | | NADP-ME  (µmol m^-2^ s^-1^) | | PEPC  (µmol m^-2^ s^-1^) | | Chlorophyll a/b  (mol mol^-1^) | |
| --- | --- | --- | --- | --- | --- | --- | --- | --- | --- | --- | --- | --- |
|  | 200 | 800 | 200 | 800 | 200 | 800 | 200 | 800 | 200 | 800 | 200 | 800 |
| C_3_ |  |  |  |  |  |  |  |  |  |  |  |  |
| *F. cronquistii* | 33.0 ± 2.0 | 31.8 ± 9.1 | 20.0 ± 2.8 | 22.5 ± 2.6 | 34.4 ± 4.1 | 41.6 ± 6.8 | -0.3 ± 0.1 | -0.3 ± 0.4 | 10.4 ± 2.3 | 10.1 ± 2.1 | 3.39 ± 0.07 | 3.36 ± 0.04 |
| *H. glutinosa* | 25.4 ± 2.1 | 22.5 ± 3.1 | 11.2 ± 2.0 | 9.8 ± 1.3 | 11.6 ± 0.9 | 15.1 ± 0.4 | 1.6 ± 0.2 | 1.6 ± 0.7 | 6.8 ± 0.7 | 8.1 ± 1.4 | 3.22 ± 0.06 | 3.11 ± 0.20 |
| *F. pringlei* | 31.3 ± 1.6 | 50.5 ± 7.1 | 16.3 ± 0.4 | 17.0 ± 3.1 | 33.2 ± 9.0 | 27.5 ± 3.4 | 0.5 ± 0.2 | 0.9 ± 0.5 | 4.9 ± 1.0 | 9.3 ± 1.9 | 2.96 ± 0.10 | 3.00 ± 0.06 |
| Proto-Kranz |  |  |  |  |  |  |  |  |  |  |  |  |
| *T. forrestii* | 65.4 ± 16.2 | 70.4 ± 11.9 | 34.3 ± 2.3 | 37.6 ± 4.9 | 46.7 ± 9.4 | 40.5 ± 3.6 | 2.6 ± 0.3 | 2.6 ± 0.2 | 25.4 ± 4.9 | 19.6 ± 0.7 | 3.74 ± 0.19 | 3.64 ± 0.12 |
| *S. laxum* | 22.0 ± 3.8 | 26.3 ± 4.2 | 19.8 ± 2.3 | 20.6 ± 0.7 | 15.9 ± 0.4 | 18.5 ± 4.4 | 2.0 ± 0.3 | 1.2 ± 0.4 | 6.4 ± 1.0 | 14.7 ± 6.0 | 3.34 ± 0.07 | 3.19 ± 0.26 |
| Sub-C_2_ |  |  |  |  |  |  |  |  |  |  |  |  |
| *H. isocalycia* | 26.4 ± 2.2 | 28.0 ± 6.2 | 24.5 ± 2.3 | 27.0 ± 4.5 | 31.3 ± 6.6 | 34.6 ± 5.0 | 2.7 ± 0.2 | 3.0 ± 0.7 | 4.5 ± 0.8 | 7.9 ± 1.4 | 3.35 ± 0.10 | 3.38 ± 0.16 |
| *F. sonorensis* | **72.0 ± 10.6*** | **44.5 ± 7.7*** | **63.0 ± 6.1*** | **30.0 ± 6.4*** | **27.5 ± 3.9*** | **13.9 ± 3.1*** | 2.9 ± 0.9 | 1.9 ± 0.2 | 9.9 ± 1.5 | 8.1 ± 1.0 | 3.00 ± 0.04 | 2.95 ± 0.09 |
| *F. angustifolia* | 72.2 ± 13.4 | 50.9 ± 5.6 | **61.2 ± 10.3*** | **35.4 ± 4.7*** | 26.2 ± 8.6 | 20.0 ± 2.4 | 3.5 ± 0.9 | 3.6 ± 1.2 | 9.9 ± 2.2 | 9.3 ± 1.3 | 3.02 ± 0.07 | 2.99 ± 0.11 |
| C_2_ |  |  |  |  |  |  |  |  |  |  |  |  |
| *T. cristatus* | 103.2 ± 12.2 | 130.2 ± 9.1 | 103.7 ± 13.6 | 92.8 ± 3.3 | 41.1 ± 11.9 | 43.1 ± 4.4 | 4.2 ± 1.3 | 5.2 ± 0.2 | 29.3 ± 1.6 | 28.4 ± 4.0 | 3.05 ± 0.13 | 3.18 ± 0.08 |
| *H. aturensis* | 29.1 ± 2.5 | 22.9 ± 4.4 | 24.4 ± 2.5 | 20.7 ± 2.6 | 23.7 ± 6.1 | 28.8 ± 5.3 | 5.1 ± 1.7 | 3.3 ± 0.1 | 8.8 ± 1.9 | 9.6 ± 0.9 | 2.98 ± 0.08 | 3.13 ± 0.17 |
| *S. hians* | 51.1 ± 6.5 | 44.4 ± 5.5 | 59.9 ± 10.3 | 56.6 ± 7.4 | 46.0 ± 11.3 | 24.6 ± 3.7 | 3.7 ± 0.6 | 1.9 ± 0.7 | 15.0 ± 1.7 | 11.9 ± 2.3 | 3.24 ± 0.03 | 3.27 ± 0.03 |

Values represent means ± SE, based on *n* = 4 plants per CO_2_ treatment. The sub-headers ‘200’ and ‘800’ indicate the growth CO_2_ concentration in µmol mol^-1^. Asterisks (*) denote a statistically significant difference between 200 and 800 µmol mol^-1^ CO_2_ treatments, using one-tailed *t*-tests for greater activities in 200 µmol mol^-1^ CO_2_-grown plants (*p* < 0.05).

**Table S6.** Modelled values of C_2_ photosynthesis parameters.

| Species | *V_c_*_max_  (µmol m^-2^ s^-1^) ^c^ | | *C_s_*  *(*µmol mol^-1^) | | *g_bs_*  (mmol m^-2^ s^-1^ bar^-1^) ^b^ | | Glycine shuttle rate  (µmol m^-2^ s^-1^) ^a^ | | *A_s_*  (µmol m^-2^ s^-1^) ^a^ | | *L*  (µmol m^-2^ s^-1^) ^b^ | |
| --- | --- | --- | --- | --- | --- | --- | --- | --- | --- | --- | --- | --- |
|  | 200 | 800 | 200 | 800 | 200 | 800 | 200 | 800 | 200 | 800 | 200 | 800 |
| C_3_ |  |  |  |  |  |  |  |  |  |  |  |  |
| *F. cronquistii* | 142 ± 15 | 136 ± 11 | 212 ± 13 | 207 ± 7 | 116 ± 1 | 112 ± 2 | **0.75 ± 0.08*** | **0.51 ± 0.04*** | **0.61 ± 0.07*** | **0.39 ± 0.05*** | 0.14 ± 0.04 | 0.12 ± 0.02 |
| *H. glutinosa* | 86 ± 4 | 78 ± 5 | 209 ± 5 | 203 ± 7 | 96 ± 6 | 84 ± 3 | 0.18 ± 0.01 | 0.16 ± 0.01 | 0.35 ± 0.02 | 0.39 ± 0.02 | -0.17 ± 0.01 | -0.22 ± 0.01 |
| *F. pringlei* | 127 ± 13 | 106 ± 7 | 239 ± 6 | 250 ± 10 | 95 ± 1 | 98 ± 1 | **0.24 ± 0.03*** | **0.09 ± 0.01*** | **0.26 ± 0.03*** | **0.08 ± 0.01*** | **-0.02 ± 0.01*** | **0.02 ± 0.00*** |
| Proto-Kranz |  |  |  |  |  |  |  |  |  |  |  |  |
| *T. forrestii* | 165 ± 9 | 153 ± 14 | 210 ± 12 | 195 ± 16 | 150 ± 9 | 126 ± 8 | **4.29 ± 0.22*** | **1.32 ± 0.15*** | **3.20 ± 0.32*** | **1.77 ± 0.12*** | 1.09 ± 0.21 | -0.45 ± 0.11 |
| *S. laxum* | 85 ± 10 | 97 ± 5 | 220 ± 10 | 214 ± 13 | 114 ± 22 | 95 ± 24 | 1.70 ± 0.21 | 2.22 ± 0.15 | 1.26 ± 0.12 | 1.16 ± 0.08 | **0.44 ± 0.10*** | **1.06 ± 0.13*** |
| Sub-C_2_ |  |  |  |  |  |  |  |  |  |  |  |  |
| *H. isocalycia* | 87 ± 6 | 93 ± 6 | 257 ± 12 | 226 ± 13 | 11 ± 2 | 29 ± 23 | **2.29 ± 0.19*** | **1.77 ± 0.12*** | 1.75 ± 0.10 | 1.59 ± 0.13 | 0.54 ± 0.11 | 0.19 ± 0.10 |
| *F. sonorensis* | 147 ± 18 | 141 ± 6 | 558 ± 114 | 302 ± 26 | 8 ± 4 | 26 ± 11 | 6.69 ± 0.83 | 6.13 ± 0.31 | 5.34 ± 0.64 | 4.67 ± 0.32 | 1.35 ± 0.38 | 1.46 ± 0.28 |
| *F. angustifolia* | 146 ± 15 | 133 ± 7 | **576 ± 109*** | **277 ± 9*** | **9 ± 6*** | **29 ± 4*** | 7.14 ± 0.73 | 6.24 ± 0.30 | **5.72 ± 0.61*** | **4.39 ± 0.30*** | 1.42 ± 0.35 | 1.85 ± 0.14 |
| C_2_ |  |  |  |  |  |  |  |  |  |  |  |  |
| *T. cristatus* | 165 ± 16 | 131 ± 18 | **1080 ± 32*** | **651 ± 74*** | 2 ± 0 | 2 ± 0 | **8.38 ± 0.89*** | **5.30 ± 0.79*** | **6.64 ± 0.69*** | **4.32 ± 0.65*** | 1.74 ± 0.20 | 0.98 ± 0.14 |
| *H. aturensis* | 125 ± 10 | 104 ± 11 | 256 ± 8 | 349 ± 30 | 3 ± 0 | 4 ± 2 | 5.48 ± 0.45 | 4.40 ± 0.48 | **5.33 ± 0.45*** | **3.96 ± 0.49*** | **0.15 ± 0.04*** | **0.44 ± 0.07*** |
| *S. hians* | 99 ± 13 | 108 ± 3 | **611 ± 30*** | **390 ± 11*** | 2 ± 0 | 3 ± 1 | 4.74 ± 0.64 | 4.02 ± 0.12 | 4.15 ± 0.59 | 3.42 ± 0.13 | 0.59 ± 0.06 | 0.60 ± 0.12 |

Values represent means ± SE, based on *n* = 4 plants per CO_2_ treatment. The sub-headers ‘200’ and ‘800’ indicate the growth CO_2_ concentration in µmol mol^-1^. Asterisks (*) denote a statistically significant difference between 200 and 800 µmol mol^-1^ CO_2_ treatments (*p* < 0.05). Superscripts on column headers indicate test type: ^a^ One-tailed *t*-test for greater value in 200 µmol mol^-1^ CO_2_-grown plants. ^b^ one-tailed *t*-test for lower value in 200 µmol mol^-1^ CO_2_-grown plants; ^c^ two-tailed *t*-test.

Abbreviations: *V_c_*_max_, maximal rubisco carboxylase activity; *g_bs_*, bundle sheath conductance to CO_2_, *A_bs_*; bundle sheath net CO_2_ assimilation rate; *C_s_*, bundle sheath chloroplastic CO_2_ concentration; *L*, rate of CO_2_ leakage from bundle sheath to mesophyll tissues.

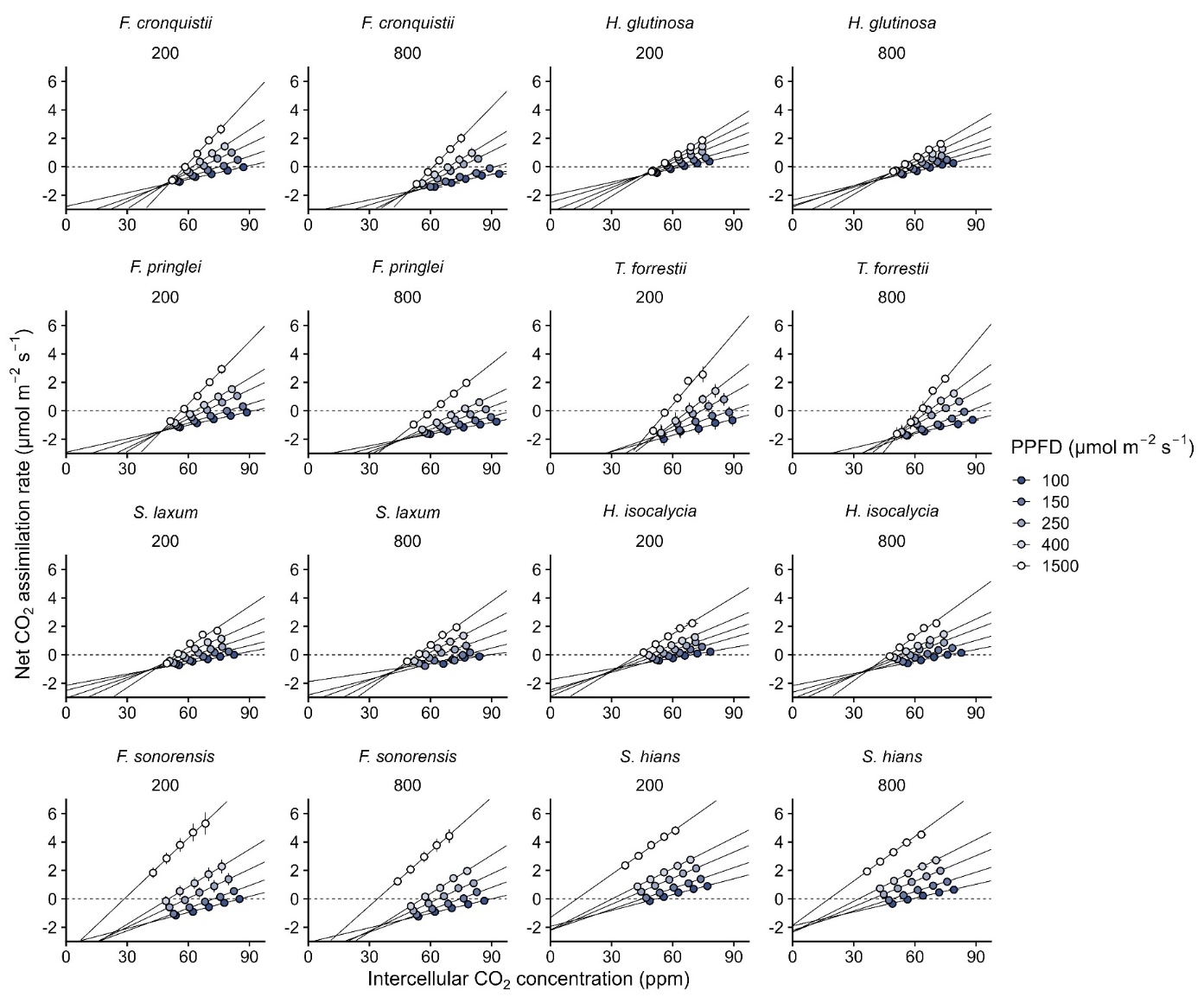

**Fig. S1.** Determination of the CO_2_ compensation point in the absence of day respiration (*C_i_**) and day respiration (*R_d_*) using the Laisk method. Mean ± SE (*n* = 4 plants) response of net CO_2_ assimilation rate (*A*) to intercellular CO_2_ concentration (*C_i_*) measured at various photosynthetic photon flux densities (PPFD; 100, 150, 250, 400, 1500 µmol m^-2^ s^-1^) in eleven species grown at 200 or 800 µmol mol^-1^ CO_2_ (indicated above each panel). The *C_i_* value at the common intersection point of the *A*/*C_i_* responses approximates *C_i_**.

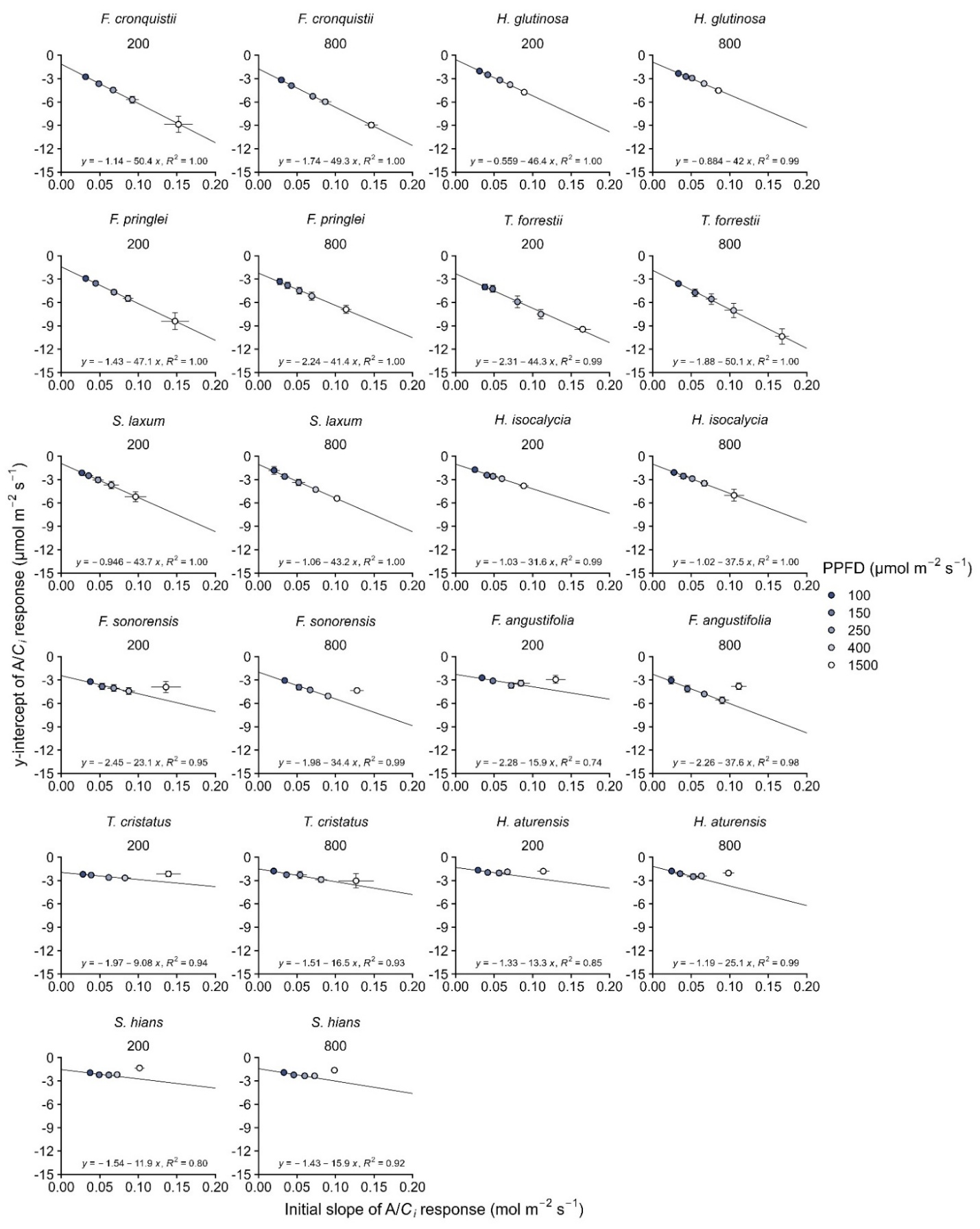

**Fig. S2.** Analysis of Laisk curves using the slope-intercept regression method. Linear regressions of the y-intercepts against the initial slopes of *A*/*C_i_* response curves measured at five photosynthetic photon flux densities (100, 150, 250, 400, and 1500 µmol m^-2^ s^-1^). To determine the intersection point, all five light intensities were used for C_3_ and proto-Kranz species, whereas only the four lowest intensities were used for sub-C_2_ species, and the three lowest intensities for C_2_ species. Data are shown for all eleven species grown at 200 and 800 µmol mol^-1^ CO_2_ (indicated above each panel). Each point represents the mean ± SE (*n* = 4 plants) of the slope and intercept parameters derived from the low-*C_i_* gas exchange data at a specific irradiance. The data presented in the main text were obtained from analysis of data collected from each individual plant, while the regressions here use the mean values as an example. The CO_2_ compensation point in the absence of day respiration (*C_i_**) is determined as the negative inverse of the slope of the linear regression, and day respiration (*R_d_*) is determined as the y-intercept of the regression (Walker & Ort, 2015). The high coefficient of determination (*R*^2^) across all panels confirms the consistency of the common intersection point estimate.

(see previous page for the figure)

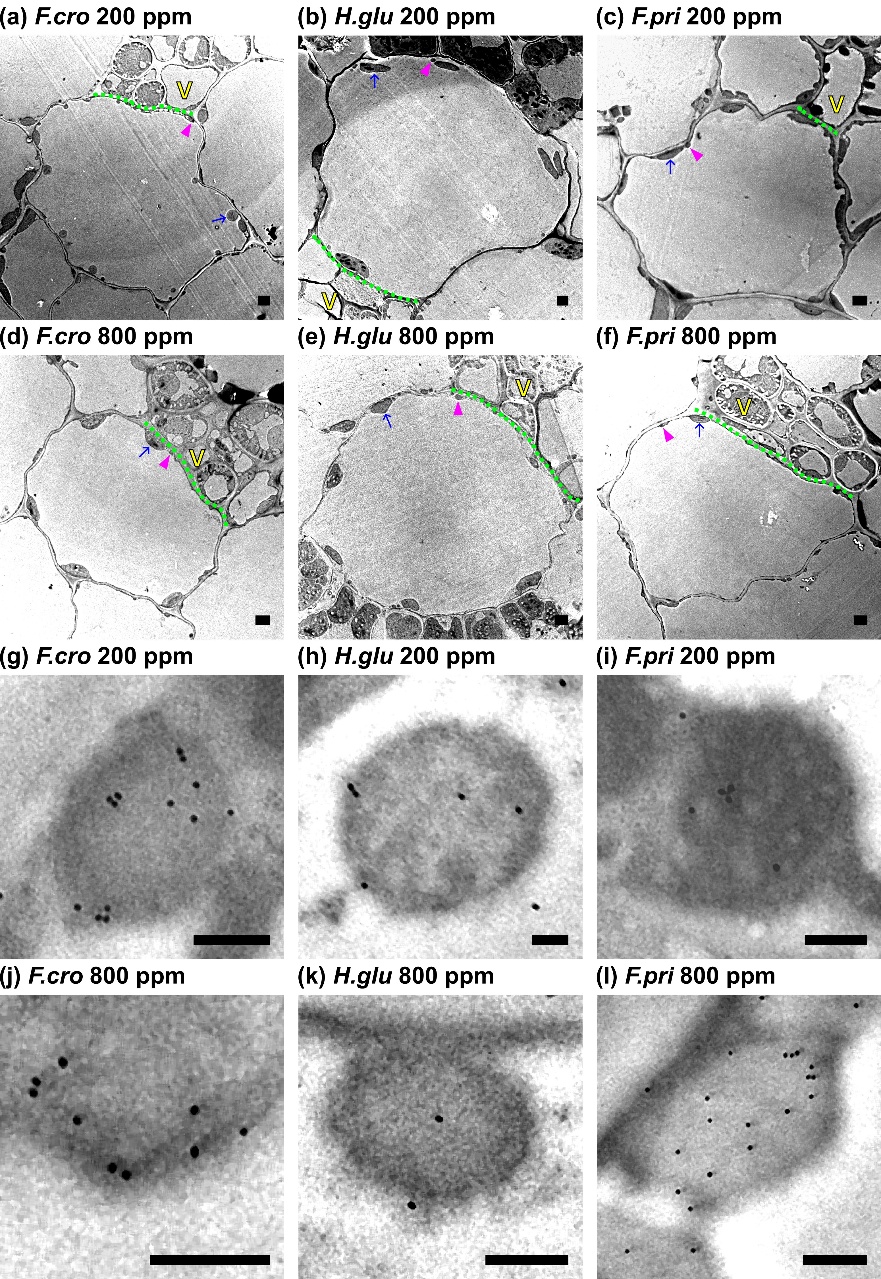

**Fig. S3.** Transmission electron microscopy and immunolocalization of GLDP in C_3_ species. Representative transmission electron micrographs of bundle sheath cells from three C_3_ species grown at 200 (**a–c**) and 800 (**d–f**) µmol mol^-1^ CO_2_. High-magnification views of bundle sheath mitochondria are shown for 200 (**g–i**) and 800 (**j–l**) µmol mol^-1^ CO_2_. Black dots are colloidal gold particles indicating the presence of the glycine decarboxylase P-protein (GLDP). Columns correspond to *Flaveria cronquistii* (**a,d,g,j**), *Homolepis glutinosa* (**b,e,h,k**), and *F. pringlei* (**c,f,i,l**). Arrows, chloroplasts; arrowheads, mitochondria; M, mestome sheath; V, vein. The green dotted line indicates the cell wall in contact with the vein in eudicots or the mestome sheath in grasses. Scale bars: 2 µm (whole-cell images), 0.5 µm (mitochondrial images).

(see previous page for the figure)

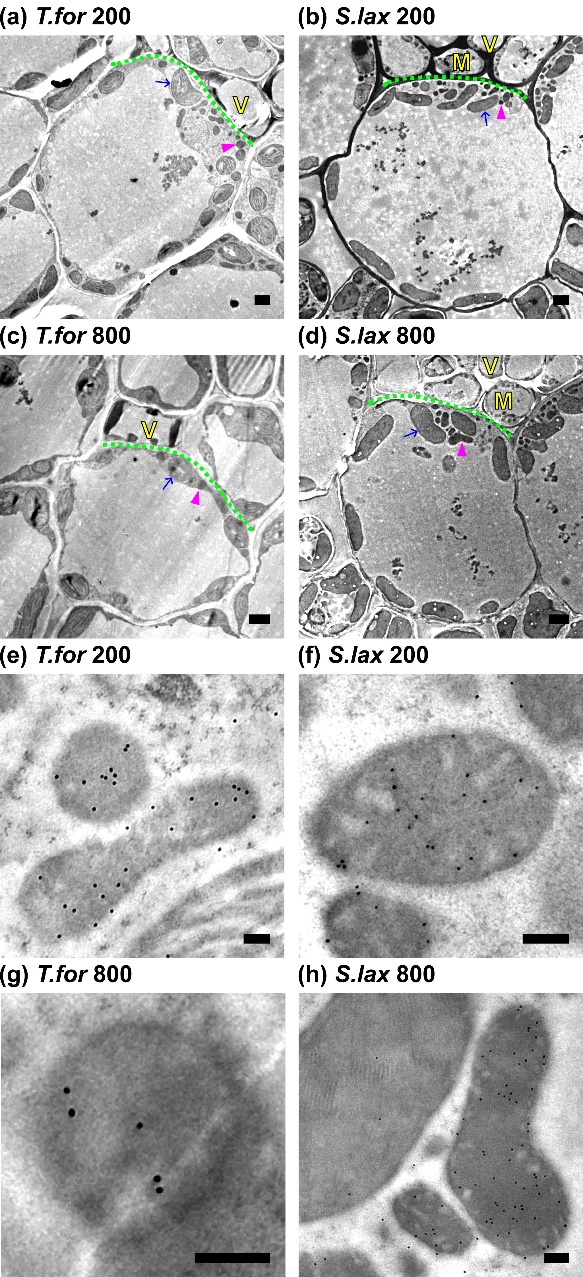

**Fig. S4.** Transmission electron microscopy and immunolocalization of GLDP in proto-Kranz species. Representative transmission electron micrographs of bundle sheath cells from two proto-Kranz species grown at 200 (**a,b**) and 800 (**c,d**) µmol mol^-1^ CO_2_. High-magnification views of bundle sheath mitochondria are shown for 200 (**e,f**) and 800 (**g,h**) µmol mol^-1^ CO_2_. Black dots are colloidal gold particles indicating the presence of the glycine decarboxylase P-protein (GLDP). Columns correspond to *Tribulus forrestii* (**a,c,e,g**) and Steinchisma *laxum* (**b,d,f,h**). Arrows, chloroplasts; arrowheads, mitochondria; M, mestome sheath; V, vein. The green dotted line indicates the cell wall in contact with the vein in eudicots or the mestome sheath in grasses. Scale bars: 2 µm (whole-cell images), 0.5 µm (mitochondrial images).

(see previous page for the figure)

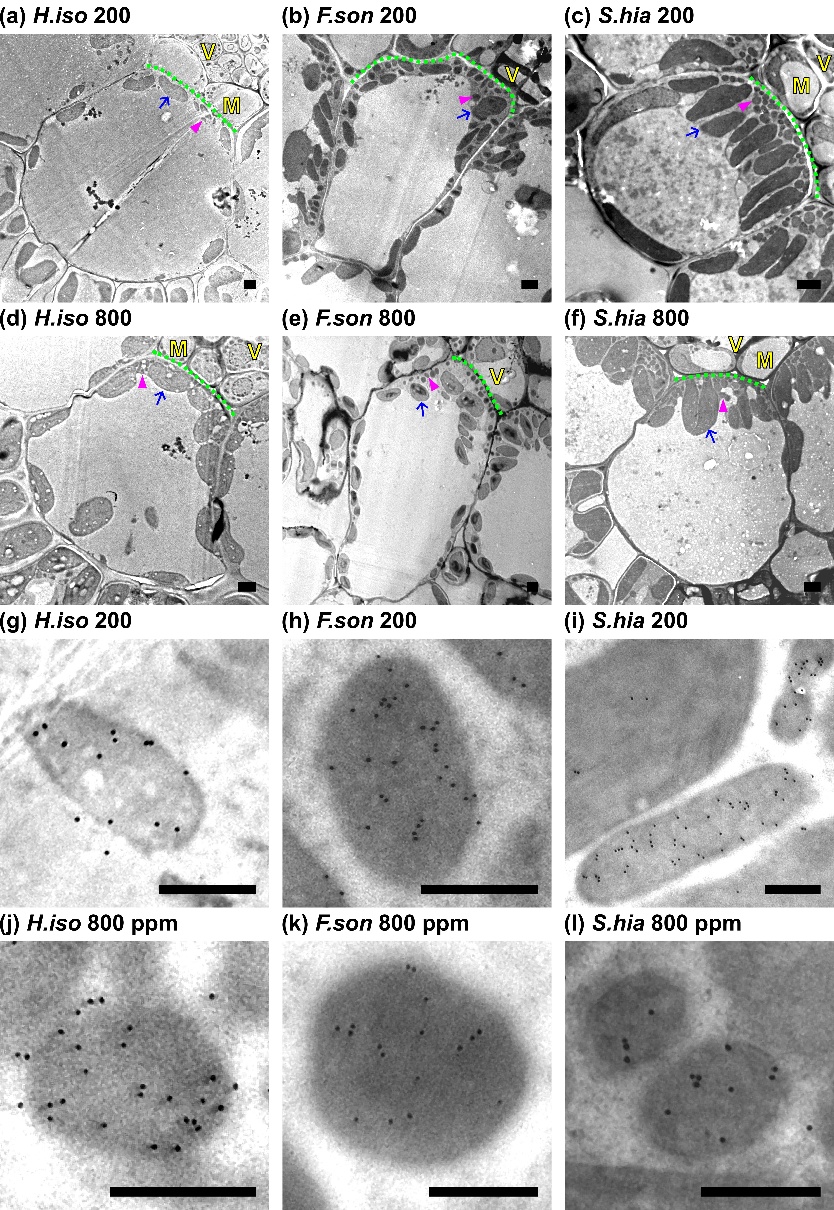

**Fig. S5.** Transmission electron microscopy and immunolocalization of GLDP in sub-C_2_ and C_2_ species. Representative transmission electron micrographs of bundle sheath cells from sub-C_2_ and C_2_ species not shown in Fig. 2, grown at 200 (**a–c**) and 800 (**d–f**) µmol mol^-1^ CO_2_. High-magnification views of bundle sheath mitochondria are shown for 200 (**g–i**) and 800 (**j–l**) µmol mol^-1^ CO_2_. Black dots are colloidal gold particles indicating the presence of the glycine decarboxylase P-protein (GLDP). Columns correspond to *Homolepis isocalycia* (sub-C_2_; **a,d,g,j**), *Flaveria sonorensis* (sub-C_2_; **b,e,h,k**), and *Steinchisma hians* (C_2_; **c,f,i,l**). Arrows, chloroplasts; arrowheads, mitochondria; M, mestome sheath; V, vein. The green dotted line indicates the cell wall in contact with the vein in eudicots or the mestome sheath in grasses. Scale bars: 2 µm (whole-cell images), 0.5 µm (mitochondrial images).

(see previous page for the figure)

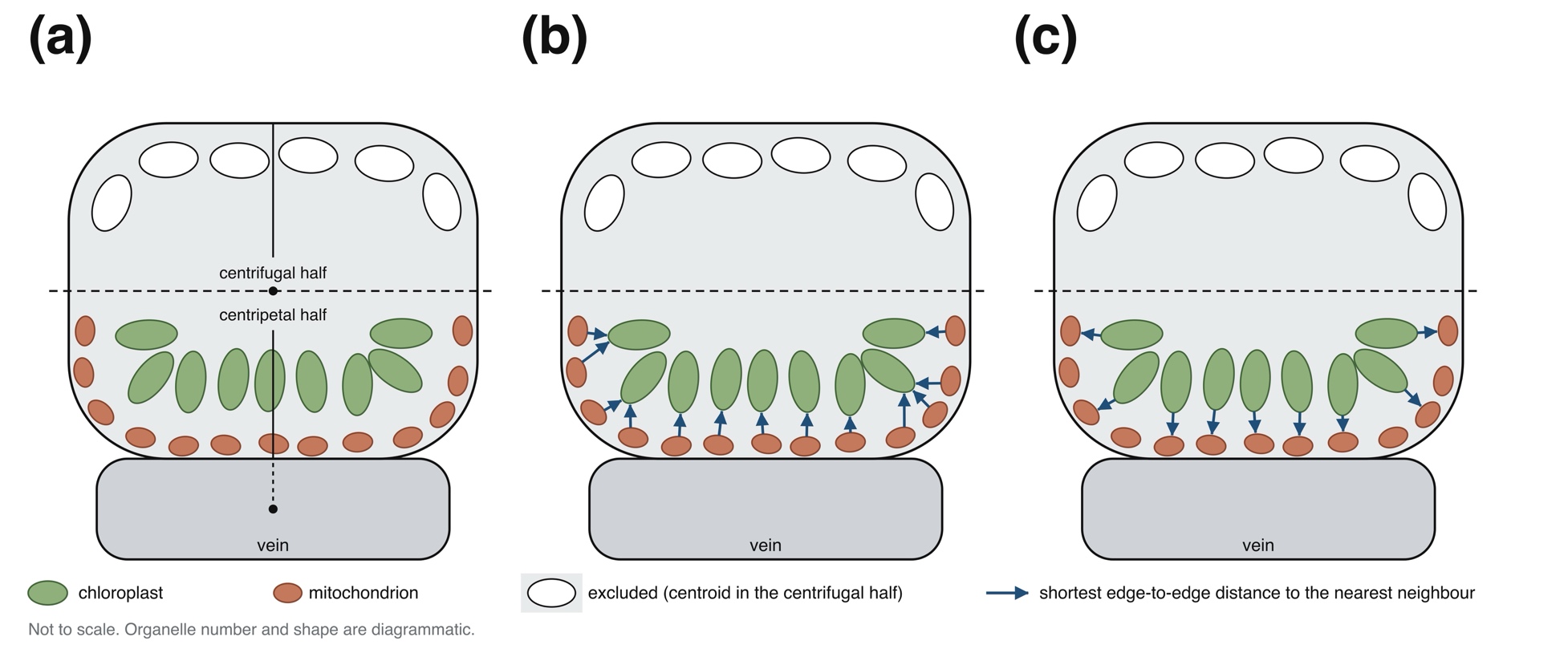

**Fig. S6.** Schematic of the quantification of edge-to-edge chloroplast-mitochondrion and mitochondrion-chloroplast distances in the centripetal half of a bundle sheath (BS) cell. In BS cells of sub-C_2_ and C_2_ species, mitochondria are positioned against the centripetal wall and chloroplasts are positioned over them towards the centrifugal side; mitochondria are rare in the centrifugal half. (a) A radial line is drawn from the centre of the vein through the cell, positioned so that it divides the cell profile into left and right halves of equal area, and the cell is divided at the midpoint of the segment of the radial line inside the cell walls by a perpendicular (dashed). Organelles whose centroid falls in the centrifugal half, drawn as open outlines, are excluded from the calculation. (b) Mitochondrion-chloroplast distance. For each mitochondrion in the centripetal half, the shortest distance between its traced edge and the traced edge of the nearest chloroplast (arrows), averaged over all mitochondria in the centripetal half. (c) Chloroplast-mitochondrion distance. For each chloroplast in the centripetal half, the shortest distance between its traced edge and the traced edge of the nearest mitochondrion (arrows), averaged over all chloroplasts in the centripetal half. The two directions are not reciprocal; a mitochondrion's nearest chloroplast need not have that mitochondrion as its own nearest neighbour, and one chloroplast can be the nearest neighbour of several mitochondria. Centroid-to-centroid distances were computed by the same procedure from organelle centroids in place of traced edges. Values are the mean of five cell profiles per biological replicate. The schematic is representative and not to scale; organelle number and shape are diagrammatic, and the mestome sheath that lies between the BS and the vein in *Steinchisma* and *Homolepis* is omitted. In the grasses the radial line is likewise drawn from the centre of the vein.

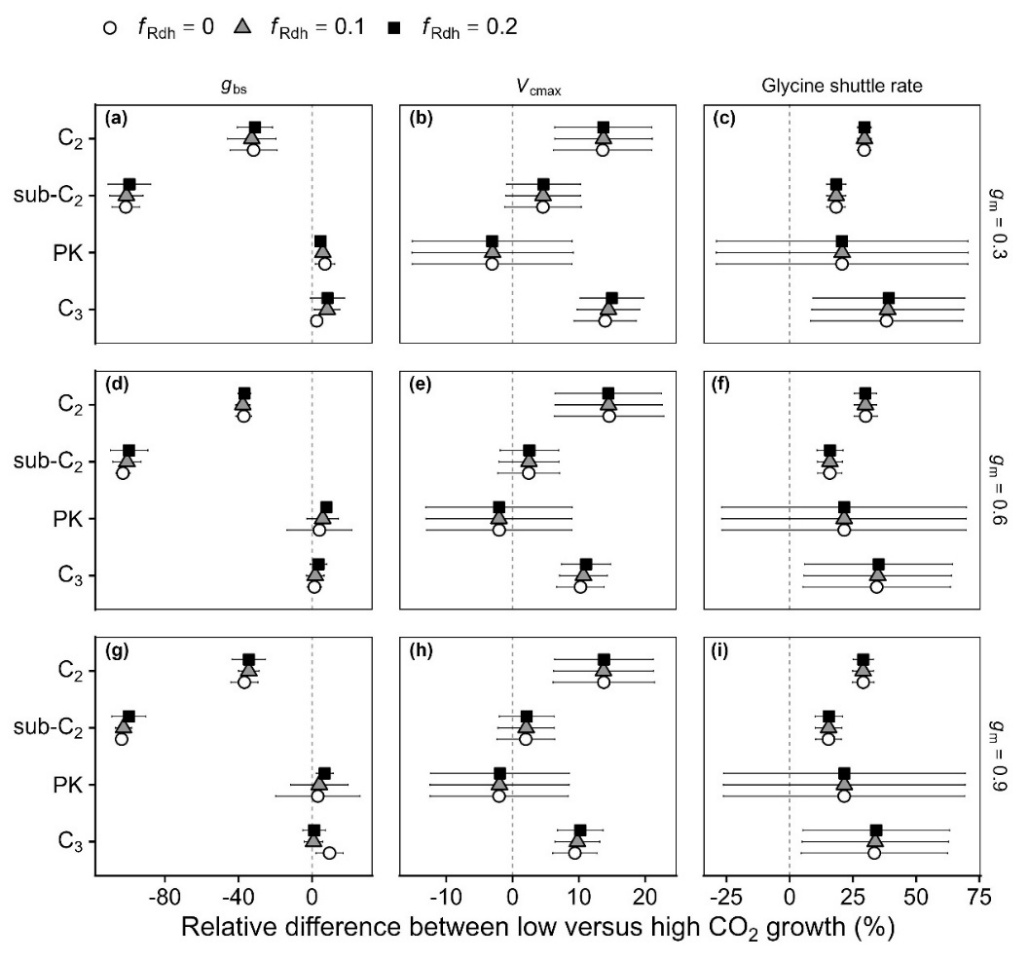

**Fig. S7.** Sensitivity of modelled CO_2_ treatment effects to assumed mesophyll conductance and heterotrophic tissue fraction of day respiration. The model was re-run across nine combinations of two assumed quantities: mesophyll conductance (*g_m_*, three values, one per row) and the fraction of day respiration originating from heterotrophic tissues (*f*_Rdh_, three values, one per symbol; Tcherkez *et al.*, 2017). Each panel plots the relative difference in a modelled parameter between plants grown at 200 and 800 µmol mol^-1^ CO_2_, calculated as (value at 200 - value at 800) / mean of both values × 100. Values to the left of the dashed line are lower at low growth CO_2_; values to the right are higher. Symbols within a photosynthetic phenotype show three assumed *f*_Rdh_ values (white circles, *f*_Rdh_ = 0; grey triangles*, f*_Rdh_ = 0.1; black squares*, f*_Rdh_ = 0.2). Rows: *g_m_* = 0.3 (a-c), 0.6 (d-f), and 0.9 (g-i) mol m^-2^ s^-1^ bar^-1^. Columns: bundle sheath conductance (*g*_bs_; a,d,g), maximum rubisco carboxylation rate (*V*_cmax_; b,e,h), and glycine shuttle rate (c,f,i). Within each panel, points and error bars are means ± SE across the species of each photosynthetic phenotype (C_3_, proto-Kranz (PK), sub-C_2_, C_2_).

The rows of figure panels representing different *g*_m_ values used to model photosynthetic parameters show qualitative similar results, indicating that the relative CO_2_ treatment differences are not affected by values of *g*_m_ that are 50% lower or higher than the value of 0.6 mol m^-2^ s^-1^ bar^-1^ used in this study.

Likewise, the symbols of differing colours and shapes each panel representing different *f*_Rdh_ values within are qualitatively similar within and between figure panels, indicating *f*_Rdh_ values that are 100% lower or higher than the value of 0.1 result in similar relative CO_2_ treatment differences.

Note the error bars are large when there are small absolute CO_2_ treatment differences (e.g., in the modelled glycine shuttle rate in C_3_ and proto-Kranz species).

(see previous page for the figure)

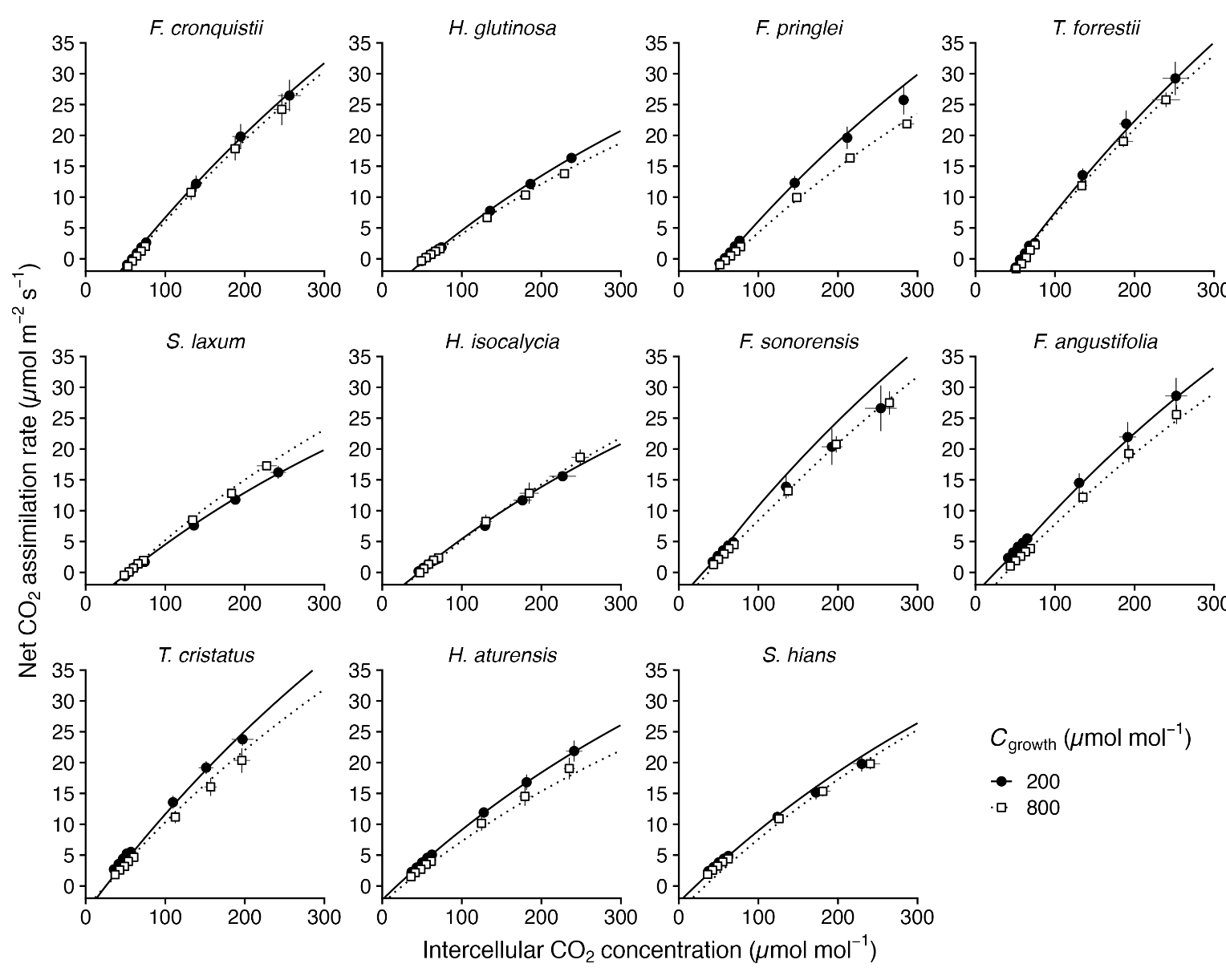

**Fig. S8.** Fitting steady-state C_2_ photosynthesis model to gas exchange data. Observed and modelled response of net CO_2_ assimilation rate (*A*) to intercellular CO_2_ concentration (*C_i_*) in eleven species grown at 200 µmol mol^-1^ (closed circles, solid lines) and 800 µmol mol^-1^ (open squares, dashed lines) growth CO_2_ concentrations (*C*_growth_). Points represent empirical gas exchange measurements (mean ± SE, n = 4 plants) measured at an irradiance of 1500 µmol m^-2^ s^-1^. Lines represent the mean of the model predictions fitted to data collected from each individual plant. The model was parameterized using the measured ultrastructural parameters (bundle sheath allocation of glycine decarboxylase P-protein and of chloroplast and mitochondria planar area) quantified in this study, while maximum rubisco carboxylation rate (*V_c_*_max_) and bundle sheath conductance (*g*_bs_) were estimated using a Bayesian framework using `emcee` in Python (Foreman-Mackey *et al.*, 2013).

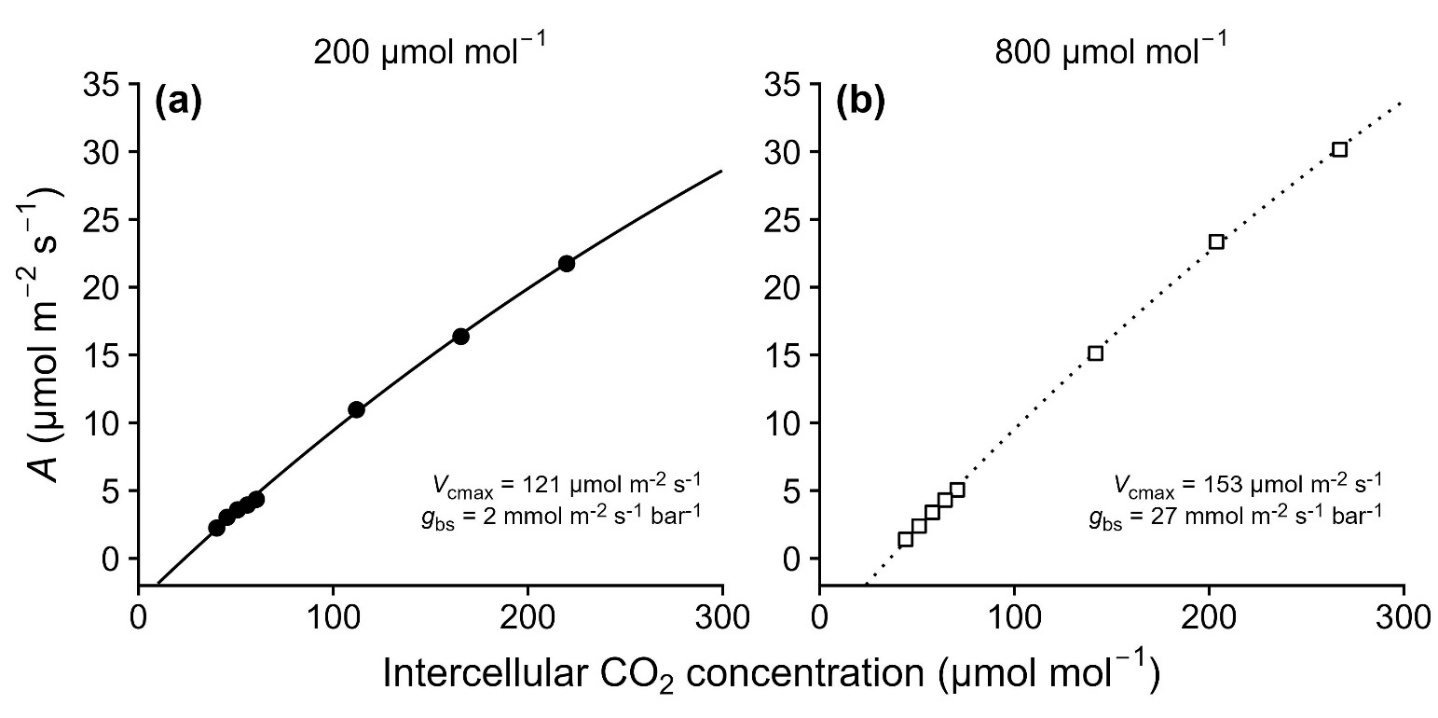

**Fig. S9.** Representative fits of the steady-state C_2_ photosynthesis model to gas exchange data from single *Flaveria angustifolia* (sub-C_2_) plants grown at each growth CO_2_ concentration (indicated above each panel). Observed net CO_2_ assimilation rate (*A*) against intercellular CO2 concentration (*C_i_*) measured at an irradiance of 1500 µmol m^-2^ s^-1^ (points), with the corresponding modelled response (line). (a), a plant grown at 200 µmol mol^-1^ CO_2_ (closed circles, solid line); (b), a plant grown at 800 µmol mol^-1^ CO_2_ (open squares, dotted line). Maximum rubisco carboxylation rate (*V_c_*_max_) and bundle sheath conductance to CO_2_ (*g*_bs_) estimated for each plant are given on each panel. Only points measured at a *C_a_* of 400 µmol mol^-1^ or below were used for fitting, following the rubisco carboxylation-limited formulation by von Caemmerer (1989).

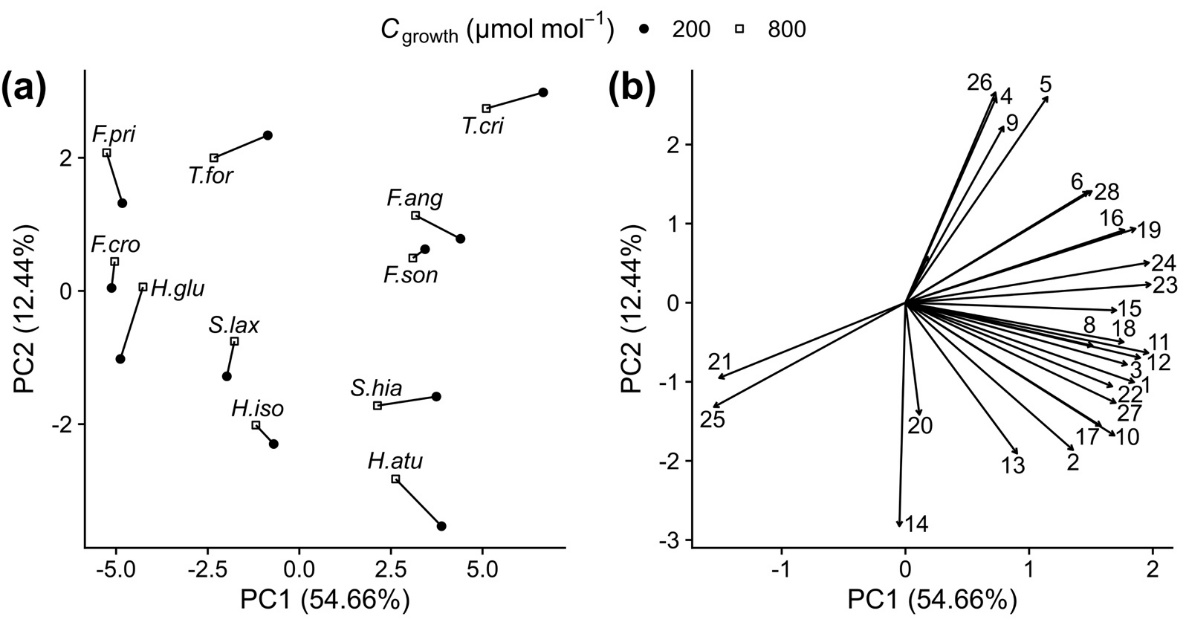

**Fig. S10.** Principal component analysis of all traits reported in Tables 1, 2, and 3, in contrast to subset shown in Fig. 7. Twenty-eight physiological, biochemical, and ultrastructural traits were measured in eleven species grown at two growth CO_2_ concentrations (*C*_growth_; closed circles, 200 µmol mol^-1^; open squares, 800 µmol mol^-1^). **(a)** Species scores, with points from the same species linked by lines representing the plastic response to low CO_2_. **(b)** Loading vectors, numbered as listed below. PC1 is oriented so that positive values correspond to more C_2_-like phenotypes. Acclimation to low CO_2_ shifted sub-C_2_ and C_2_ phenotypes towards more positive PC1 values in the same direction as evolution from C_3_ to C_2_ photosynthesis, matching the pattern in Fig. 7.

Loading vectors are numbered: 1, CO_2_ compensation point at 1500 µmol m^-2^ s^-1^ irradiance (*Γ*); 2, CO_2_ compensation point at 250 µmol m^-2^ s^-1^ irradiance (*Γ*_250_); 3, CO_2_ compensation point in the absence of day respiration (*C_i_**); 4, day respiration rate (*R_d_*); 5, alanine aminotransferase activity; 6, aspartate aminotransferase activity; 7, NAD-malic enzyme activity; 8, NADP-malic enzyme activity; 9, phosphoenolpyruvate carboxylase activity; 10, Bundle sheath (BS) allocation of chloroplast area; 11, BS allocation of glycine decarboxylase P-protein (*f*_GLDP_); 12, BS allocation of mitochondrial area; 13, BS chloroplast-to-mitochondrion nearest neighbour edge distance; 14, GLDP labelling density in BS mitochondria; 15, GLDP labelling density in mesophyll (M) mitochondria; 16, BS chloroplast number; 17, BS mitochondria number; 18, BS chloroplast size; 19, BS mitochondria size; 20, BS vacuole perimeter-to-area ratio; 21, BS vacuole circularity; 22, BS mitochondrion-to-chloroplast nearest neighbour edge distance; 23, BS net CO_2_ assimilation rate (*A_s_*); 24, M-to-BS glycine shuttle rate; 25, CO_2_ leakage rate from BS to M (*L*); 26, maximum rubisco carboxylation rate (V*_c_*_max_); 27, BS conductance to CO_2_ (*g*_bs_); 28, BS CO_2_ concentration (*C_s_*).
